# A human monoclonal antibody bivalently binding two different epitopes in streptococcal M protein protects against infection

**DOI:** 10.1101/2021.03.01.433494

**Authors:** Wael Bahnan, Lotta Happonen, Hamed Khakzad, Vibha Kumra Ahnlide, Therese de Neergaard, Sebastian Wrighton, Oscar André, Eleni Bratanis, Di Tang, Thomas Hellmark, Lars Björck, Oonagh Shannon, Lars Malmström, Johan Malmström, Pontus Nordenfelt

**Affiliations:** Lund University, Faculty of Medicine, Department of Clinical Sciences Lund, Infection Medicine, SE-22184 Lund, Sweden; Equipe Signalisation Calcique et Infections Microbiennes, Ecole Normale Superieure Paris-Saclay, 91190 Gif-sur-Yvette, France; Institut National de la Sante et de la Recherche Medicale (INSERM) U1282, 91190 Gif-sur-Yvette, France; Lund University, Skane University Hospital, Department of Clinical Sciences Lund, Nephrology, Lund, Sweden; Institute for Computational Science, University of Zurich, Winterthurerstrasse 190, CH-8057 Zurich, Switzerland

**Keywords:** group A streptococci, M protein, antibody binding, dual-Fab binding, antibody function, structural mass spectrometry, phagocytosis, in vivo model

## Abstract

Group A streptococci have evolved multiple strategies to evade human antibodies, making it challenging to create effective vaccines or antibody treatments. Here, we have generated antibodies derived from the memory B cells of an individual who had successfully cleared a group A streptococcal infection. The antibodies bind with high affinity in the central region of the surface-bound M protein. Such antibodies are typically non-opsonic. However, one antibody could effectively promote vital immune functions, including phagocytosis and *in vivo* protection. Remarkably, this antibody primarily interacts through a bivalent dual-Fab cis mode, where the Fabs bind to two distinct epitopes in the M protein. The dual-Fab cis binding phenomenon is conserved across different groups of M types. In contrast, other antibodies binding with normal single-Fab mode to the same region can not bypass the M protein’s virulent effects. A broadly binding, protective monoclonal antibody could be a candidate for anti-streptococcal therapy. Our findings highlight the concept of dual-Fab cis binding as a means to access conserved, and normally non-opsonic regions, for protective antibody targeting.

## Introduction

Antibodies are essential components of the immune system used to recognize and neutralize external intruders such as pathogenic bacteria. They are produced by B cells after their B cell receptor reacts with a specific antigen in the lymphoid tissue. B cell maturation and antibody responses have evolved to allow for an extraordinary variety enabling the binding to most foreign antigens. V(D)J recombination events, as well as somatic hypermutation, give rise to a vast repertoire of antibody variable domains (1, 2). B cell activation, clonal expansion, maturation, and class switching result in the generation of IgG antibodies that offer long term protection against infectious agents (3).

An IgG antibody is a Y-shaped molecule composed of two identical Fab fragments and one Fc domain, where the unique binding specificity is mediated via the Fab interaction. IgG typically binds the antigen with either one of the two Fabs. The two Fabs can bind to two copies of the same antigen to increase binding strength through avidity, a process that is dependent on antigen density and organization (4). We designate the latter form of binding as dual-Fab trans binding. When bound to their target, IgG molecules carry out effector functions by triggering clustering of Fc receptors on immune cells (5). This induces cell signaling and leads to a variety of downstream effects such as phagocytosis, immune recognition, and activation (6).

Group A streptococcus (GAS) is a common human pathogen causing significant morbidity and mortality in the human population and is an important causative agent of severe invasive infections (7, 8). The bacterium has evolved an extensive array of measures to counteract the human immune response (9), including resistance to phagocytosis (10, 11), and several immunoglobulin-targeting mechanisms (IdeS (12), EndoS (13), protein M/H (14)). The streptococcal M protein, a virulence determinant, has a long coiled-coiled structure with different regions (A, B, S, C, and D). These regions are typically associated with distinct protein interactions and bind many components of the humoral immune response (15, 16) such as C4BP and factor H, and forms complexes with fibrinogen (17) that can induce vascular leakage (18) and contribute to phagocytosis resistance (19, 20). The M protein can also reduce phagocytosis by reversing the orientation of IgG by capturing IgG Fc domains ((14, 21). These pathogenic mechanisms deprive the immune system of crucial defenses, allowing GAS to disseminate within a host and across the population. Additionally, the M protein has been implicated in autoimmune sequelae to streptococcal infections. Post-streptococcal sequelae such as rheumatic heart disease and rheumatic fever are due to its molecular mimicry to cardiac tissues (22), further contributing to the pathogenicity of this bacterium.

Although GAS infections generate a humoral immune response, repeated exposures seem to be required to generate protective memory B cell immunity (23). There are few candidates for anti-bacterial monoclonal antibody therapy in general (24), and none available for GAS. Much effort has been allocated to developing vaccines against GAS (25), with the prime immunizing antigen being M protein, particularly M protein-based peptides (26). Yet, no effective vaccine against GAS has been approved to date. It is unclear what makes it so difficult to generate immunity, but potentially formation of antibody subsets is suppressed by immunodominant regions (27) or the presence of cryptic epitopes (28). In severe life-threatening invasive GAS infections, intravenous IgG antibodies (IVIG) from human pooled plasma have been used as therapy, even though reports on their efficacy show contradictory results (29, 30, 31, 32). In the virus field it is well-established that certain antibodies have strong neutralizing abilities depending on epitope, binding angle, and glycan composition (33). So far, few studies have investigated the properties that constitute a protective antigen-specific antibody response against GAS.

Here, we have generated anti-M protein antibodies, derived from a healthy donor who had previously recovered from a GAS infection. When exposed to GAS and M protein, the antibodies bind and exert various effects, and we identify a new type of interaction where the two identical Fabs of one of the monoclonal IgG antibodies simultaneously bind to two distinct epitopes. We designate this form of bivalent binding as dual-Fab cis binding. Importantly, this broadly-binding antibody efficiently promoted all types of studied protective immune functions, including bacterial agglutination, NFkB activation, phagocytosis, and *in vivo* protection.

## Results

### Antigen-baiting allows the development of human single-cell derived M-specific antibodies

To understand what constitutes a protective antibody against GAS infection, we generated functional human antibodies and analyzed their effects on virulence. We chose M protein as a target antigen, with a donor that had successfully cleared a streptococcal infection as a source of M protein-specific antibodies. To identify human antibodies with specificity towards streptococcal M protein, we isolated CD19^+^ CD3^*−*^ IgG^+^ M^+^ B cells by baiting donor B cells with fluorescently conjugated M protein (**Fig. 1A**). The IgG+ B cells that did not react with the M protein could have specificities to any antigen, including other streptococcal antigens. Still, the focus of this work was on those B cells that were strongly reactive to M protein. The M protein we used for baiting is derived from an M1 serotype strain. We opted to use an M1 protein without knowing which serotype infected the original donor to increase the likelihood of identifying cross-strain reactive antibodies. Cloning RT-PCR of the variable regions of the heavy and light chains yielded ten antibody pairs (**Supp. Fig. 1A**). SDS-PAGE and mass spectrometry analysis of the antibodies expressed in HEK293 cells showed correct expression of the intact antibodies (**Supp. Fig. 1B**). Several antibodies showed clear reactivity to surface-bound M1 protein on GAS (strain SF370), the most common M protein among GAS isolates (**Fig. 1B**). Further experiments with three selected antibodies using a ΔM SF370 mutant strain lacking the M1 protein, demonstrated that binding of Ab25, 32, and 49 to the streptococcal surface was M protein-dependent (**Fig. 1C**). We could confirm that all three antibodies could be found in the donor serum through mass spectrometric analysis of proteotypic peptides.

**Fig. 1.**
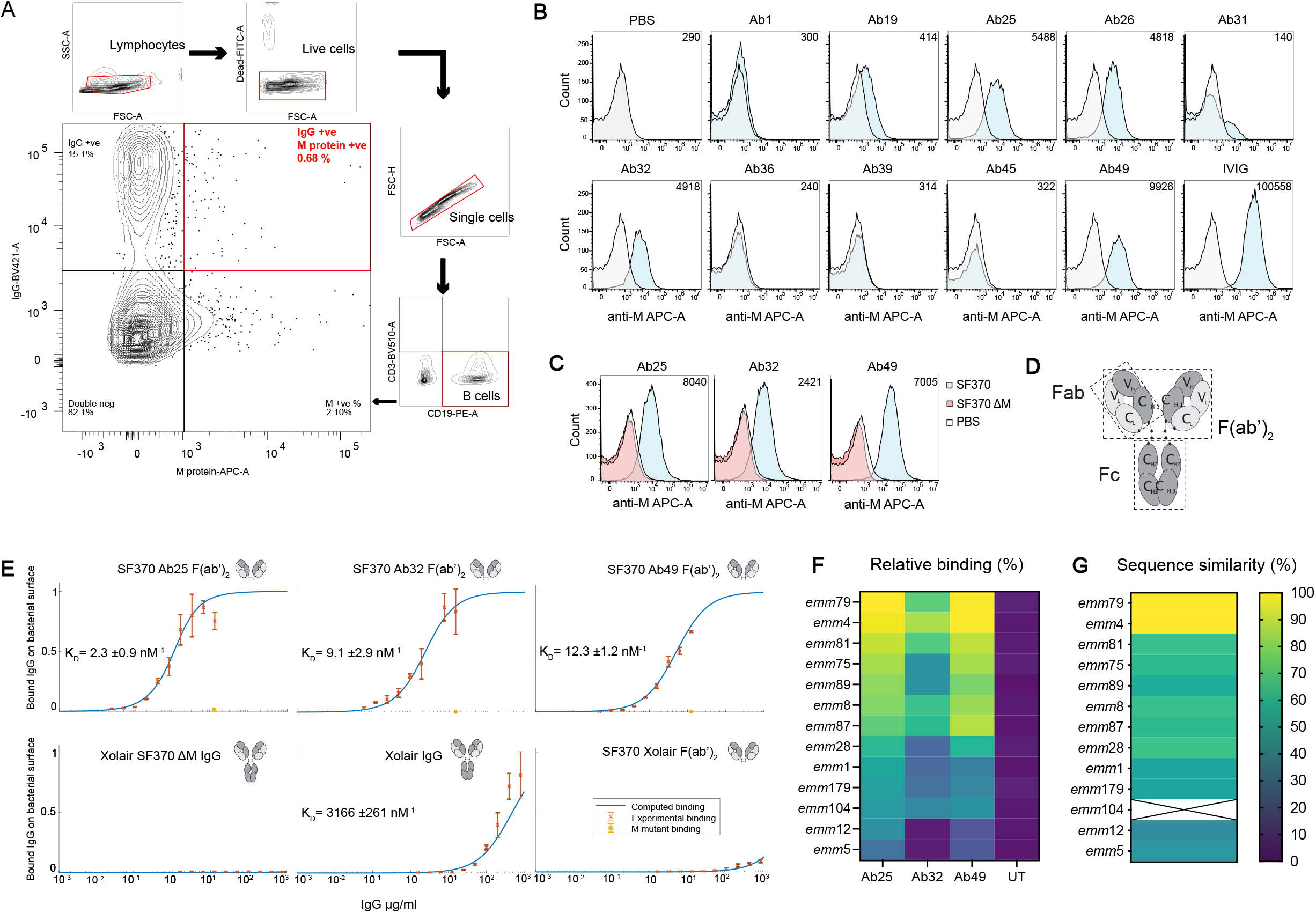
Antigen-baiting allows the development of human single cell-derived M-specific antibodies. **A** B cells were isolated using Rosettesep B and were FACS sorted into single lymphocytes which were live (Syto9-FITC negative), expressing CD19 (PE), lacking CD3 (BV510), and which were dually positive for IgG and M protein (BV410 and AF647, respectively). **B** SF370 (GFP-transformed) bacteria were stained with the shown antibody and a secondary Fab anti-Fab antibody (conjugated to AF647) was used to generate the signal for flow cytometric analysis. The number in the upper right corner of the flow histograms indicates the median fluorescence intensity. **C** The specificity of the successful antibodies was assessed by comparing the staining of WT SF370 and ΔM SF370. The bacteria were stained as in (b) with the aforementioned antibodies and were analyzed by flow cytometry. **D** A schematic representation of an IgG antibody. **E** Binding curves of antibodies to SF370 give an estimate of affinity to M-protein. The affinity of the specific antibodies is approximately 250-1500 times higher than that of IgGFc binding to M-protein. Each plot here shows the measured binding (red error bars) of specific antibodies to mid-log grown SF370 or the M mutant as a function of the total antibody concentration, together with a fitted binding curve using the least-squares method (blue curve). The bottom three plots are controls with a non-specific antibody that have been normalized according to the fitting of the bottom left binding curve, making the binding comparable across the three controls. KD values for the fitting are given in each plot, together with a confidence interval calculated using the Bootstrap method. The binding was measured using flow cytometry. The antibodies (antibody fragments) used are shown on the top-right side of each graph. The plots show no binding of the specific antibodies to the M mutant (orange error bar) at the highest measured total antibody concentration. **F** Different M serotypes of GAS were heat-killed and then prepared as in (B) and tested for reactivity against IdeS-cleaved Ab25, 32, 49 or untreated (UT, only secondary). The bacteria were analyzed by flow cytometry and the data was presented as a heat map. The displayed data is the combined result of three or more independent experiments. **G** Whole-genome sequencing was performed on the strains used in (F). The contigs were assembled and the *emm* sequence similarity was compared to *emm*79. The heat map shows the *emm* sequence comparison results. Sequencing of *emm*104 failed and is therefore designated by X.

We measured the antibody binding affinities to the surface of SF370 with either intact antibodies or F(ab’)_2_ fragments (**Fig. 1D**), the latter to avoid contribution from M1’s binding to IgGFc (Åkesson et al., 1994). Intact Xolair (Omalizumab, anti-IgE) showed a K_*D*_ of (3.2×10^*−*6^ M^*−*1^), signifying a low binding affinity in concordance with previous reported IgGFc affinity for purified M1 protein (3.4×10^*−*6^) (14). F(ab’)_2_ fragments of Ab25, 32, and 49 had considerably higher affinities for M1; 2.3×10^*−*9^, 9.1×10^*−*9^, and 12.3×10^*−*9^ M^*−*1^, respectively (**Fig. 1E**). To assess the reactivity of Ab25, 32 and 49 across different GAS M serotypes, we measured the binding of the antibodies to GAS *emm* serotypes 1, 4, 5, 8, 12, 28, 75, 79, 81, 87, 89, 104, and 179. We discovered that Ab25 and Ab49 have similar broad reactivity against these serotypes, with the strongest relative binding to *emm*4 and *emm*79, and with lowest relative binding to *emm*5 (**Fig. 1F**). The cross-reactivity pattern was different for Ab32, including lack of binding to some strains. Whole-genome sequencing of the strains was performed, and genome contigs were assembled. When comparing the DNA sequences of the *emm* genes used in Fig. 1F strains, we saw that overall genetic similarity could serve as a good predictor of relative antibody binding strength (**Fig. 1G)**.

### Characterization of anti-M antibodies

To characterize the identified anti-M antibodies, we performed a panel of biochemical and immunological assays. We used structured illumination microscopy (SIM) immunofluorescence (IF) to visualize the anti-M binding pattern on the surface of SF370. Binding to the M protein shows a similar punctate distribution along the surface of the organism with all the monoclonal antibodies, including the Fc-mediated Xolair binding (**Fig. 2A**). IVIG, which contains pooled IgG from thousands of donors, instead stained the whole surface of the bacteria indicating expected polyspecific coverage (**Fig. 2A**). Ab25 showed the best reactivity with M protein using an anti-M ELISA (**Fig. 2B**), whereas Ab32 showed the best binding to M protein in Western blot (WB) experiments (**Fig. 2C**). Taken together, the data from IF, ELISA, and WB indicate that the three monoclonals have different modes of binding to the M1 protein, most likely due to different M protein conformations under the assay conditions.

**Fig. 2.**
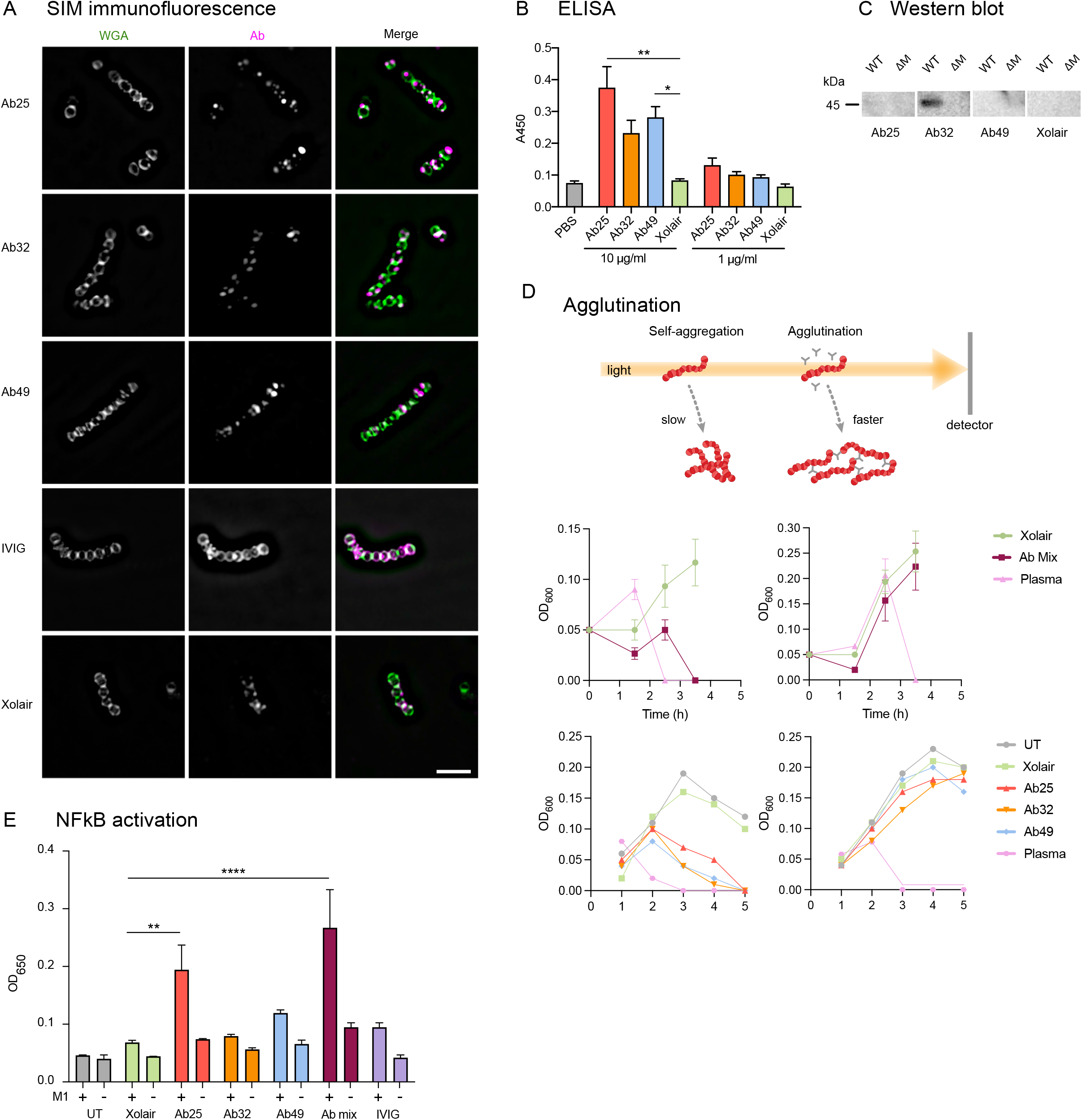
Figure 2. Characterization of anti-M antibodies. **A** Structured illumination microscopy (SIM) super-resolution imaging was performed on bacteria stained with IdeS cleaved α-M antibodies and probed with Dylight488-conjugated Fab α-Fab fragments. AF647-conjugated WGA was used as a counter-stain to highlight the bacterial structures. The scale bar represents 5 μm. **B** Antibodies were tested for reactivity against the M protein in an ELISA assay where the M protein was immobilized onto the well surface. Xolair and PBS were used as negative controls. The absorbance at 450 nm was measured and plotted as an average of quadruplicate wells. **C** Protein lysates from logarithmically grown WT and ΔM SF370 bacteria were run on SDS-PAGE and, after blotting, were probed with the mentioned antibodies at a concentration of 2 μg/ml. **D** Time course analysis of the effect of the antibody mix or individual antibodies used at 100 μg/ml each on bacterial agglutination was assessed by OD_600_ measurements. **E** THP-1-Blue cells were treated with M1 protein (2 μg/ml) with or without anti-M specific antibodies. The data shown are from triplicate samples, representing experiments done three times. Error bars represent the SD. Statistical significance was evaluated using a two-way ANOVA with Dunnett’s multiple comparison correction. * denotes p 0.05, ** for p 0.01, *** for p 0.001 and **** for p 0.0001).

Antibody-mediated bacterial agglutination is a well-documented antibody function and has important biological significance such as enchaining bacteria for effective immune clearance (34, 35). Another well-known interbacterial, GAS-specific phenomenon is the formation of M-dependent bacterial aggregates at the bottom of the growth tube (36). While it is impossible to grow GAS without self-aggregation, the antibodies greatly enhanced bacterial agglutination. Both the triple antibody cocktail and individual antibodies led to dose-dependent agglutination, as is the case with donor serum from the patient from which the M-reactive B cells were obtained (**Fig. 2D, Supp. Fig. 2B**). This agglutination was not observed for the ΔM strain, further validating that the antibody-dependent agglutination is an M-specific phenomenon. Serum, however, can agglutinate GAS without binding to the M protein as it contains antibodies targeting non-M surface proteins. Agglutinated bacteria and the typical GAS aggregates could be dissipated by vigorous vortexing in the presence of the anti-M antibodies or plasma (**Supp. Fig. 2C**). GAS agglutination and aggregate dissolution (after vortexing) were most pronounced in Ab25, 49, and lesser with Ab32, while Xolair (with only IgGFc-binding) had no effect. This effect has not been seen before and its importance is unknown.

By ligating their antigens and mediating antigen uptake, antibodies also activate macrophages leading to prion-flammatory cytokine production (37). We addressed the antibody-dependent M protein-induced immune activation using THP-1 X-Blue reporter cells, which secrete SEAP (secreted embryonic alkaline phosphatase) as a quantitative indicator of NF-B activation. We found that M protein alone cannot induce NF-kB signaling. However, combining M protein with Ab25 led to a significant 2.8-fold increase in NF-kB activation compared to M protein with Xolair (**Fig. 2E**). Combining M protein with Ab49 had a modest effect on NF-kB activation (1.6-fold, ns), whereas its combination with Ab32 had no impact (1.2-fold). Combining all three antibodies led to a significant cumulative 3.9-fold increase in NF-kB activation, probably due to the combined amount of Ab25 and 49. Interestingly, we found that IVIG does not elicit the same antibody-mediated NF-kB activation upon THP-1 X-Blue exposure to the IVIG-treated M protein (1.4-fold).

Some anti-M protein antibodies have been known to cross-react with human cardiac tissues. We tested the autoreactivity of our antibodies using tissue microarrays and saw no detectable cross-reactivity with any human tissue (**Supp. Fig. 3A, B**). The only antibody which showed a clear level of autoreactivity with multiple tissue samples (seen as brown color) was the Troponin-specific antibody. Troponin is a cardiac protein and we used an anti-Troponin antibody to replicate a cardiac-reactive antibody. The monoclonals’ combined biochemical and immunological characterization shows that all are specific for M protein, bind to different epitopes, and can induce immunological effects without being autoreactive and that Ab25 has the most potent effect overall.

### Anti-M antibody promotes efficient phagocytosis

Phagocytosis is a receptor-mediated process where prey are internalized into phagosomes, followed by their maturation into acidic, hostile compartments (38). To investigate the ability of the antibodies to trigger phagocytosis, we used persistent association-based normalization (39) to study both the antibodies’ ability to increase phagocyte association as well as internalization. We incubated phagocytic THP-1 cells with pH-sensitive CypHer5-stained bacteria (**Supp Fig. 3A**) at increasing multiplicities of prey (MOP). We combined Ab25, 32, and 49 to assess their cumulative effect on bacterial association with THP-1 cells. Compared to Xolair, the antibody mix and, to a lesser extent, IVIG modestly increased the overall association of THP-1 cells to the bacteria (**Fig. 3A**). This can be seen as a left shift in the curve, meaning fewer bacteria are required to achieve maximal association. When tested individually and assaying the number of bacteria per cell, only Ab25 showed a statistically significant increase in association of bacteria to phagocytes, indicating that the antibody mix-mediated increase in the association is solely due to Ab25 (**Fig. 3B**). The antibody mixture increased internalization compared to Xolair, and the divergence between the two treatments was increased as a function of MOP (**Fig. 3C**). Upon more detailed examination of individual antibodies, only Ab25 showed a statistically significant increase in internalization (**Fig. 3D**). Dose-response analysis showed that Ab25 is significantly more effective than Xolair in mediating internalization (concentration at which 50% of THP-1 cells have internalized bacteria; EC_50_ 0.8 vs. 40.2 μg/ml) (**Fig. 3E**). Expanding on our previous cross-strain reactivity data (**Fig. 1F**), we used Ab25 to set up an imaging-based binding assay (**Supp. Fig. 5A, 5B**). By utilizing multiple strains, we show that Ab25 has broadly binding potential against GAS compared to Xolair (**Supp. Fig. 5C,5D**).

**Fig. 3.**
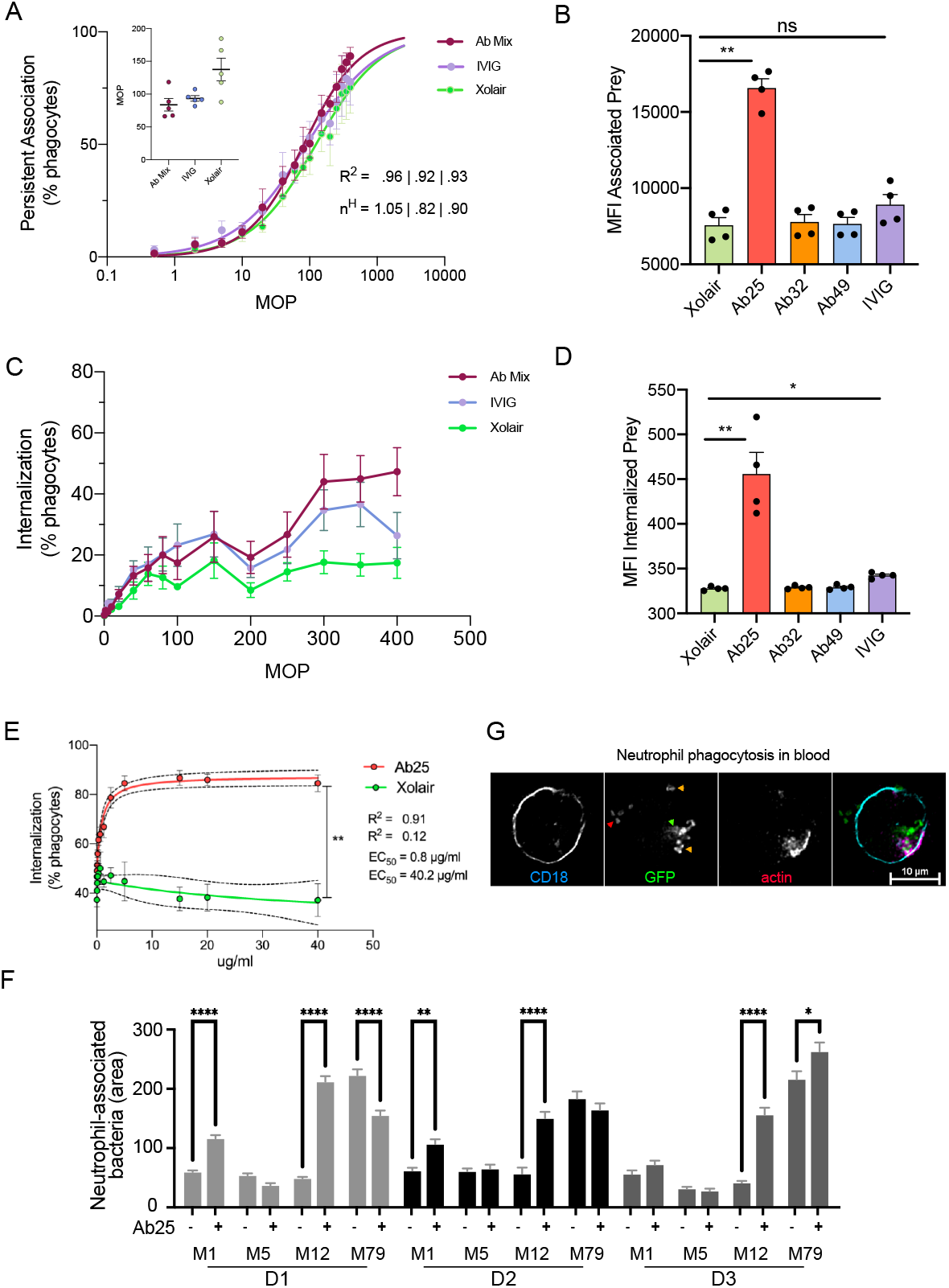
Anti-M antibody promotes efficient phagocytosis. **A** THP-1 cells were incubated with increasing MOPs of heat-killed SF370 bacteria (opsonized with 10 μg/ml Xolair, anti-M antibody mix, or IVIG). The THP-1 cells were allowed to associate with and internalize the bacteria for 30 minutes before flow cytometric analysis. The curves represent the percentage of cells associated with bacteria as a function of the MOP. The inset displays the MOP50 for each opsonization condition. (nH) represents the Hill coefficient). **B** The MFI of THP-1 cells in the FITC channel indicates the effect of each antibody on bacteria association to THP-1 cells at MOP 400. **C** THP-1 cells were incubated as in (a), but only the percentage of cells with internalized bacteria were plotted for each MOP. **D** MFI of THP-1 cells which had internalized bacteria pre-opsonized with 10 μg/ml individual antibodies, are presented at an MOP of 400 (internalized bacteria channel). **E** Heat-killed SF370 opsonized with Ab25 or Xolair at concentrations of (0.017-40 μg/ml) were incubated with THP-1 cells at MOP 300. Phagocytosis was assessed as in (b). The data shown in this figure is from the pooled results of three independent experiments. **F** Whole blood phagocytosis was performed with samples from 3 donors (D1, D2, and D3). The blood was diluted and then infected (MOP 10) with live GFP-expressing bacteria (pre-opsonized with Xolair or Ab25 (10 μg/ml)) for 30 mins before PFA fixation. The cells were then stained with SiR-Actin (Em:674), and anti-CD18 (BV421, blue) were immobilized on CD29-coated glass plates and imaged with 20X magnification. From each donor, the area of GFP signal associated with at least 440 cell data points were recorded. The data shown represent at least 560 cell events collected per condition. Similar results were seen in two independent experiments (4 donors). **G** An example image of Ab25-opsonized M12 bacteria infecting a neutrophil with internalized bacteria (green arrow), extracellular bacteria (red arrow), and active internalization events (yellow arrow, indicated by actin polymerization) is shown. The cell was imaged with 60X magnification. Error bars represent the SEM. Statistical significance was assessed using one-way ANOVA with Kruskal-Wallis multiple comparison correction and * denotes p 0.05, ** for p 0.01, *** for p 0.001, and **** for p 0.0001

The phagocytosis data showed that despite strong Fab-mediated binding and induction of other immunological effects by all monoclonals, only one antibody, Ab25, can promote phagocytosis of group A streptococci. We have shown earlier (**Fig. 1F**) that Ab25 possesses broadly reacting potential against different M types. For that reason, we tested its ability to opsonize and mediate the phagocytosis of non-M1 serotypes. Opsonization with Ab25 led to increased phagocytic efficiency of the M1 (SF370 and AP1), M12, and M89 serotype strains, whereas the M5 was efficiently phagocytosed by THP-1 cells regardless of whether a binding opsonin was employed or not (**Supp. Fig. 4C**). We established a microscopy-based (**Supp. Fig. 6A**) whole-blood phagocytosis assay to validate our *in vitro* findings with THP-1 cells. Upon infecting blood with antibody-opsonized live GFP-expressing M1, M5, M12, and M79 strains, we studied the effect of antibody opsonization on the association between the bacteria and the CD18+ cells (**Fig. 3F**). The phagocytosis enhancement was most pronounced in M1 and M12 (**Fig. 3F, Supp. Fig. 6B**). This effect was not present in M5, which is in line with our previous data from THP-1 cells (**Supp. Fig. 4C**). The Ab25-mediated opsonization of M79 was donor-dependent as one donor showed decreased phagocytosis upon bacterial opsonization, and one donor showed enhancement. Donor variability was also seen with the M1 strain, albeit less pronounced than with M79. We assessed the interaction between the CD18+ cells (neutrophils in this case) with the GFP-expressing Ab25-opsonized bacteria. We saw that a large proportion of the neutrophils had internalized bacteria, and an example is shown in **Figure 3G**. Overall, the results from opsonization and phagocytosis show that Ab25 can bind to and opsonize multiple M strains and that this leads to internalization by human phagocytes.

### Anti-M antibody protects mice from GAS infection

The induction of phagocytosis and NF-kB, as seen with Ab25, are important indicators of immune function. To test the potential protective effects of Ab25 *in vivo*, we used a mouse model of subcutaneous infection with GAS. We opted to only test Ab25 in our *in vivo* model since it had the strongest *in vitro* immune activity. The mice were pretreated with intraperitoneal injections of Ab25 or IVIG. High-dose IVIG has been used in mice models of severe GAS infections (40) and served as a positive control. Treatment with IVIG or Ab25 reduced the bacterial burden in the spleen, kidney, and liver compared to untreated controls, with Ab25 exhibiting better protection than IVIG (**Fig. 4A**). Ab25 or IVIG treatment also reduced the cytokine mobilization of TNFα, MCP-1, and IL-6 in plasma (**Fig. 4B**). The levels of IFNγ, IL-10 and IL-12p70 were below the level of detection under our experimental conditions. Taken together, the agglutination, NF-kB, phagocytosis, and animal experiments show that Ab25 has an immunomodulatory effect, which can protect an animal from GAS infection.

**Fig. 4.**
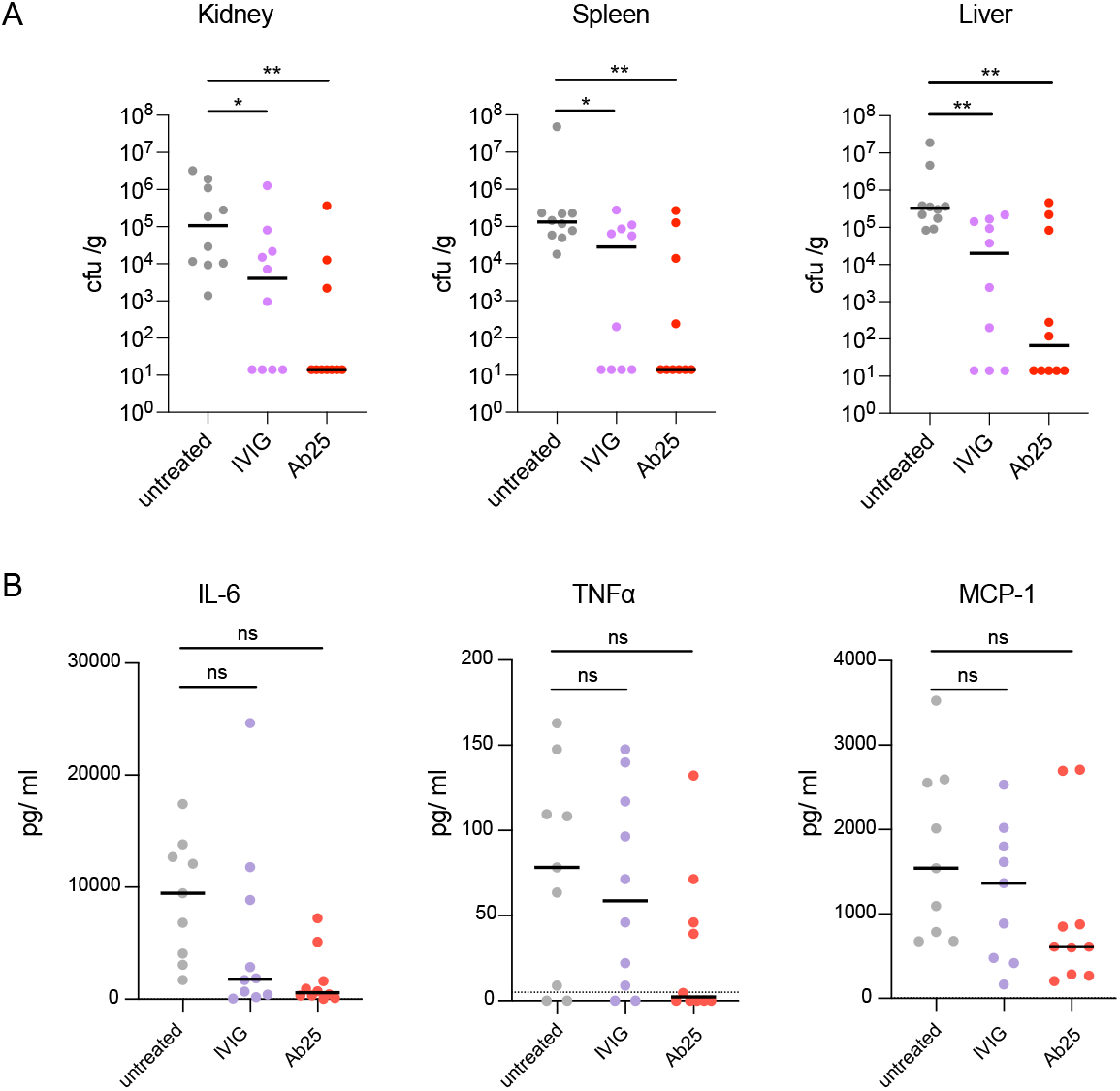
Anti-M antibody protects mice from GAS infection. C57/BL6 mice were pretreated with 10 mg IVIG or 0.4 mg Ab25 per mouse before subcutaneous infection with 10^6^ CFU of GAS (*S. pyogenes* AP1). The animals were sacrificed after 24 hours. **A** The bacterial burden in spleen, kidney, and liver tissue was measured by colony counts (CFU/gram of tissue). **B** Cytokine levels in plasma were assessed using a cytometric bead array. The data are pooled from two independent experiments (5 mice per condition, per set). Statistical significance was evaluated using one-way ANOVA with Kruskal-Wallis multiple comparison corrections and * denotes p 0.05, ** for p 0.01, *** for p 0.001, and **** for p 0.0001.

### Structural epitope characterization reveals bivalent dual-Fab cis binding mode of interaction

Antibodies that bind via their Fabs with high affinity are typically expected to promote an immune response. However, only Ab25 promoted all the tested immune effector functions. To assess structural differences and their mode of binding, we performed targeted cross-linking coupled to mass spectrometry (TX-MS) (41) of the M-protein and the antibodies.

TX-MS first models quaternary conformations of protein complexes that are, in a second step, confirmed by targeted cross-linking analysis of the most high-scoring models. The high-resolution models of the Fab fragments of Ab25, Ab32, and Ab49 binding to their respective epitopes on M protein (**Fig. 5**) revealed a high degree of structural similarity between Ab25 to Ab49 over the CDR H3 loop (RMSD 1.0Å), whereas the conformation of Ab32 CDR H3 is more divergent (**Fig. 5B-C**). The conservation at the primary amino acid sequence level of Ab25 and Ab49 is less evident (**Fig. 5A**). The subsequent targeted cross-linking analysis resulted in the identification of ten cross-linked peptides between Ab25 and M1-protein. These cross-links are found between the F(ab’)_2_ and two different regions on the M-protein, indicating that Ab25 has two different binding-sites in the B-S-C region (**Fig. 5D** and **Fig. 5F, Supp. Table 1, Supp. Fig. 7A-J**). Superimposing the cross-linked distant constraints onto high-resolution docking models, shows that Ab25 F(ab’)_2_ can simultaneously bind the two cross-linked epitopes without inducing large conformational changes in the hinge region. This implies that Ab25 is capable of dual-Fab binding in an intramolecular bivalent cis-binding fashion to two distinct, non-identical epitopes. In contrast, the cross-link analysis of M-Ab49 generated two unique cross-linked peptides confined to only the upper epitope found with Ab25 (**Fig. 5E, Supp. Table 1, Supp. Fig. 7K-L**).

**Fig. 5.**
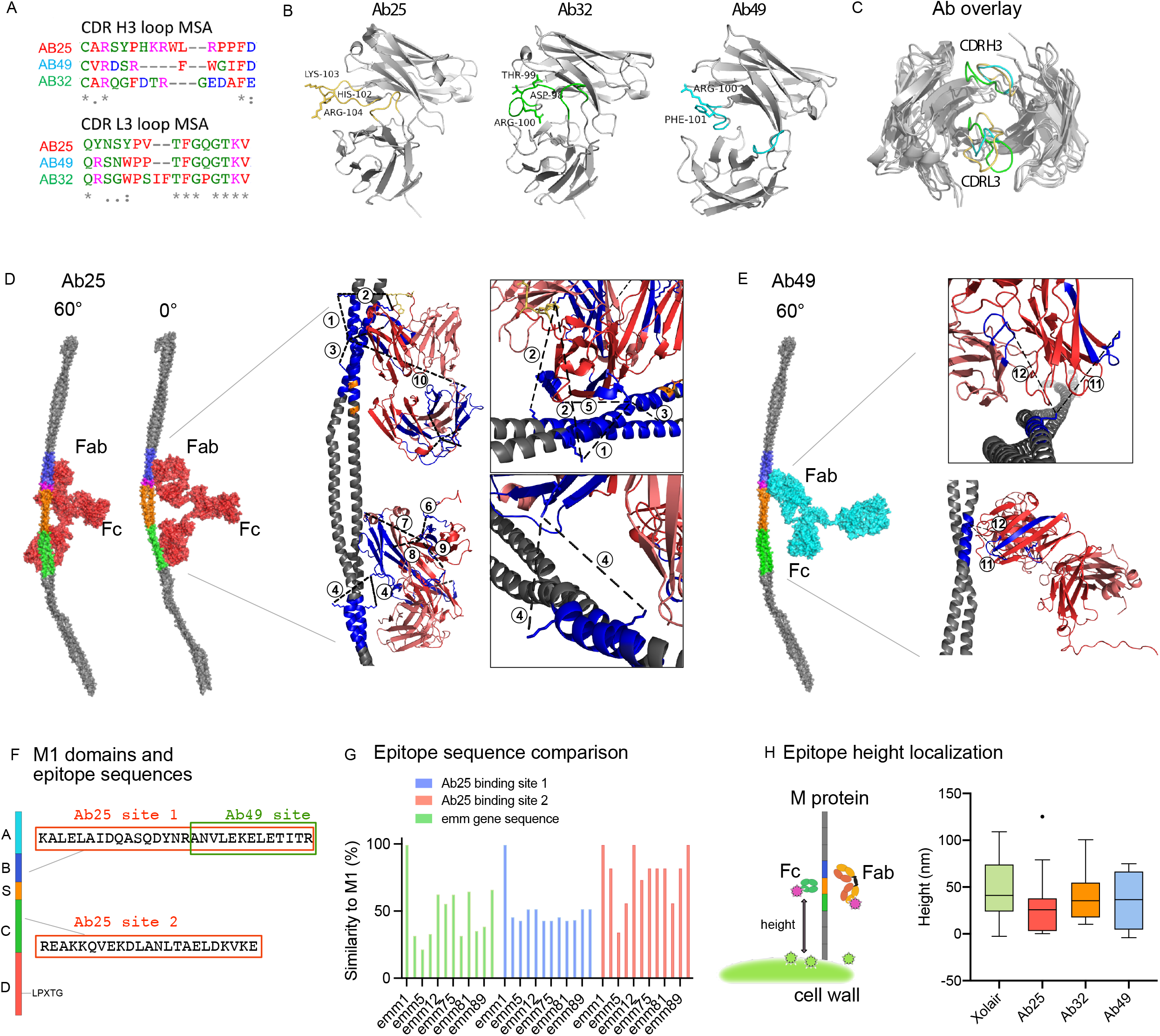
Structural epitope characterization reveals bivalent dual-Fab cis binding mode of interaction. **A** Multiple sequence alignment reflects the L3-H3 loop differences on the amino acid sequence level. **B** Representations of the Fab fragments of the modeled antibodies Ab25, Ab32, and Ab49. The CDR H3 and L3 loops are indicated (Ab25, yellow; Ab32, green; and Ab45, turquoise). **C** Structural alignment of Ab25 (yellow CDR loops), Ab32 (green loops), and Ab49 (turquoise loops). **D** The M1 interaction sites with Ab25 (red) reveal the binding epitopes. A rotation of 60°along the y-axis has been applied between the views. The contact surface length is 45Å in the upper binding site (B2-B3-domain) and 18Å in the lower (C-domain). **E** The M1 interaction site with Ab49 (turquoise) is presented. The length of the contact site for Ab49 (B2-B3 domain) is approximately 16Å. Insets in D and E represent a full-length model generated by TX-MS (41), showing the Fab-mediated interaction of Ab25 or Ab49 (heavy chain in dark red, light chain in a lighter shade) with the M1 protein (in gray). Crosslinked peptides are indicated in dark blue and crosslinks between lysine (K) residues as dashed black lines. The CDR H3 loop is shown in yellow (for Ab25) or red (for Ab49). There are two interaction sites for Ab25 on the M1 protein, the upper one in the B2-B3-domain (inset) supported by 4 crosslinks and the lower one in the C1-domain (inset) by 6 crosslinks. Only one interaction site for Ab49 with M1 exists and is supported by 2 crosslinks. **F** Schematic presentation of the M1 protein with the two binding crosslinks for Ab25 (binds both) and 49 (binds upper) being displayed. **G** Sequence similarity on genetic sequences was performed to compare the two identified epitopes across strains. **H** The M protein is associated with the bacterial cell wall through the LPXTG motif in the C-terminal D domain. The Fab epitopes and the Fc-binding region on the M protein are estimated by a fluorescence localization averaging method using structured illumination microscopy (SIM) images (44). A relative binding site is determined by resolving the distance between the antibody and a reference channel (WGA, binds cell wall) using the cumulative signals from many images. The results shown here are from 4 independent experiments with N=9,13,9,9, respectively.

The amino acid sequences corresponding to the M proteins obtained from whole-genome sequencing are shown in **Supp. Data 1**. Amino acid alignment and sequence comparison of the different M proteins is shown in **Supp. Data 2**. The domains of the M1 protein are shown in **Figure 5F** and the two identified epitope peptides for Ab25 and 49 are highlighted. Ab49 only interacts with the upper epitope, whereas Ab25 interacts with both. Based on our genetic sequencing shown before (**Fig. 1G, Supp. Data 1-2**), we displayed the *emm* sequence similarity as a percentage of M1 (**Fig. 5G**). In the same panel we also compared the conservation of the Ab25 binding epitopes on the M protein across the strains. We saw that the lower binding site (binding site 2) for Ab25 is relatively conserved across strains, more so than the upper site (binding site 1) (**Fig. 5G**). Overall, both sites’ conservation when compared to *emm*1 is higher than the general *emm* strain sequence conservation. The *emm* types can be divided into patterns (A-C, D, and E) (42), belonging to different clades and clusters (43), and this classification shows that the majority of the strains used here belong to the E pattern type **Supp. Table 2**, but that *emm*1 belongs to a different clade. Since the strongest Ab25 binders have no clearly conserved regions in the upper binding site, it is likely that the original upper epitope is not found in a B repeat, but that it has cross-reactivity to the B2-B3 repeats in A-C pattern *emm* types. This could be explained by structurally similar and appropriately charged motifs which allow for the stable dual-Fab cis binding.

To further investigate the differences in the observed binding sites, we used site-localization microscopy (44), where the relative distance between the fluorescently labeled cell wall and antibody binding epitopes was determined by repeated measures of multiple individual bacteria (**Fig. 5H**). The height analysis showed that all monoclonal F(ab’)_2_ fragments bind close to the Fc binding domain (S) on the M1 protein (compared to Xolair Fc binding), which supports the TX-MS crosslinking-based results.

### Dual-Fab cis mode of interaction is required for functional antibody binding

Intriguingly, Ab25 binds across the M1 protein S-region (**Fig. 5D,F**) that has previously only been associated with Fc-mediated binding (Åkesson et al., 1994; Nordenfelt et al., 2012). Since the antibodies appear to interact with epitopes close to the Fc binding S domain (as seen in (**Fig. 5D-H**), we further investigated whether the antibodies could interfere with Fc binding. The dual-Fab cis binding of Ab25 covers the S domain (colored in orange in (**Fig. 5D**) and would obstruct Fc binding, whereas single Fab interactions would lead to minor or no interference with Fc-binding. We measured the binding of fluorescent Xolair to SF370 bacteria that had been preincubated with antibody samples (**Fig. 6A**). Both blood plasma from the original B cell donor and Ab25 significantly obstructed Fc-binding, whereas IVIG, Ab32, and Ab49 did not. Since Ab49 and Ab25 share one similar epitope, located above the S region, it indicates that binding there alone is not sufficient to break the Fc interaction, and strongly suggests that this is due to a dual-Fab cis binding quality of Ab25.

**Fig. 6.**
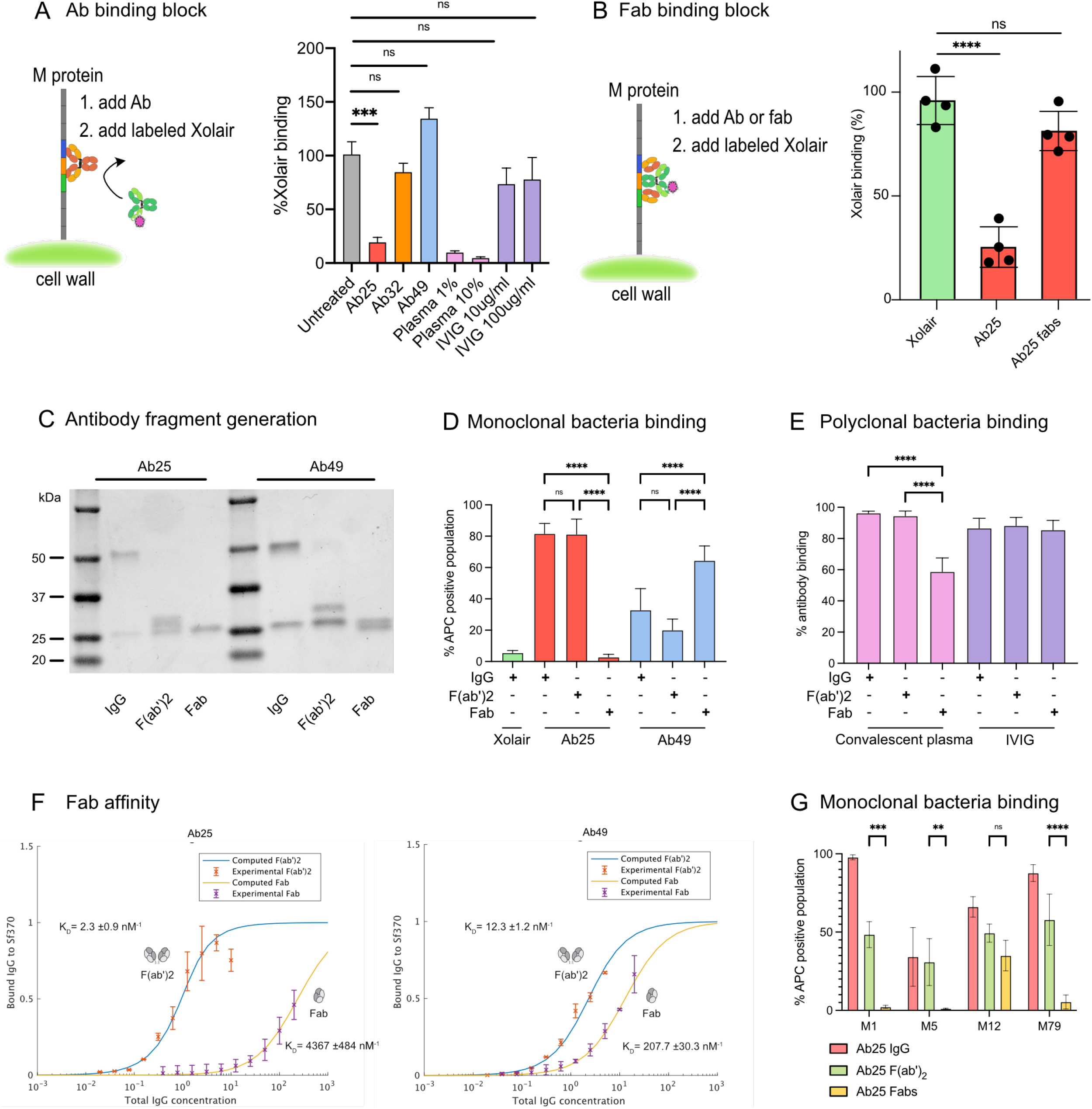
Dual-Fab mode of interaction is required for functional antibody binding. **A** Binding of Alexa647-conjugated Xolair (100 μg/ml) to SF370 after pre-treatment with indicated antibody samples (10 μg/ml). The data is from 3 independent experiments pooled together. **B** Heat killed bacteria previously stained with Oregon Green were incubated with 10 μg/ml of Ab25 or IgdE-digested (Spoerry et al., 2016) Ab25 (Fabs). The bacteria were then incubated with fluorescently labeled Xolair (AF647) before being analyzed with flow cytometry. Bacteria that bound Xolair via its Fc binding are shown as positive. **C** SDS-PAGE analysis of the antibodies used for the binding analysis. **D** Binding properties of Xolair, IdeS-cleaved or IgdE-cleaved Ab25 or Ab49 were studied as previously by mixing the antibodies with heat-killed, Oregon Green bacteria. The bacteria were then washed and stained with AlexaFluor 647-conjugated Fab anti-Fabs and analyzed by flow cytometry. Data is from four independent experiments with mean and SD shown. **E** Binding properties of IdeS-cleaved or IgdE-cleaved convalescent donor plasma or IVIG were studied as in D. Data is from four independent experiments with mean and SD shown. **F** Affinities for Fabs generated by IgdE treatment of Ab25 and Ab49 were performed as described previously in Fig. 1. The new Fab data is overlaid on the data from Figure 1. The data represent pooled results from four independent experiments. **G** Binding properties of IdeS-cleaved or IgdE-cleaved Ab25 were studied by mixing the antibodies (20 μg/ml) with indicated strains of live GFP-expressing bacteria. The bacteria were then washed and stained with AlexaFluor 647-conjugated Fab anti-Fabs and analyzed by flow cytometry. Data is from three independent experiments with mean and SD shown. Statistical significance was evaluated using two-way ANOVA with Tukey multiple comparison corrections and * denotes p 0.05, ** for p 0.01, *** for p 0.001, and **** for p 0.0001.

Dual-Fab cis antibody binding of two identical Fabs to two different epitopes on a single protein is, to our knowledge, a novel, previously undescribed mode of antibody interaction. Dual-Fab cis binding is also associated with an apparent gain in immunological protective function. We took multiple approaches to verify these findings and elucidated the precise nature of Ab25’s dual-Fab binding capacity. We investigated the ability of single Ab25 Fabs to obstruct Fc binding. We used IgdE protease (45) to prepare intact single Fabs of Ab25. If the binding of single Fabs to either Ab25 binding site on M protein could sustain a steric hindrance, we should see a reduction in Fc binding. However, Fc binding was not affected by single Ab25 Fabs (**Fig. 6B**), suggesting that binding occurs via dual-Fab binding. Full cleavage of the antibodies into Fabs was verified by SDS-PAGE analysis (**Fig. 6C**). We wanted to see how well different forms of the antibodies could bind to bacteria. We compared the binding of whole IgG, F(ab’)_2_, and single Fabs to SF370. Strikingly, Ab25 single Fabs could not bind to the bacteria, whereas Ab49 Fabs increased their binding (**Fig. 6D**). The latter is the expected result since single Fabs should have easier access, yet the results reveal that Ab25’s predominant mode of interaction with M protein is via dual-Fab cis binding. We attempted to perform M-peptide ELISA experiments to further validate the dual-Fab cis binding but only full-length M protein worked, so no further details could be extracted through that approach (data not shown). This is not surprising as the complexity of the M protein folding has been well-established (46, 47).

The results using IgG-cleaving enzymes provide a possible way to investigate the occurrence of dual-Fab cis binding antibodies in polyclonal samples. By treating convalescent donor plasma and IVIG with IdeS and IgdE, we saw no effect on binding with F(ab’)_2_compared to intact IgG (**Fig. 6E**). However, we saw a significant reduction (around 40%) in binding only with the convalescent plasma when comparing single Fabs to IgG (**Fig. 6E**). It should be noted that the IgdE enzyme only cleaves IgG1. This indicates that there could be a significant portion of dual-Fab cis binding antibodies in GAS-convalescent individuals. To assess how much weaker the binding via single Fabs is, we performed affinity measurements of single Fabs. These showed that Ab25 Fab binding has a 2000-fold lower affinity (4.4 μM^*−*1^) compared to F(ab’)_2_binding (2.3 nM^*−*1^, (**Fig. 6F**). There was still a question of whether dual-Fab cis binding occurs across different M types. To answer this, we performed similar binding experiments to those in **Figure 6D** but used live bacteria expressing GFP. We found that single Fab fragments of Ab25 lost binding to both M5 and M79, as seen with M1, while they retained binding to the M12 strain. In contrast, binding with F(ab’)_2_occurred for all the strains (**Fig. 6G**). This means that for M12 we cannot determine the mode of interaction, only that both single and dual-Fab are possible. Our data shows also that for the A-C *emm* pattern types (such as M1 and M5 strains) and for the E *emm* type strain M79, Ab25 only binds via dual-Fab interaction.

## Discussion

In this study, we describe a new broadly-binding monoclonal antibody (Ab25) that binds to the M protein’s central region and mediates bacterial clearance. Interestingly, this antibody has a peculiar mode of interaction with its target as it interacts with the M protein at two distinct epitopes simultaneously. We now understand that in addition to the classical antibody-antigen interactions, which are extensively studied, dual-Fab binding of antibodies to antigens could potentially enhance antibody function. Mass spectrometric analysis of anti-M antibodies cross-linked to M protein, coupled with biochemical and immunological analysis, revealed that dual-Fab cis binding represents a bivalent interaction around the M protein S region. Importantly, our data show that dual-Fab cis biding can occur across different M types, even when their sequences are not highly conserved. We confirmed the dual-Fab cis binding phenomenon using three independent assays (crosslinking TX-MS, inhibition of Fc-mediated binding, and loss of single-Fab binding). This is a testament to the complexity of antigen-epitope interactions where binding requires recognizing compatible 3D structures.

This antibody-M protein interaction can mediate phagocyte engagement and streptococcal phagocytosis. As seen with Ab25, dual-Fab cis binding seems to be a contributing factor in the triggering of an immune response. In contrast, as seen with the closely located Ab49, single-Fab binding has little or no immunological effect in the assays tested. It has been previously shown that antibodies against the hypervariable region or C-repeats (10, 16, 48, 49) of the M protein can also be opsonic and mediate phagocytosis. We employed two independent approaches to assay phagocytosis, using THP-1 cells, and human blood infections. THP-1 cells are incubated with heat-killed bacteria and are analyzed by flow cytometry. The blood-based experiment, however, utilizes live bacteria and whole donor blood. This creates a more complex system that can discern nuances that require multiple cell types or molecules, such as complement activation. Both experimental approaches suggest that Ab25 is a potentially opsonic antibody across multiple GAS *emm* types. Therefore, while in the context of the M protein central region, dual-Fab binding antibodies are opsonic, other functional interactions between antibodies and antigens clearly exist. It is interesting to note that earlier studies on M6 protein revealed the central M region to not only be conserved amongst many M proteins but also non-opsonic (48), mirroring the results we observe with Ab32 and Ab49. The dual-Fab interaction of Ab25 is correlated with it being opsonic while still retaining the cross-reactivity with M proteins due to the conserved nature of the central M region.

The fact that Ab25 and Ab49 bind to such similar locations, yet their binding leads to such different outcomes, provides both an excellent internal control for dual-Fab cis binding and prompts questions as to why it is beneficial for immune stimulation. From a structural perspective, a two-point attachment is much more stable than a one-point attachment, probably stabilizing IgGFc-Fcγ receptor interactions. It would also provide a consistent angle of interaction, which might aid the clustering of receptors. The angle is relevant because the presentation of IgGFc’s pointing in different directions could interfere with the zipper-like mechanism that Fcγ receptors typically use during phagocytosis (50). The specific orientation that Ab25 adapts when binding M protein results in the IgGFc being oriented perpendicular relative to M protein. This happens to be an optimal angle at which Fcγ receptors interact with IgGFc (51). Instead, interactions between IgA and Fcα receptors would have benefitted from an antibody being rotated 90° from that of Ab25 (52). Recent findings indicate that both antigen height, where below 10 nm is beneficial (53) as well as pericellular barriers where transmembrane pickets obstruct the interaction (54) are important factors to consider for effective phagocytosis. Given that all three of our monoclonals bind at a similar height (40-50 nm), this in and of itself cannot be an explanation here. However, it is possible that the stable dual-Fab interaction shifts the perceived antigen height and thus results in a closer bacteria-phagocyte interaction. In this case, the pericellular barrier would be reduced by the same distance, allowing a more effective phagocytic interaction. Another obvious mechanism could be the reduced IgGFc binding that Ab25 achieves by spanning the IgGFc binding site (obstructing the S region). However, this is most likely not the case for two reasons. First, the IgGFc binding has a low affinity in the μM range and only becomes relevant at high IgG concentrations, magnitudes larger than those used in the phagocytosis experiments. Second, if the Fc binding could block phagocytosis, then the phagocytosis experiments with mixed monoclonals would see an additive effect, as Ab25 would block Fc binding, allowing the other antibodies to function. As we saw no additive effect whatsoever, this cannot be the case. The reduced IgGFc binding that Ab25 confers might have additional protective effects on the host that remain to be investigated in future studies.

The requirement of dual-Fab cis binding to separate epitopes that Ab25 exhibits, as demonstrated by experiments with single Fabs, is an unexpected mode of functional antibody interaction. Dual-Fab cis binding is thus adding to the already astounding diversity found in antibody binding mechanisms (55). A related phenomenon to the dual-Fab binding is the case of the anti-HIV 2G12 antibody (56), which has a mutation in its hinge region leading to Fab dimerization (57). The two fabs of 2G12 bind to high-mannose sugars but due to the fact of their unorthodox dimerization, essentially behave as one large Fab (58). In fact, normal single Fab-based interactions between multiple anti-HIV glycan antibodies gave similar biological outcomes as 2G12 (for a review see (59)). This indicates that while 2G12 has an unorthodox structure, its function is correlated to the specific epitope and not due to a distinct mode of interaction. In the context of unorthodox antibodies; bispecific antibodies (60) or antigen clasping antibodies (61) have been engineered for improved functionality. We show through dual-Fab cis binding that evolution has resulted in similar outcomes. The results presented here reveal an, up till now, unknown added value of using F(ab’)_2_ fragments rather than single Fabs when screening for functional antibodies.

The evolutionary mechanisms leading to dual-Fab cis binding antibodies may partly explain why children and adolescents suffer the most from recurring GAS infections (62). The time and exposures required to generate dual-Fab cis binding or other unorthodox antibodies could be related to the observation that frequent infections are needed to generate long-term immunity (23). We speculate that through multiple rounds of GAS infection during childhood, antibody hypermutation could generate dual-Fab cis binding antibodies, providing one potential mechanism to bypass the elaborate bacterial immune evasive mechanisms. It is difficult to predict how common this mode of binding is and how important it is in humans. Still, it is not unlikely that other bacterial proteins (such as adhesins or other molecules with long repetitive structures) might be targeted by dual-Fab binding antibodies (63).

Dual-Fab cis binding antibodies might have implications for antibody function and vaccine design. Our findings could explain why attempts at generating an effective vaccine against GAS have so far been unsuccessful (25). Even if a vaccine can induce high antibody titers, a large portion of these antibodies could be non-functional. If it turns out that dual-Fab cis binding antibodies are critical for function, antibody-based vaccine development strategies could be modified to take this into account. If vaccine peptides do not maintain conformational integrity (64) or span dual-Fab possible epitopes (10-15 nm separation), they might not be able to exhibit effective antibody-based immunity. Cell-mediated immunity would not be affected since T cell-MHC interaction, by definition, needs to be based on short peptides. The discovery and characterization of a putatively protective antibody against experimental GAS infection opens the possibility for monoclonal immunotherapy, but further validation of *in vivo* functions is required before that could be explored.

## Methods

See table in manuscript file for list of reagents.

### Single B cell purification, baiting and isolation

B cell isolation was performed as described previously (65), with some modifications. Briefly, 35 ml of blood was drawn (into citrated collection tubes) from a young woman who had recently recovered from a group A streptococcal infection. Collection and analysis of human blood samples were approved by the regional ethics committee, permit number 2015/801. The blood was treated with 2.5 μl/ml Rosettesep B (Stemcell technologies) for 20 mins at room temperature.

The blood was then diluted 1:1 in phosphate buffered saline (PBS) and layered onto Lymphoprep gradients. After centrifugation (30 mins at 800 x g), the plasma was collected and frozen while the B cell layer (around 7 ml) was removed, diluted with 43 ml of PBS, and centrifuged again. This washing step was repeated twice. The B cells were counted and kept at room temperature for staining (typical yields are 2-5 million cells per 30-40 ml of blood).

### B cell staining, baiting and sorting

The B cells were concentrated into a final volume of 500 μl in PBS. The cells were then blocked with 5% BSA for 20 minutes before being stained with antibodies against CD19-PE (BD-555413), CD3-BV510 (BD-564713), and IgG-BV421 (BD-562581). The B cells were also labelled with the Sytox-FITC live/dead stain (Thermofischer-S34860). Baiting of the B cells was done using soluble M1 protein isolated from an MC25 group A streptococcus M1 strain. The M1 protein isolation procedure was previously described elsewhere (66). The M1 protein was directly conjugated to Alexa Fluor 647 using the microscale labeling kit (Invitrogen). In addition to the antibodies and live/dead stains, 0.1 μg/ml of AF694-M1 was added to the cells and the mixture was incubated at 32°C for 20 minutes (M1 undergoes a conformational change at 4°C which could obscure important epitopes (67)). After the incubation, the cells were washed with PBS twice and were kept on ice until further analysis. The gates for sorting were set on a FACSAriaFusion sorter using unstained cells and FMO-1 samples. A total of 100 cells were sorted from 2.5 million B cells directly into 10 μl of water containing RNase inhibitor in 96-well plates and were immediately transferred to a -80 °C freezer. The cells at this point would have been lysed due to osmotic pressure and the RNA stabilized in solution.

### Reverse transcription, family identification and cloning

The cells previously frozen in plates were thawed on ice and RT-PCR was performed using the OneStep RT-PCR kit (Qiagen) protocol without modification. The primer sequences used in the PCR steps were taken directly from the Smith et al (2009) paper without any modifications. After the RT-PCR, the nested PCR was performed and the bands corresponding to the variable regions of the heavy and light chains were sequenced to identify the antibody families. Family-specific cloning primers were used to clone the variable chains into the plasmids containing the constant regions of the heavy and light chains. The expression plasmids were generously donated by Dr. Patrick Wilson’s group.

### General cell culture and transfection

THP-1 cells (Leukemic monocytes) were maintained in RPMI media supplemented with L-Glutamine and 10% FBS. The cells were kept at a cell density between 5-10×10^5^ cells per ml. THP-1-XBlue cells were maintained like regular THP-1 cells. HEK293 cells were maintained in DMEM supplemented with L-Glutamine and 10% FBS. The cells were never allowed to grow to 100% confluency. The day before transfection, 8×10^6^ cells were plated in circular 150 mm dishes. This transfection format allowed for the most efficient antibody recovery.

### Transfection, expression and purification

In total, 10 antibody construct pairs were successfully generated from 100 starting cells. The Antibody pairs were transformed into Mix’n’go E. coli. Transformant colonies were verified by sequencing and the plasmids were further propagated and DNA was extracted using a Zymoresearch midiprep kit. Plasmid pairs encoding full mature antibodies were co-transfected into HEK293 cells using the PEI transfection method (https://www.addgene.org/protocols/transfection/). Cells were briefly treated with 25 μM Chloroquine for 5 hours. Thereafter, 20 μg of heavy and light chain expression plasmid DNA were diluted in OptiMEM (Life technologies) media containing polyetheleneimine (PEI) at a 1:3 ratio (for 50 μg of DNA, 114 μl of a 1mg/ml PEI stock was used). The cells were incubated at 37°C for 18 hours before they were washed 2 times with PBS and the DMEM media was exchanged with OptiMEM. The cells were incubated for a further 72 hours before the supernatants were collected. The antibodies in the supernatants were purified using Protein G beads in a column setup. The antibodies were then titrated by comparing their concentrations on an SDS-PAGE to serial dilutions of a known concentration of Xolair (commercially bought Omalizumab, stored at 150 mg/ml).

### Bacterial strains, growth, and transformation

*Streptococcus pyogenes* strain SF370 (*emm*1 serotype), and AP1 (*emm*1 serotype) was grown in Todd-Hewitt Yeast media (THY) at 37C. The bacteria were maintained on agar plates for 3 weeks before being discarded. We chose to use SF370 in all of our experiments since it is an M1 serotype strain lacking protein H, which is a complicating factor (due to its strong Fc binding capacity and extensive homology with M protein (68)). The different M serotypes used in our cross-strain comparison were clinical isolates previously deposited into our in-house biobank. For experiments, overnight cultures were prepared in THY and were diluted at 1:20 on the day of the experiments. After dilution, three hours of growth at 37C ensured that the bacteria were in mid-log growth. For the generation of GFP-expressing strains, the SF370 and its ΔM isogenic counterpart, as well as an M5 (Manfredo), M12, M79, and M89 strains, were grown to mid-log before being washed with ice-cold water. The electrocompetent bacteria were electroporated with 20 μg of the pGFP1 plasmid and plated on Erythromycin supplemented THY plates. The successful transformants were fluorescent when examined under ultraviolet light. All strains were successfully transformed with the pGFP1 plasmid and yielded fluorescent colonies except for the M89 strain. Heat-killing the bacteria was done by growing the cultures to mid-log, washing them once in PBS, and incubating them on ice for 5 minutes. The bacteria were then heat-shocked at 80C for 5 minutes before being placed on ice for 15 minutes. For the phagocytosis assay, the heat-killed bacteria were centrifuged at 8000 x g for 3 minutes and resuspended in Na-medium ((5.6 mM glucose, 127 mM NaCl, 10.8 mM KCl, 2.4 mM KH_2_PO_4_, 1.6 mM MgSO_4_, 10 mM HEPES, 1.8 mM CaCl_2_; pH adjusted to 7.3 with NaOH). Under gentle rotation, heat-killed bacteria were stained with 5 μM Oregon Green 488-X succinimidyl ester (Thermofischer) at 37°C for 30 min. The bacteria were then centrifuged and resuspended in Sodium carbonate buffer (0.1 M, pH 9.0) for an additional staining step with the pH-sensitive dye CypHer5E (Fisher scientific). This was used at a concentration of 7 μg/ml in a volume of 1.5 ml for 2h at room temperature under gentle rotation, protected from light. The samples were washed once with Na-medium to remove excess dye and stored at 4C for later use.

### NGS of bacterial genomes

Whole-genome sequencing was done at the Center for Translational Genomics at Lund University. NextSeq 550 Illumina sequencing was used to sequence the bacterial genomes. The genome sequencing data were searched against the CDC database of M protein families to detect the target M protein sequence. Each M protein sequence was pairwise aligned with the target M1 protein using EMBOSS Needle web server, and the result is reported.

### Antibody screening and flow cytometry

For ELISAs: ELISA plates were coated overnight with 10 μg/ml M1 at 4°C, which had been purified from MC25 culture supernatants (66). After a 1-hour incubation at 37°C, the wells were washed 3 times with PBST and blocked with 2% BSA in 300 μl PBST for 30 minutes. After blocking, 300 μl of antibody containing supernatants were added to the wells, or diluted donor plasma as a control. The samples were incubated for 1 hour at 37°C, washed, and a solution of Protein G-HRP (diluted 1:3000) was added to the wells and incubated at 37°C for 1 hour. The samples were then washed and developed with 100 μl developing reagent (20 ml Substrate buffer NaCitrate pH 4.5 + 1 ml ABTS Peroxide substrate + 0.4 ml H2O2). Absorbance was read at OD450 following 5-30 minutes of color development at room temperature.

For flow cytometric screening: Overnights of SF370-GFP bacteria or its ΔM counterpart were diluted 1:20 into THY and grown until mid-log. 100 μl of the bacteria were distributed into wells of a 96 well plate. Antibodies purified from cell culture supernatants were diluted to 5 μg/ml and were digested with 1 μg/ml of IdeS for 3 hours at 37C. The digested antibodies were further diluted 1:10 into the bacterial suspension, reaching a final concentration of 0.5 μg/ml. The bacteria were incubated for 30 minutes at 37C before being washed twice with PBS. AF647-conjugated Fab α-Fab antibody fragments were used as secondary antibodies to detect binding of the primary α-M antibodies. After a 30-minute incubation with the Fab α-Fab fragments, the bacteria were washed and analyzed on a Cytoflex flow cytometer (Beck-man Coulter). The gates for the GFP-expressing bacteria were set using the SF370 parent strain (not expressing the GFP plasmid). GFP-expressing bacteria within the GFP-expressing gate were assessed for antibody staining (APC channel). Antibody staining reflects the presence of surface-bound primary-secondary antibody complexes and is indicative of bound anti-M antibodies.//

For western blotting: Antibody reactivity to linear epitopes was assessed by probing the lysates of SF370 and its ΔM mutant using western blotting. Briefly, pellets of logarithmically grown bacteria were incubated with phospholipase C for 30 minutes in PBS until the lysates became clear. The lysates were sonicated and cleared by centrifugation (15,000 x g for 3 minutes). We loaded 40 μg of 5 replicate sets of SF370 vs ΔM mutant protein on a gradient SDS-PAGE gel (4-20%). The gel electrophoresis was run for 60 minutes to achieve protein separation. The proteins were transferred from the gel to a PVDF membrane which was blocked for 45 minutes with 5% skimmed milk in PBST. The replicate lanes of the membrane were then cut and probed with 2 or 10 μg/ml of Xolair, Ab25, 32, 49 or IVIgG overnight at 4 °C. The membranes were washed 3 times with PBST and probed with the secondary HRP-conjugated goat anti-human IgG secondary (Rockland) antibody for 1 hour at room temperature. The secondary was later washed, and the membrane developed using a chemilumunescence reagent (WestFemto substrate, Thermofischer).

### Tissue microarray immunohistochemistry

Tissue microarrays (Novus biologicals, NBP2-78123) containing 90 tissue sections from human donors were deparaffinized and rehydrated before antigen retrieval (boiling in 10 mM Sodium Citrate buffer (pH 6.0)). The tissue sections were permeabilized and blocked before staining with 10 μg/ml of Xolair, Ab25, 32, 49, or 2.5% donor plasma. IVIG was used at 500 μg/ml, whereas anti-Troponin was used at 1:100 dilution to replicate a cardiac tissue-reactive antibody. The HRP-conjugated Rockland anti-human IgG antibody (609-103-123) was used as a secondary antibody at a 1:250 dilution.

### Agglutination assays

For agglutination assays: Overnight cultures of SF370 and its ΔM strain were diluted 1:5 in RPMI and were treated with 100 μg/ml of the anti-M antibodies or with 5% donor plasma. It is crucial for this series of experiments that the bacteria are incubated in a cuvette and are not shaken or vortexed during incubation. At indicated time points, the OD600 of the bacteria was measured, and at the 3.5-hour mark, the cuvettes were photographed. For aggregate dissolution experiments: SF370 bacteria were grown overnight, diluted 1:20 in THY, and left to grow for two hours. The bacteria were then supplemented with 100 μg/ml of the appropriate antibody. Two hours after inoculation, the bacteria were vortexed, imaged (randomly), and the aggregate areas were analyzed using Image J by segmenting the bacteria through automatic processing and then quantifying the pixel area of the identified bacterial regions.

### SIM imaging

Logarithmic phase bacteria were sonicated (VialTweeter; Hielscher) for 0.5 minutes to separate any aggregates and incubated fixed in 4% paraformaldehyde for 5 minutes on ice. The bacteria were thereafter washed with PBS twice (10,000 x g for 3 min). SF370 was stained with Alexa Fluor 647-conjugated wheat germ agglutinin (WGA). Bacteria were incubated with IdeS-cleaved Xolair, Ab25, Ab32, and Ab49 and stained with fluorescently labelled IgGFab or IgGFc specific F(ab’)_2_ fragments (DyLight488-conjugated anti-human IgGFc or IgGFab; Jackson ImmunoResearch Laboratory). Samples were mounted on glass slides using Prolong Gold Antifade Mountant with No. 1.5 coverslips. Images of single bacteria were acquired using an N-SIM microscope with LU-NV laser unit, CFI SR HP Apochromat TIRF 100X Oil objective (N.A. 1.49) and an additional 1.5x magnification. The camera used was ORCA-Flash 4.0 sCMOS camera (Hamamatsu Photonics K.K.) and the images were reconstructed with Nikon’s SIM software on NIS-Elements Ar (NIS-A 6D and N-SIM Analysis). Images of the bacteria were acquired with 488 and 640 nm lasers. For site localization, single bacteria were manually identified and imaged in time series with 50 frames. The analysis pipeline for site localization was implemented in Julia and is available on GitHub (44). A cut off of initial signal-to-noise ratio (SNR) was set to 0.3 and timeframes included were the ones with at least 70% of the initial SNR.

### Binding curves

SF370 bacteria were grown to mid-log, washed, and 10 ml of culture were concentrated into 1000 μl of PBS. The bacteria were stained with halving serial dilutions of the anti-M antibodies. 30 μl of bacteria were used per every 100 μl of IdeS treated antibody. The staining was performed at 4°C for 30 minutes (with shaking) before the bacteria were washed and stained with an excess of AF647-conjugated Fab anti-Fab fragments in a volume of 30 μl for 30 minutes at 4C with shaking. The bacteria were then diluted to 250 μl in PBS and analyzed by flow cytometry. The theoretical fit was done in MATLAB using fminsearch for an ideal binding curve with the dissociation constant as an unknown variable.

### Crosslinking of antibody F(ab’)_2_-fragments to the M1-protein

For the crosslinking of Ab25, Ab32 and Ab49 F(ab’)_2_ fragments to the M1 protein, we used two different preparations of the M1 protein; one expressed and purified as recombinant in E. coli as described for the B1B2B3 and C1C2C3 constructs above, and one purified from the culture supernatant of the S. pyogenes MC25 strain (66). The antibody F(ab’)_2_ fragments were cleaved and purified from the expressed intact antibodies using the FragIT-kit with Fc-capture columns (Genovis) according to the manufacturer’s instructions. For crosslinking, 25 μg of the recombinant M1 protein or 8 μg of the MC25 M1 protein were incubated with 5 μg of the respective F(ab’)_2_ fragments in 1× PBS pH 7.4 at 37 °C, 800 rpm, 30 min. Heavy/light disuccinimidylsuberate (DSS; DSS-H12/D12, Creative Molecules Inc.) resuspended in dimethylformamide (DMF) was added to final concentrations 250 and 500 μM and incubated for a further of 60 min at 37°C, 800 rpm. The crosslinking reaction was quenched with a final concentration of 50 mM ammonium bicarbonate at 37 °C, 800 rpm, 15 min.

### Sample preparation for MS

The crosslinked samples mixed with 8 M urea and 100 mM ammonium bicarbonate, and the cysteine bonds were reduced with 5 mM TCEP (37 °C for 2h, 800 rpm) and alkylated with 10 mM iodoacetamide (22 °C for 30 min, in the dark). The proteins were first digested with 1 μg of sequencing grade lysyl endopeptidase (Wako Chemicals) (37 °C, 800 rpm, 2h). The samples were diluted with 100 mM ammonium bicarbonate to a final urea concentration of 1.5 M, and 1 μg sequencing grade trypsin (Promega) was added for further protein digestion (37 °C, 800 rpm, 18 h). Samples were acidified (to a final pH 3.0) with 10% formic acid, and the peptides purified with C18 reverse phase spin columns according to the manufacturer’s instructions (Macrospin columns, Harvard Apparatus). Peptides were dried in a speedvac and reconstituted in 2% acetonitrile, 0.2% formic acid prior to mass spectrometric analyses.

### Liquid chromatography tandem mass spectrometry (LC-MS/MS)

All peptide analyses were performed on Q Exactive HF-X mass spectrometer (Thermo Scientific) connected to an EASY-nLC 1200 ultra-high-performance liquid chromatography system (Thermo Scientific). Peptides were loaded onto an Acclaim PepMap 100 (75μm x 2 cm) C18 (3 μm, 100 Å) pre-column and separated on an EASY-Spray column (Thermo Scientific; ID 75μm x 50 cm, column temperature 45°C) operated at a constant pressure of 800 bar. A linear gradient from 4 to 45% of 80% acetonitrile in aqueous 0.1% formic acid was run for 65 min at a flow rate of 350 nl min-1. One full MS scan (resolution 60000 @ 200 m/z; mass range 390–1 210m/z) was followed by MS/MS scans (resolution 15000 @ 200 m/z) of the 15 most abundant ion signals. The precursor ions were isolated with 2 m/z isolation width and fragmented using HCD at a normalized collision energy of 30. Charge state screening was enabled, and precursors with an unknown charge state and a charge state of 1 were rejected. The dynamic exclusion window was set to 10 s. The automatic gain control was set to 3e6 and 1e5 for MS and MS/MS with ion accumulation times of 110 ms and 60 ms, respectively. The intensity threshold for precursor ion selection was set to 1.7e4.

### Computational modeling

Several protocols of Rosetta software suit (69) were employed for macromolecular modeling of this study. To model the full-length antibodies, first the antigen-binding domains were characterized using Rosetta antibody protocol (70). Then, comparative models have been generated for both heavy and light chains using RosettaCM protocol (71) and aligned on the antigen-binding domains to represent the initial structure of the antibody. HSYMDOCK (72), and DaReUS_loop (73) web servers were used for symmetric docking of the Fc-domains and to model the hinge regions, respectively. Finally, 4K models were produced for each antibody as the final refinement and the top-scored models were selected based on XLs derived from mass spectrometry combined with Rosetta energy scores. Moreover, to characterize the M1 antibody interactions, TX-MS protocol were used (41), through which 2K docking models were generated and filtered out using distance constraints from DDA data. A final round of high-resolution modeling was performed on top models to repack the sidechains using RosettaDock protocol (74).

### Fluorescent Xolair competition experiments

Logarithmically grown SF370 bacteria were heat killed and labelled with Oregon Green (as described previously). The bacteria were mixed with antibodies or plasma/IVIG and incubated for 30 minutes at 37°C while shaking. Fluorescently conjugated Xolair (conjugated to Alexafluor 647 using the protein labeling kit (Invitrogen) according to the manufacturer’s instructions) was then added to the bacteria at a concentration of 100 μg/ml for an additional 30 minutes before being directly analyzed by flow cytometry. For experiments in which Fabs were used, the Fabs were generated using the Fabalactica digestion kit (Genovis) according to manufacturer’s instructions.

### Imaging-based binding assays

All images were acquired using an inverted Nikon Ti2-E widefield fluoresence microscope using a Plan Apo 20X objective (NA=0.75). Fluorescence was excited with the SPECTRA X light engine® (Lumencore inc, Beaverton, OR, USA) and collected with a Nikon DS-Qi2 CMOS controlled with NIS-Elements AR (v5.21.02). Multiple stage coordinates were automatically generated and imaged using NIS-Elements JOBs and a Nikon motorized stage. For image analysis, cells (anti-CD18, BV421) and bacteria (GFP) were segmented based on a background threshold algorithm for each channel separately. The cell masks were dilated to account for the whole cell area. Interaction between cells and bacteria was quantified by generating masks where cell and bacteria overlap, and the area and intensity profiles (mean, median, std) within this region were measured. Described image analysis was written in Julia (v1.6.0).

### Phagocytosis assay

The phagocytosis experiments were performed using persistent association normalization (39). Before opsonization, the CypHer5E-and Oregon Green-stained SF370 bacteria were sonicated for up to 5 min (VialTweeter; Hielscher) to disperse any large aggregates of bacteria. Sonication was deemed sufficient when clump dispersal was confirmed by microscopy. Staining and bacterial count (events/μl in the FITC +ve gate) was assessed by flow cytometry (CytoFLEX, Beckman-Coulter). The pH responsiveness of CypHer5E was tested by measuring the bacterial fluorescent staining in the APC channel before and after the addition of 1 μl of sodium acetate (3 M, pH 5.0) to 100 μl of the bacterial suspension. The presence of an acid-induced shift in fluorescence indicated successful staining. On the day of experiments, the appropriate number of bacteria were opsonized to suit each experiment. The opsonization with our M-specific antibodies, Xolair or IVIG, was performed at 37C for 30 minutes. For experiments with a variable MOP, serial dilutions of the opsonized bacteria were made and used to incubate with the THP-1 cells. By gating on the leukocyte population (Supp. Fig. 4A), specifically on single cells, we were able to group the cells into those associated with bacteria (FITC positive) and with internalized bacteria (FITC and APC positive). The panels in **Supp. Fig. 4A** show non-interacting cells compared to interacting cells as a result of phagocytosis at 4°C (or Cytochalasin D in **Supp. Fig. 4B**) and 37°C, respectively. In experiments where antibody concentration was the variable, serial dilutions of the antibodies were made in Na-medium in 96-well plates, and the bacteria were directly added to the antibodies for opsonization. THP-1 cells were washed in PBS on the day of the experiment and resuspended in Na-medium. The concentration of THP-1 cells was measured prior to phagocytosis by flow cytometry and adjusted to 2000 cells/μl (100 000 cells per well). The cells were then added to the 96-well plates previously prepared with varying concentrations of previously opsonized bacteria (MOP) or with different antibody concentrations. Finally, 50 μl of THP-1 cells were added on ice resulting in a final phagocytic volume of 150 μl. After a 5-minute incubation on ice, the plate was directly transferred to a shaking heating block set to 37°C while being protected from light or kept on ice as a control for internalization. Phagocytosis was stopped by putting the samples on ice for at least 15 min before data acquisition. Three experiments were performed to assess the association curves and four experiments were performed at MOP 400 to compare different antibodies.

For blood phagocytosis experiments, blood was drawn into heparinized tubes and was used immediately. Bacterial overnight cultures were diluted and grown to mid-log, washed with PBS, and resuspended in Na-medium. The bacteria were opsonized with 10 μg/ml of Xolair or Ab25 for 30 minutes at 37°C. After opsonization, the bacteria were sonicated for 1 minute and analyzed by flow cytometry for confirming live bacteria which were then used to inoculate the blood (which was diluted 10-fold in Na-medium). The infection was allowed to proceed for 30 minutes at 37°C in 96-well plates while shaking. The infection was then terminated by adding 1% paraformaldehyde for 30 minutes at room temperature. The cells were then washed with PBS and stained with SiR-Actin (emission at 674 nm) an anti-CD18 mAb (in BV421) which stains leukocytes. The cells were washed again and immobilized onto glass-bottom 96-well plates which were coated with an anti-CD29 antibody to capture cells. The wells were washed twice with PBS and imaged. More than 560 cells were imaged and their data quantified to generate the presented figures.

Flow cytometric acquisition was performed using a CytoFLEX (Beckman-Coulter) with 488 nm and 638 nm lasers and filters 525/40 FITC and 660/10 APC. Threshold was set at FSC-H 70,000 for phagocytosis and for bacteria FSC-H 2000 and SSC-H 2000. Gain was set to 3 for FITC and 265 for APC. Acquisition was set to capture at least 5 000 events of the target population with a velocity of 30 μl/min taking approximately 30 min to assess all samples. The 96-well plate was kept on an ice-cold insert throughout the data acquisition to inhibit further phagocytosis.

### NFkB activity luciferase assay

THP-XBlue-CD14 (Invivogen) cells were seeded at a density of 200,000 cells per well in 96 well plates. The cells were treated with the appropriate antibodies (at 0.5 μg/ml) with or without M1 protein (2 μg/ml) for 18 hours at 37 °C. After the incubation, 20 μl of the cell supernatant were aspirated and mixed with the developing reagent, as described by the assay instructions (QuantiBlue solution, Invivogen). The samples were incubated at 37 °C until development was appropriate and the OD650 measurement of the samples was done using a multi-well spectrophotometer.

### Animal model

The local Malmö/Lund Institutional Animal Care Care and Use Committee approved all animal use and procedures, ethical permit number 03681-2019. Nine-week-old female C57BL/6J mice (Scanbur/ Charles River Laboratories) were used. Monoclonal antibody Ab25 (0.4 mg/mouse), or intravenous immunoglobulin (10 mg/mouse) was administered intraperitoneally 6h pre-infection. *S. pyogenes* AP1 were grown to logarithmic phase in Todd–Hewitt broth (37°C, 5% CO2). Bacteria were washed and resuspended in sterile PBS. 10^6^ CFU of bacteria were injected subcutaneously into the scruff leading to systemic infection within 24 h. Mice were sacrificed 24 h post-infection, and organs (blood, livers, spleens, and kidneys) were harvested to determine the degree of bacterial dissemination (reported as CFU/g of tissue). The blood cell counts were analyzed by flow cytometry. Cytokines were quantified using a cytometric bead assay (CBA mouse inflammation kit, BD) according to manufacturer’s instructions.

## Author contributions

Conceptualization: WB, LB, OS, LM, JM and PN. Experimentation and data analysis: WB, LH, HK, VKA, TdN, SW, OA, EB, DT, and TH. Writing original draft: WB and PN. All authors contributed to reading and editing the final manuscript.

## Acknowledgements

WB, LH, OS, LB, LM, JM and PN is funded by the Knut and Alice Wallenberg Foundation. TH and equipment were funded by IngaBritt och Arne Lundbergs Forskningsstiftelse. HK is funded by Swiss National Science Foundation (grant no. P2ZHP3_191289). We thank Åsa Petersson for help with flow sorting, Dr. Berit Olofsson for antibody production and Gisela Hovold for technical assistance. We thank Center for Translational Genomics, Lund University and Clinical Genomics Lund, SciLifeLab for providing sequencing service. We thank Lund University Bioimaging Center (LBIC) for providing microscope services.

