## Supplemental Figures, Tables, and Data for "A human monoclonal antibody bivalently binding two different epitopes in streptococcal M protein protects against infection"

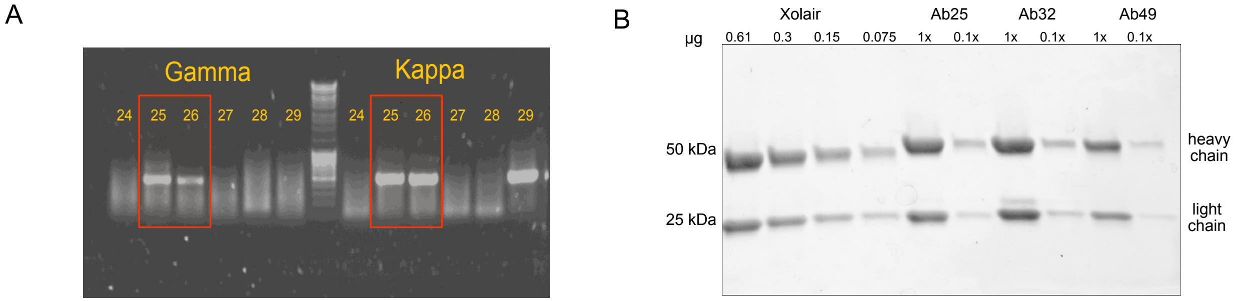


**Supplementary Figure 1. Single cell RT-PCR and antibody quality control**

**A** RT-PCR was performed to amplify the VDJ and VJ regions coding for heavy and light variable chains, respectively. Two successful PCR examples are Ab25 and 26, shown in the red boxes with both heavy and light chains yielding amplicons. **B** SDS-PAGE (4-20% gradient gel) analysis of Ab25, 32 and 49 compared to a serial dilution of Xolair. The gel was stained with Coomassie blue, destained and imaged.

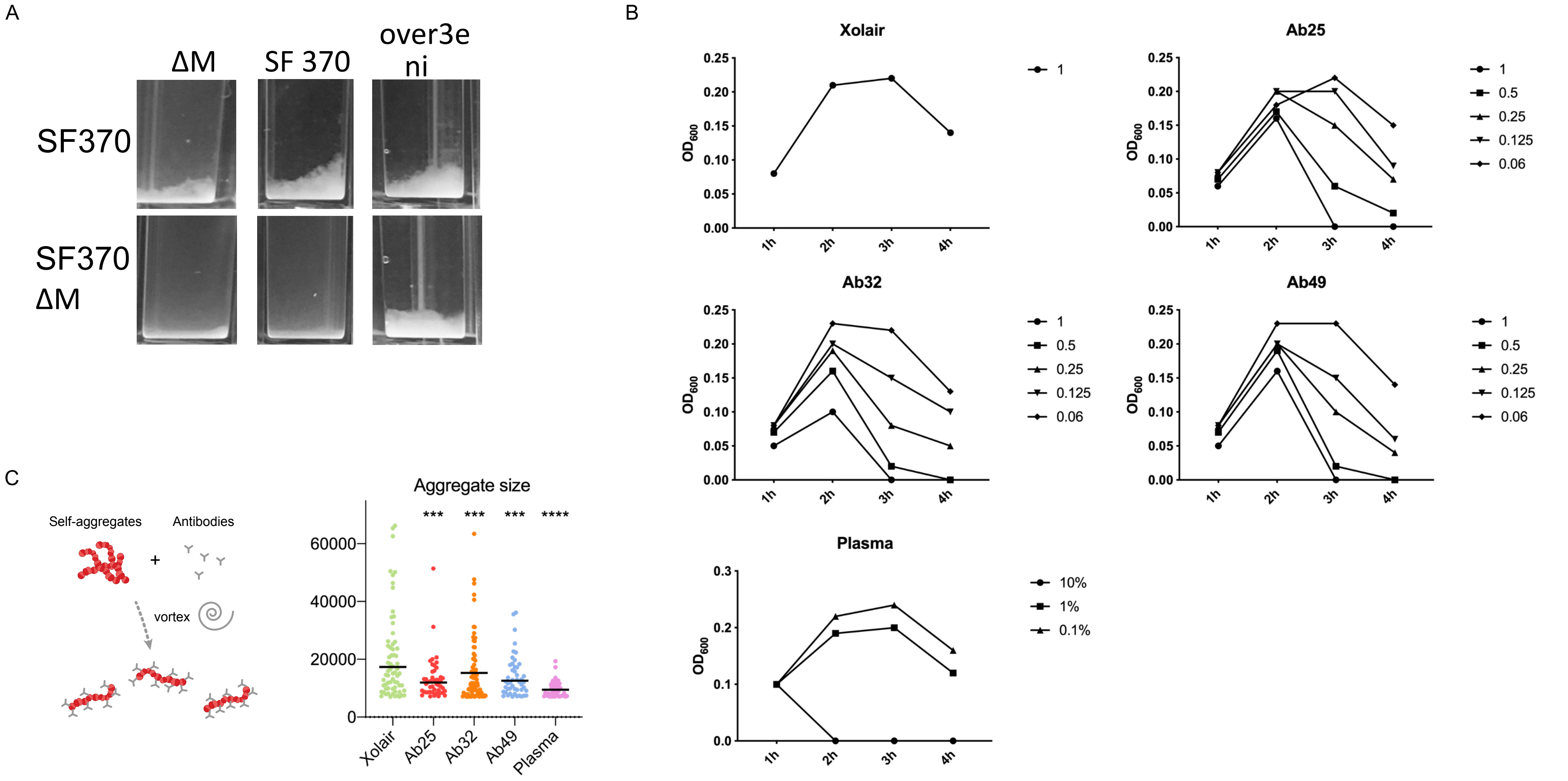


**Supplementary Figure 2. Agglutination and aggregate dissolution data.**

**A** SF370 overnight cultures were diluted 1:5 into THY and incubated with our antibodies (100 µg/ml) for 3 hours in plastic cuvettes at 37°C before measuring their $OD_{600}$. The agglutination effects of the antibody mix (Ab25,32 and 49 at 100 µg/ml each) were compared on WT and ΔM bacteria. Donor plasma was used as a positive control. **B** Dose-response experiment of individual antibodies in terms of agglutination, performed as in a with indicated antibody concentrations (µg/ml). **C** Bacteria treated with individual antibodies were left standing for 3 hours at 37°C before being vortexed. Aggregate size were quantified and tested using One-way ANOVA with multiple comparisons. The data in all the panels represents the results seen in at least 3 independent experiments. * denotes $p \leq 0.05$, ** for $p \leq 0.01$, *** for $p \leq 0.001$ and **** for $p \leq 0.0001$)

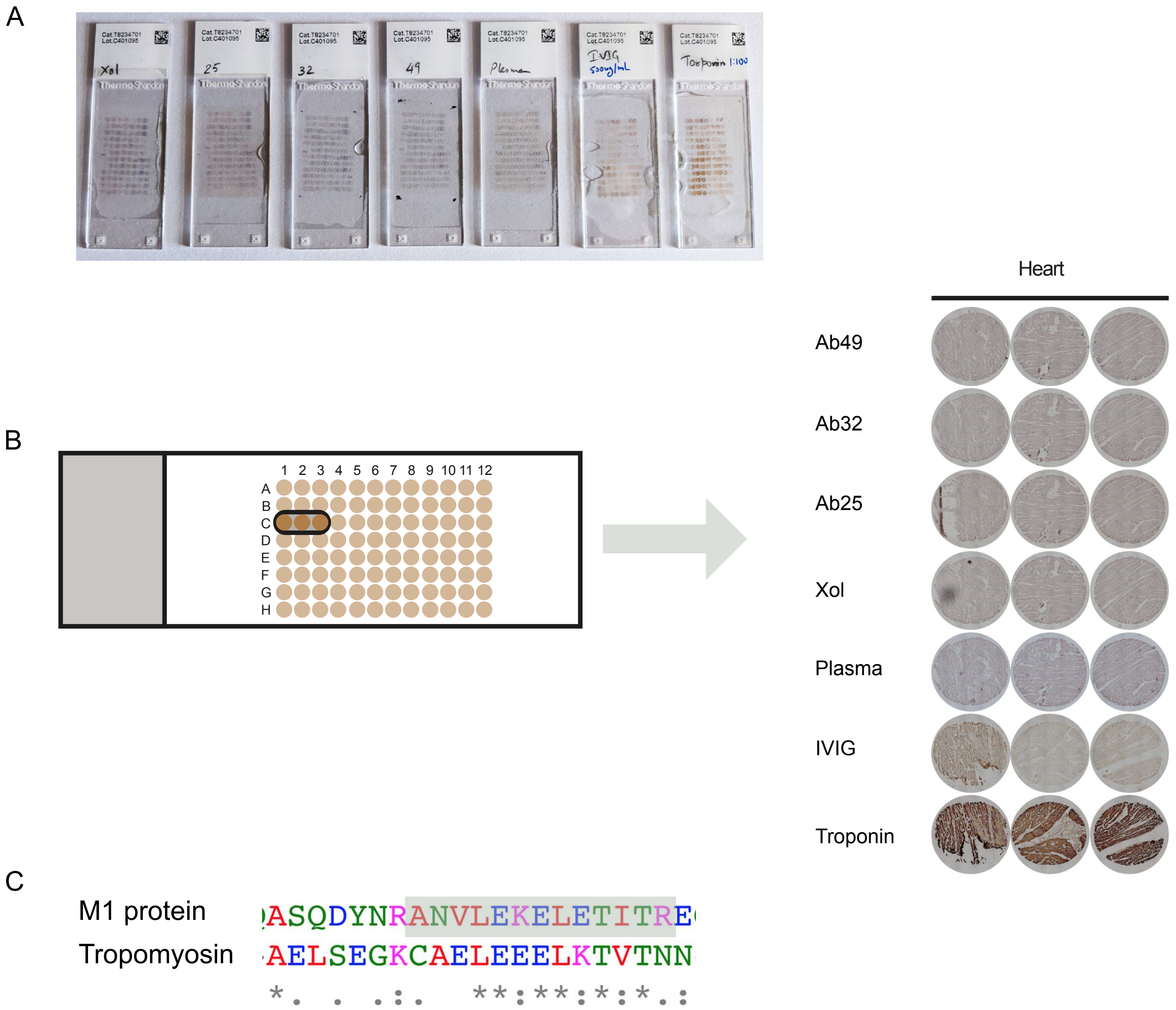


**Supplementary Figure 3. Tissue microarray analysis for anti-M antibodies.**

A) Tissue microarray stained with 10 µg/ml of Xolair, Ab25, 32, and 49, or 2.5% plasma, 500 µg/ml IVIG or 1:100 anti-Troponin antibody. Each spot is a sampling from different tissue. Tissue sections are 6 µm thick, 1.5 mm wide, and were mounted on positively charged glass slides. None of the tissues showed reactivity with the monoclonal samples, except the positive control anti-Troponin, which was reactive with cardiac tissue, as shown. Out of the 30 tissue types present on the slides, the cardiac samples would otherwise be the tissue type that had the largest risk of reactivity due to the M-protein mimicry with heart tissue. B) Zoomed in images of tissue spots representing cardiac tissue. C) Sequence alignment between M1 protein and tropomyosin. The highlighted area indicates identified the crosslinked peptide (closest to the interaction site) between Ab25 and Ab49.

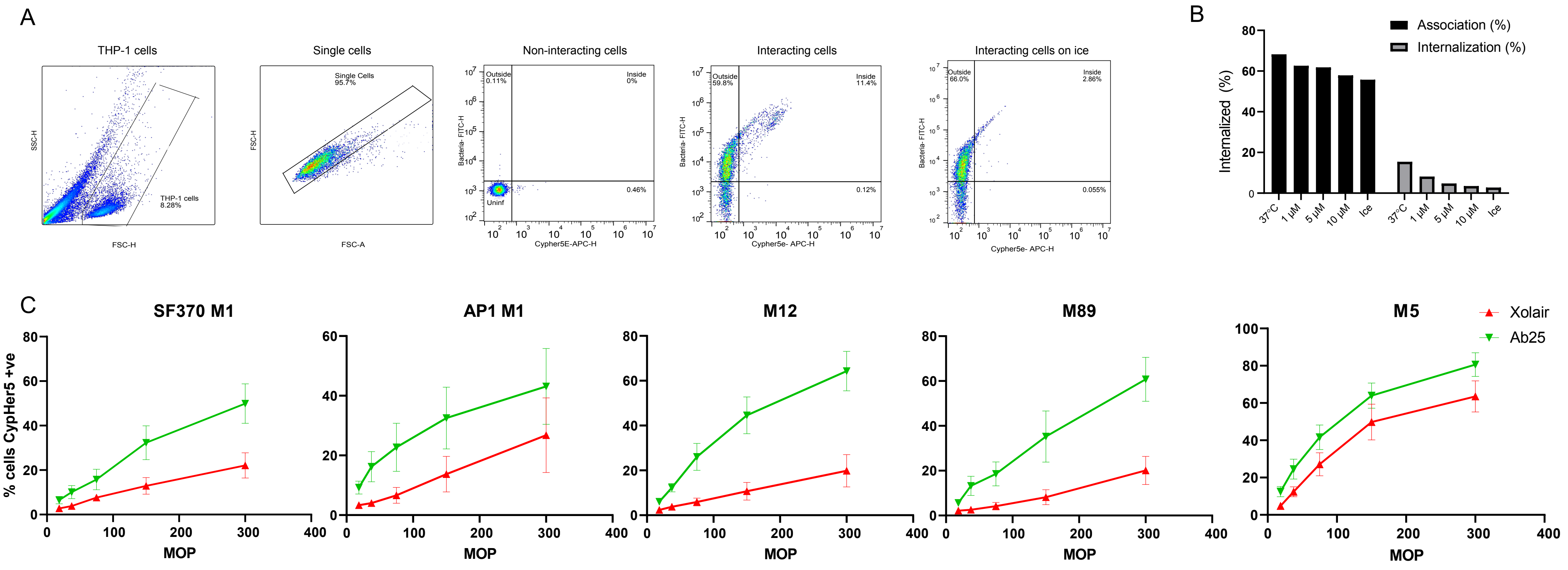


**Supplementary Figure 4. Cross-strain phagocytosis of Ab25 and gating strategy for phagocytosis experiments.**

**A.** From left to right: THP-1 cells are gated based on FSC and SSC. Single cells were selected based on FSC- Area and Height parameters. Non-interacting cells which have no FITC (bacteria) or CypHer5E (internalized bacteria) signal. Interacting cells associated with bacteria show a shift to the upper left quadrant (outside) whereas THP-1 cells with internalized bacteria shift to the upper right quadrant due to acquisition of the APC signal. THP-1 cells inoculated with bacteria were kept on ice to reduce the phagocytosis. This is visible as a reduction in the number of events in the upper right quadrant.

**B**. Fluorescent streptavidin beads previously opsonized with 1 mg/ml of IVIG beads were incubated with THP-1 cells as in (A). In addition to the experiment at 37°C, interactions were allowed to occur on ice or with 1,5, or 10 µM Cytochalasin D. The cells were then analyzed by flow cytometry and the % of cells which had internalized beads was displayed.

**C.** 10 µg/ml of Xolair or Ab25 were used to study phagocytic enhancement across different M serotypes. GAS with serotypes M1 (AP1), M12, M89 and M5 were compared in addition to the M1 SF370 strain. The effect of the antibody treatment on internalization rate was measured using flow cytometry. The error bars represent the standard error of the mean (SEM). The data is from three independent experiments.

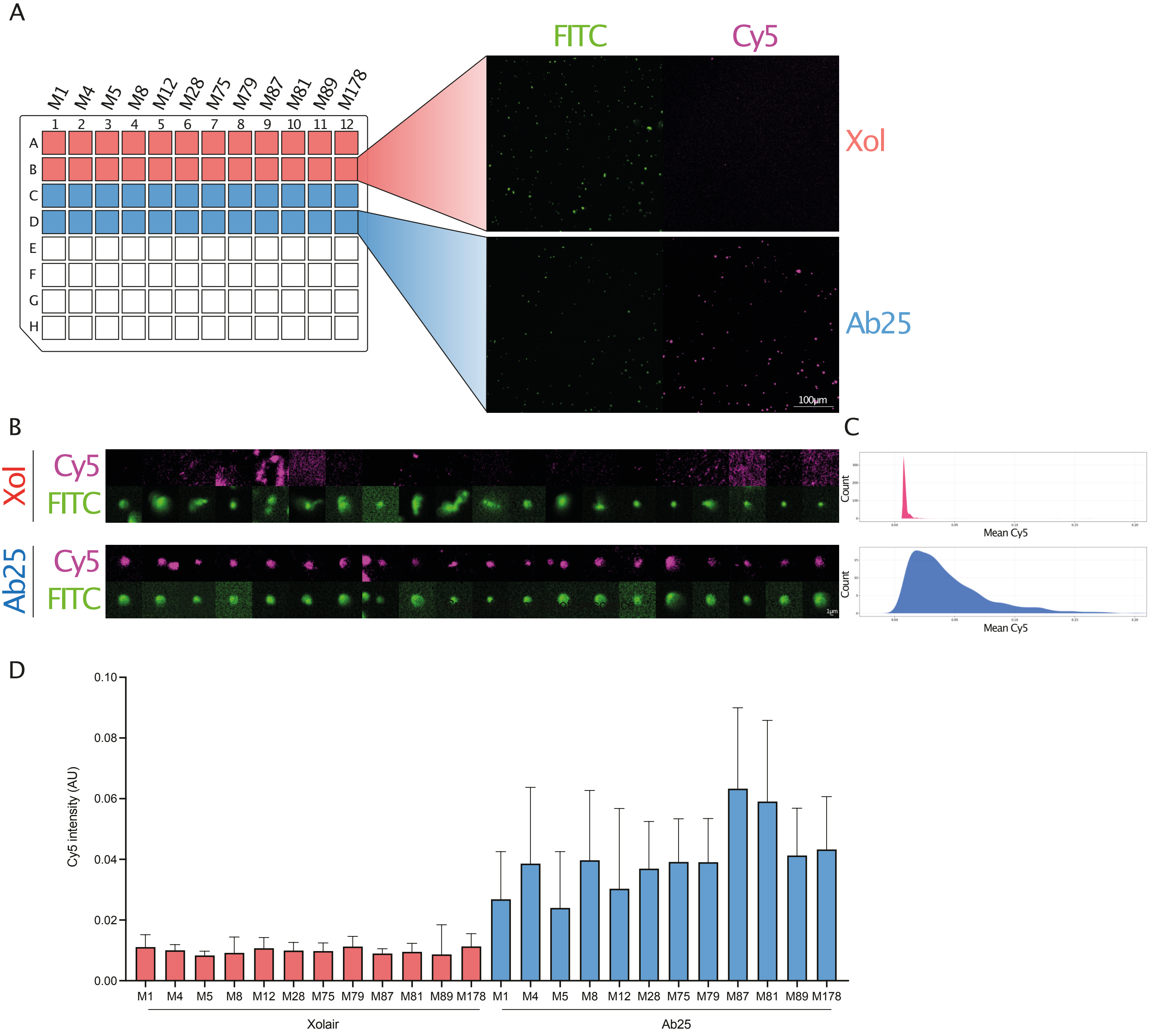


**Supplementary Figure 5. Cross-strain immunofluorescence.** Heat-killed bacteria were stained with Oregon Green (FITC), and incubated with 10 µg/ml Xolair or Ab25, followed by fluorescent secondary antibody detection (Cy5). A. Images were acquired on a wide-field epifluorescence microscope using a 20X objective (NA = 0.75). The acquisition of images was automated using Nikon JOBs by scanning through a 96-well plate acquiring two images/sample. FITC (bacteria) was segmented by a background segmentation algorithm (> median + 5 std of background signal). B, C. Individual bacteria were labeled and the overlapping signal in Cy5 was measured. Contrast adjustment is set to the same for all shown images. The images shown are from randomly selected bacteria corresponding to the median data population. As the bacteria were chosen by the software without human intervention, some events could be mildly out-of-focus, explaining the blurry appearance. D. Data is from two separate experiments and with a total of ~18000 bacteria analyzed.

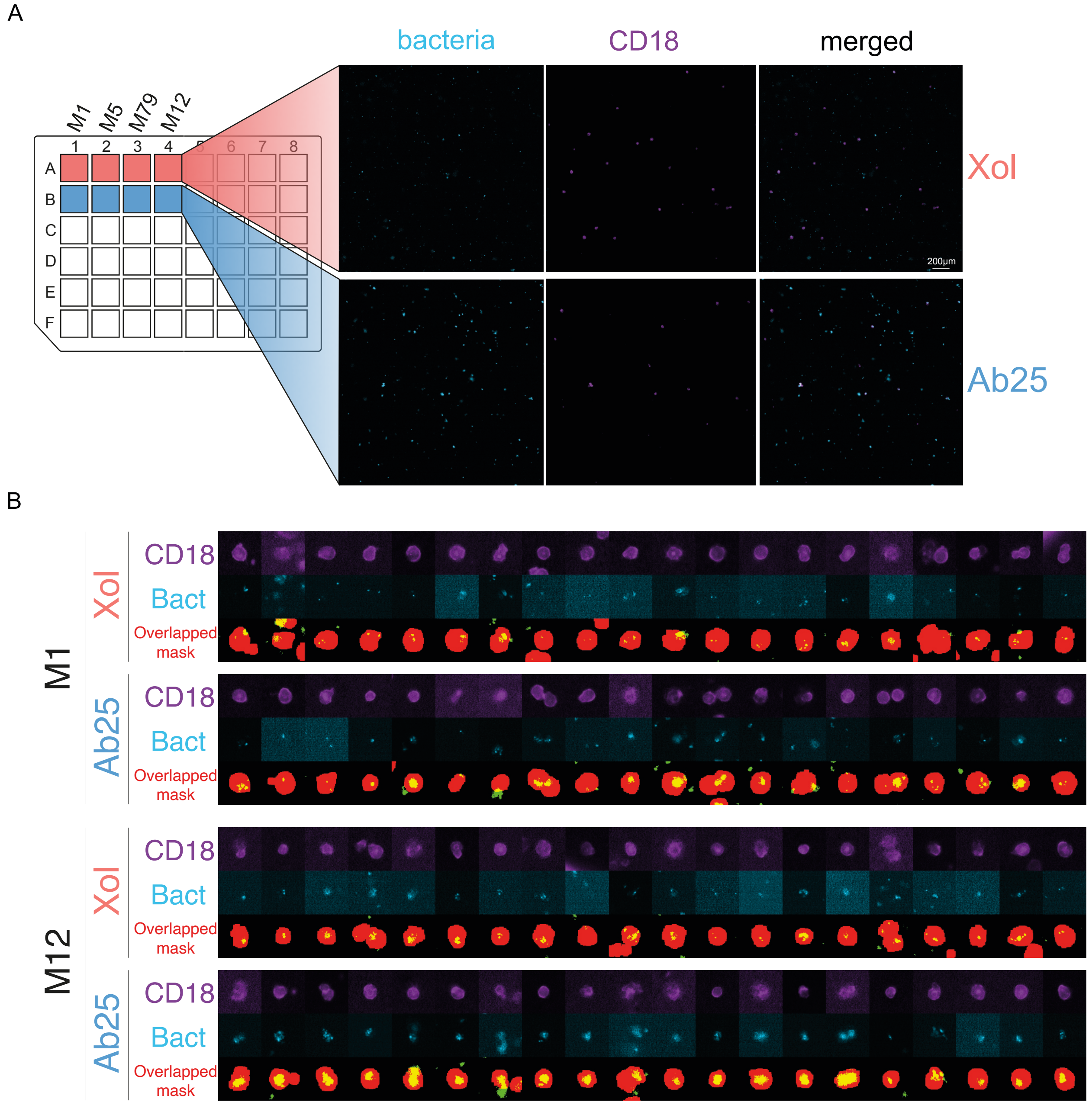


**Supplementary Figure 6. Imaging-based whole blood phagocytosis.**
Imaging-based whole blood phagocytosis. A) Whole blood was diluted and infected with 4 different strains of GAS expressing GFP (FITC). After 30 minutes, the infection was fixed with PFA and the cells were stained with anti-CD18 (BV421, shown in magenta). The blood was immobilized onto glass-bottom 96-well plates which were precoated with anti-CD29. Images were acquired on a widefield epifluorescence microscope using a 20X objective (NA = 0.75). The acquisition of images was automated using Nikon JOBs by scanning through a 96 well plate acquiring 25 images/sample. GFP (bacteria) and BV421 (CD18-positive leukocytes) were segmented based on a signal above 5 and 4 std of the background respectively. The overlapping mask for both channels was generated and the area of overlap was measured. Contrast adjustment is set to the same for all shown images. More than 560 cells were analyzed for this experiment B) Magnified images of cells representative cells were randomly selected and compared side-by-side to highlight the GFP area change with and without pre-opsonization. The presented images are chosen from a dataset representing the median of each population. The differences seen across the two groups, therefore, are fully representative of the data shown in Fig 3F.

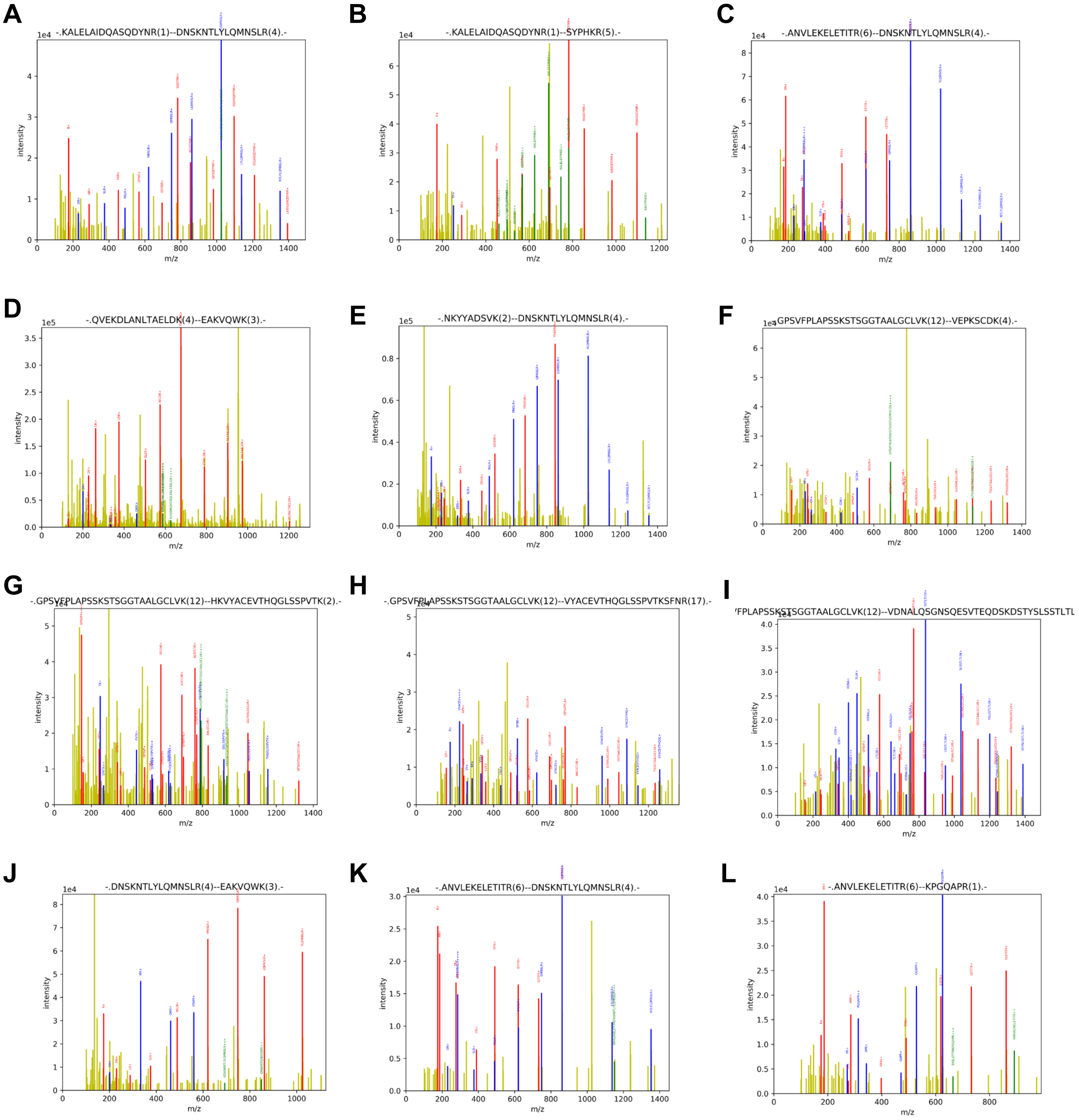


**Supplementary Fig 7. DDA peptide spectra**

Example DDA peptide spectra of cross-linked interactions identified (see also supplemental Table 1 for more details). One spectrum is shown for each cross-linked peptide pair. The red and the blue peptides represent parent peptides, respectively, whereas the green signal arises from the aforementioned peptides containing an additional DSS cross-linker arm.

| Cross-link no. | Peptide1 | Peptide2 | Protein1 | Protein2 | Cross-link distance (Å) | Spectrum no. in Fig. S4 |
|---|---|---|---|---|---|---|
| 1 | **K**ALELAIDQASQDYNR | DNS**K**NTLYLQMNSLR | M1-B2 repeat | AB25-heavy | 21.7 | A |
| 2 | **K**ALELAIDQASQDYNR | SYPH**K**R | M1-B2 repeat | AB25-heavy | 35.4 | B |
| 3 | ANVLE**K**ELETITR | DNS**K**NTLYLQMNSLR | M1-B3-S domain | AB25-heavy | 12.7 | C |
| 4 | QVE**K**DLANLTAELDK | EA**K**VQWK | M1-C1-C2 repeats | AB25-light | 22.5 | D |
| 5 | N**K**YYADSVK | DNS**K**NTLYLQMNSLR | AB25-heavy | AB25-heavy | 39.3 | E |
| 6 | GPSVFPLAPSS**K**STSGGTAALGCLVK | VEP**K**SCDK | AB25-heavy | AB25-heavy | 11.3 | F |
| 7 | GPSVFPLAPSS**K**STSGGTAALGCLVK | H**K**VYACEVTHQGLSSPVTK | AB25-heavy | AB25-light | 24.9 | G |
| 8 | GPSVFPLAPSS**K**STSGGTAALGCLVK | VYACEVTHQGLSSPVT**K**SFNR | AB25-heavy | AB25-light | 11.1 | H |
| 9 | GPSVFPLAPSS**K**STSGGTAALGCLVK | VDNALQSGNSQESVTEQDS**K**DSTYSLSSTLTLSK | AB25-heavy | AB25-light | 18.0 | I |
| 10 | DNS**K**NTLYLQMNSLR | EA**K**VQWK | AB25-heavy | AB25-light | 54.6 | J |
| 11 | ANVLE**K**ELETITR | DNS**K**NTLYLQMNSLR | M1-B3-S domain | AB49-heavy | 30.5 | K |
| 12 | ANVLE**K**ELETITR | **K**PGQAPR | M1-B3-S domain | AB49-light | 28.6 | L |

**Supplementary Table 1. Identified cross-linked peptides between M1, Ab25 and Ab49.** The cross-link number is indicated. For peptides 1 and 2, the cross-linked lysine residues are indicated in bold. Protein 1 and 2 indicate the parent proteins of the respective peptides. The cross-linked lysine residue distance is for each cross-link is given, and a reference to a corresponding spectrum in Supplementary Figure 4.

| emm type | emm pattern type (1) | emm cluster (2) | Upper epitope KALELAIDQASQDYNRANVLEKELETITR | Lower epitope QVEKDLANLTAELDK |
|---|---|---|---|---|
| *emm79* | E | E3 | No defined repeat | C repeat |
| *emm4* | E | E1 | No defined repeat | C repeat |
| *emm81* | D | E6 | No defined repeat | C repeat |
| *emm75* | E | E6 | No defined repeat | C repeat |
| *emm89* | E | E4 | No defined repeat | C repeat |
| *emm8* | E | E4 | No defined repeat | C repeat |
| *emm87* | E | E3 | No defined repeat | C repeat |
| *emm28* | E | E4 | No defined repeat | C repeat |
| *emm1* | A-C | A-C3 | B2-B3 repeat | C repeat |
| *emm179* | n.a. | Y (D) | No defined repeat | C repeat |
| *emm104* | E | E2 | No defined repeat | C repeat |
| *emm12* | A-C | A-C4 | B2-B3 repeat | C repeat |
| *emm5* | A-C | Y (A-C) | B2-B3 repeat | C repeat |

**Supplementary Table 2. Emm type analysis.** The M protein strains are listed according to Ab25 binding strength as reported in Fig. 1F. The emm pattern type, and emm cluster according to published reports by McMillan et al, and Sanderson-Smith et al. The majority belong to different E pattern type strains. All emm types have C repeats where the lower binding site of Ab25 is, whereas in most cases there are no defined repeats for the upper binding epitope. The *emm1* strain was used for the cross-linking experiment where the upper binding epitope (KALELAIDQASQDYNRANVLEKELETITR) of Ab25 was determined. In emm1, this epitope is placed in B2-B3 region. Ab49 also shares parts of this epitope (ANVLEKELETITR).


1. McMillan DJ, Drèze PA, Vu T, Bessen DE, Guglielmini J, Steer AC, et al. Updated model of group A *Streptococcus* M proteins based on a comprehensive worldwide study. Clin Microbiol Infect. 2013 May;19(5):E222-9.

2. Sanderson-Smith M, De Oliveira DMP, Guglielmini J, McMillan DJ, Vu T, Holien JK, et al. A systematic and functional classification of Streptococcus pyogenes that serves as a new tool for molecular typing and vaccine development. J Infect Dis. 2014 Oct 15;210(8):1325–38.

| | |
|---|---|
| M1-90-226-CTG | >gene_1147\|GeneMark.hmm\|306_aa\|+\|113293\|114210<br>>NODE_5_length_114211_cov_48.735082<br>MAKNNTNRHYSLRKLKTGTASVAVALTVLGAGFANQTEVKANGDGNPREVIEDLAANNPA<br>IQNIRLRHENKDLKARLENAMEVAGRDFKRAEELEKAKQALEDQRKDLETKLKELQQDYD<br>LAKESTSWDRQRLEKELEEKKEALELAIDQASRDYHRATALEKELEEKKKALELAIDQAS<br>QDYNRANVLEKELETITREQEINRNLLGNAKLELDQLSSEKEQLTIEKAKLEEEKQISDA<br>SRQSLRRDLDASREAKKQVEKDLANLTAELDKVKEEKQISDASRQGLRRDLDASREAKKQ<br>VEKDLA |
| M4-BS4-CTG | >gene_436\|GeneMark.hmm\|222_aa\|+\|185954\|186619<br>>NODE_2_length_186620_cov_148.285713<br>MARKDTNKQYSLRKLKTGTASVAVAVAVLGAGFANQTEVKAAEIKKPQADSAWNWPKEYN<br>ALLKENEELKVEREKYLSYADDKEKDPQYRALMGENQDLRKREGQYQDKIEELEKERKEK<br>QERQEQLERQYQIEADKHYQEQQKKHQQEQQQLEAEKQKLAKDKQISDASRQGLSRDLEA<br>SREAKKKVEADLAALTAEHQKLKEDKQISDASRQGLSRDLEA |
| M5-Manfredo-CTG | >gene_1398\|GeneMark.hmm\|194_aa\|+\|61232\|61813<br>>NODE_10_length_61815_cov_235.110363<br>MARENTNKHYSLRKLKKGTASVAVALSVLGAGLVVNTNEVSAAVTRGTINDPQRAKEALD<br>KYELENHDLKTKNEGLKTENEGLKTENEGLKTENEGLKTEKKEHEAENDKLKQQRDTLST<br>QKETLEREVQNTQYNNETLKIKNGDLTKELNKTRQELANKQQESKENEKALNELLEKTVK<br>DKIAKEQENKETIG |
| M8-BB3-CTG | >gene_1160\|GeneMark.hmm\|238_aa\|-\|2\|715<br>>NODE_7_length_101418_cov_178.048943<br>MARKDTNKQYSLRKLKTGTASVAVAVAVLGAGFANQTEVKAESPKSHSISNNEQLINELN<br>DLIEENNDLKDKLARNLDLLDNTREKDPQYRALMGENQDLREKEGKYQDKIKKLEEKEKN<br>LEKKSEDVERHYLKKLDQEHKEQQERQKNLEELERQSQREIDKRYQEQLQKQQQLETEKQ<br>ISEASRKSLSRDLEASRAAKKDLEAEHQKLKEEKQISDASRQGLSRDLEASREAKKKV |
| M12-BS3-CTG | >gene_1710\|GeneMark.hmm\|575_aa\|+\|13619\|15346<br>>NODE_21_length_15967_cov_54.040096<br>MAKNTTNRHYSLRKLKTGTASVAVALTVVGAGLVAGQTVRADHSDLVAEKQRLEDLGQKF<br>ERLKQRSELYLQQYYDNKSNGYKGDWYVQQLKMLNRDLEQAYNELSGEAHKDALGKLGID<br>NADLKAKITELEKSVEEKNDVLSQIKKELEEAEKDIQFGREVHAADLLRHKQEIAEKENV<br>ISKLNGELQPLKQKVDETDRNLQQEKQKVLSLEQQLAVTKENAKKDFELAALGHQLADKE<br>YNAKIAELESKLADAKKDFELAALGHQHAHNEYQAKLAEKDGQIKQLEEQKQILDASRKG<br>TARDLEAVRQAKKATEAELNNLKAELAKVTEQKQILDASRKGTARDLEAVRKAKAQVEAA<br>LKQLEEQNKISEASRKGLRRDLDASREAKKQVEKDLANLTAELDKVKEEKQISDASRQGL<br>RRDLDASREAKKQVEKALEEANSKLAALEKLNKELEESKKLTEKEKAELQAKLEAEAKAL<br>KEQLAKQAEELAKLRAGKASDSQTPDAKPGNKAVPGKGQAPQAGTKPNQNKAPMKETKRQ<br>LPSTGEAANPFFTAAALTVMATAGVAAVVKRKEEN |
| M28-BB5-CTG | >gene_950\|GeneMark.hmm\|358_aa\|-\|4735\|5811<br>>NODE_5_length_210162_cov_207.134746<br>MARKDTNKQYSLRKLKTGTASVAVAVAVLGAGFANQTEVKAAESPKSTETSANGADKLAD<br>AYNTLLTEHEKLRDEYYTLIDAKEEEPRYKALRGENQDLREKEGKYQDKIKKLEEKEKNL<br>EKKSEDVERHYLKKLDQEHKEQEERQKNLEELERQSQREIDKRYQEQLQKQQQLETEKQI<br>SEASRKSLSRDLEASRAAKKDLEAEHQKLKEEKQISDASRQGLSRDLEASREAKKKVEAD<br>LAEANSKLQALEKLNKELEEGKKLSEKEKAELQARLEAEAKALKEQLAKQAEELAKLKGN<br>QTPNAKVAPQANRSRSAMTQQKRTLPSTGEAANPFFTAAAATVMVSAGMLALKRKEEN |
| M75-BS7-CTG | >gene_561\|GeneMark.hmm\|387_aa\|-\|31358\|32521<br>>NODE_3_length_208685_cov_62.045909<br>MARKDTNKQYSLRKLKTGTASVAVAVAVLGAGFANQTEVKAEEERTFTELPYEARYKAWK<br>SENDELRENYRRTLDKFNTEQGKTTRLEEQNKKLHSELASVTETLTSVTEADDKKIKDLT<br>DRDKISSNLLGNAKDQINKLTTEKDTLAEKAKKLEEDKQISDASRKSLSRDLEASRAAKK<br>ELEANHQKLETEHKKLKEEKQISDASRQGLSRDLEASREAKKKVEADLAALTAEHQKLKE<br>EKQISDASRQGLSRDLEASREAKKKVEADLAEANSKLQALEKLNKELEEGKKLSEKEKAE<br>LQARLEAEAKALKEQLAKQAEELAKLKGNQTPNAKVAPQANRSRSAMTQQKRTLPSTGEA<br>ANPFFTAAAATVMVSAGMLALKRKEEN |

| | |
|---|---|
| M79-BTS4-CTG | >gene_435\|GeneMark.hmm\|222_aa\|+\|185954\|186619<br>>NODE_2_length_186620_cov_58.625015<br>MARKDTNKQYSLRKLKTGTASVAVAVAVLGAGFANQTEVKAAEIKKPQADSAWNWPKEYN<br>ALLKENEELKVEREKYLSYADDKEKDPQYRALMGENQDLRKREGQYQDKIEELEKERKEK<br>QERQEQLERQYQIEADKHYQEQQKKHQQEQQQLEAEKQKLAKDKQISDASRQGLSRDLEA<br>SREAKKKVEADLAALTAEHQKLKEDKQISDASRQGLSRDLEA |
| M81-BB6-CTG | >gene_94\|GeneMark.hmm\|396_aa\|-\|101371\|102561<br>>NODE_1_length_190029_cov_107.359270<br>MVRKDTNKQYSLRKLKTGTASVAVAVAVLGAGFANQTEVKAAGSEENVPKQQYNALWEEN<br>EDLRGRERKYIAKLEKEEIQNGELNEKNRKLEADIADLQDVIEDNDQEIKRKDRMYEAFL<br>KQSKDQVKDLTAEKDTLAEKAKKLEEDKQISDASRKSLSRDLEGSRAAKKELEAKHQKLE<br>TEHQKLKEDKQISDASRQGLSRDLEASREAKKKVEADLAALTAEHQKLKEEKQISDASRQ<br>GLSRDLEASREAKKKVEADLAEANSKLQALEKLNKELEEGKKLSEKEKAELQARLEAEAK<br>ALKEQLAKQAEELAKLRAGKASDSQTPDAKPGNKVVPGKGQAPQAGTKPNQNKAPMKETK<br>RQLPSTGEAANPFFTAAAATVMVSAGMLALKRKEEN |
| M85-SP5-CTG | >gene_158\|GeneMark.hmm\|402_aa\|-\|835\|2043<br>>NODE_2_length_119743_cov_614.138920<br>MVRKDTNKQYSLRKLKTGTASVAVAVAVLGAGFANQTEVKAAEQRAPQAKGTSVSADLYN<br>SLWDENKTLREKQGEYITKIQNEETKNKELDKKNKELDSRVTDLIDVIEHDDQELERKER<br>MYEAFLKQSKDQVNNLTAEKDTLAEKAKKLEEDKQISDASRKSLSRDLEGSRAAKKELEA<br>KHQKLETEHQKLKEDKQISDASRQGLSRDLEASREAKKKVEADLAALTAEHQKLKEEKQI<br>SDASRQGLSRDLEASREAKKKVEADLAEANSKLQALEKLNKELEEGKKLSEKEKAELQAR<br>LEAEAKALKEQLAKQAEELAKLRAGKASDSQTPDAKPGNKVVPGKGQAPQAGTKPNQNKA<br>PMKETKRQLPSTGEAANPFFTAAAATVMVSAGMLALKRKEEN |
| M87-BB10-CTG | >gene_1714\|GeneMark.hmm\|238_aa\|-\|3\|716<br>>NODE_26_length_15865_cov_62.755661<br>MARKDTNKQYSLRKLKTGTASVAVAVAVLGAGFANQTEVKAESPREVTNELAASVWKKKV<br>EEAKEKASKLEKQLEEAQKDYSEIEGKLEQFWHDYDKLEKENKEYASQLGKNQEEREKLE<br>LEYLRKSDEEYKEHQQYRQEQEERQKNLEELERQNKREIDKRYQEQLQKQQQLETEKQIS<br>EASRKSLSRDLEASRAAKKELEAEHQKLKEEKQISDASRKSLSRDLEASREAKKKVEA |
| M89-BSA4-CTG | >gene_1171\|GeneMark.hmm\|244_aa\|-\|2\|733<br>>NODE_8_length_78604_cov_108.090657<br>MARKDTNKQYSLRKLKTGTASVAVAVAVLGAGFANQTTVKADSDNINRSVSVKDNEKELH<br>NKIADLEEERGEHLDKIDELKEELKAKEKSSENVERHYLRKLDQEYKEQQERQKNLEELE<br>RQSQREVEKRYQEQLQKQQQLETEKQISEASRKSLSRDLEASRAAKKDLEAEHQKLKEEK<br>QISDASRQGLSRDLEASREAKKKVEADLAALTAEHQKLKEEKQISDASRQGLSRDLEASR<br>EAKK |
| M118-SP1-CTG | >gene_1703\|GeneMark.hmm\|332_aa\|-\|4820\|5818<br>>NODE_14_length_20826_cov_45.853546<br>MARKDTNKQYSLRKLKTGTASVAVAVAVLGAGFANQTEVKAAEKKVEVADSNASSVAKLY<br>NQIADLTDKNGEYLERIEELEERQKNLEKLERQSQVAADKHYQEQVKKHQEYKQEQEERQ<br>KNLEELERQNKREIDKRYQEQLQKQQQLETEKQISEASRKSLSRDLEASRAAKKDLEAEH<br>QKLKEEKQISDASRQGLSRDLEASREAKKKVEADLAEANSKLQALEKLNKELEEGKKLSE<br>KEKAELQAKLEAEAKALKEQLAKQAEELAKLKGNQTPNAKVAPQANRSRSAMTQQKRTLP<br>STGETANPFFTAAAATVMVSAGMLALKRKEEN |
| M179-BB12-CTG | >gene_92\|GeneMark.hmm\|433_aa\|-\|100604\|101905<br>>NODE_1_length_264627_cov_344.813212<br>MVRKDTNRHYSLRKLKTGTASVAVALSVLGAGLAVNQTEVSAKSVTRSTAQDPDKSRQAI<br>TEYEVENHKLTQEKNALTNRNQELTDENGELKTANEALRQRGDTLFNQRVKLEKQVQEKE<br>HNNKTLKIENGELKTENGDLTKKLDETRQELANKQQESKENEKTLNELLEKTVKDKIAKE<br>QENKETIGTLKKLLDETVKDKIAKEQKSKQDFGALKQELAKKEEQNKISDASRQGLRRDL<br>NASREAKKQVEKDLANLTAELDKVKEEKQVSDASRQGLRRDLDASREAKKQVEKALEEAN<br>SKLAALEKLNKELEESKKLTEKEKAELQAKLEAEAKALKEKLAKQAEELAKLRAGKASDS<br>QTPDAKPGNKAVPGKGQAPQAGTKPNQNKAPMKETKRQLPSTGEAANPFFTAAALTVMAT<br>AGVAAVVKRKKEN |

### Sequence similarity and alignment analysis

The genome sequencing data were searched against the CDC database of M protein families to detect the target M protein sequence. Each M protein sequence was pairwise aligned with the target M1 protein using EMBOSS Needle web server and the result is reported below.

```
########################################
# Program: needle
# Commandline: needle
#    -auto
#    -stdout
#    -asequence emboss_needle-I20210916-185110-0458-22637843-p1m.asequence
#    -bsequence emboss_needle-I20210916-185110-0458-22637843-p1m.bsequence
#    -datafile EBLOSUM62
#    -gapopen 10.0
#    -gapextend 0.5
#    -endopen 10.0
#    -endextend 0.5
#    -aformat3 pair
#    -sprotein1
#    -sprotein2
# Align_format: pair
#=======================================
# Aligned_sequences: 2
# 1: Reference_M1_sequence
# 2: M4_sequence
# Matrix: EBLOSUM62
# Gap_penalty: 10.0
# Extend_penalty: 0.5
# Length: 486
# Identity:     125/486 (25.7%)
# Similarity:   157/486 (32.3%)
# Gaps:         266/486 (54.7%)
# Score: 461.0
#=======================================

Reference_M1_      1 MAKNNTNRHYSLRKLKTGTASVAVALTVLGAGFANQTEVKANGDGNPREV     50
                     ||:.:||:.||||||||||||||||:.||||||||||||||.....|:
M4_sequence        1 MARKDTNKQYSLRKLKTGTASVAVAVAVLGAGFANQTEVKAAEIKKPQ--     48

Reference_M1_     51 IEDLAANNPAIQNIRLRYENKDLKARLENAMEVAGRDFKRAEELEKAKQ-     99
                      .|.|.|.|...|..|: ||::||...|..:..       |::.||..|
M4_sequence       49 -ADSAWNWPKEYNALLK-ENEELKVEREKYLSY-------ADDKEKDPQY     89

Reference_M1_    100 -ALEDQRKDLETKLKELQQDYDLAKESTSWDRQRLEKELEEKKEALELAI    148
                      ||..:.:||..:..:.|...           :.||||.:||:|..|
M4_sequence       90 RALMGENQDLRKREGQYQDKI-----------EELEKERKEKQERQE---    125

Reference_M1_    149 DQASRDYHRATALEKELEEKKKALELAIDQASQDYNRANVLEKELETITR    198
                      |..|.|.  ...:|..:|::|                           :
M4_sequence      126 -QLERQYQ--IEADKHYQEQQK---------------------------K    145

Reference_M1_    199 EQEINRNLLGNAKLELDQLSSEKEQLTIEKAKLEEEKQISDASRQSLRRD    248
                     .|:                  |::||..||.||.::|||||||||.|.||
M4_sequence      146 HQQ------------------EQQQLEAEKQKLAKDKQISDASRQGLSRD    177

Reference_M1_    249 LDASREAKKQVEKDLANLTAELDKVKEDKQISDASRQGLRRDLDASREAK    298
                     |:|||||||:||.|||.||||..|:||||||||||||||.|||:|
M4_sequence      178 LEASREAKKKVEADLAALTAEHQKLKEDKQISDASRQGLSRDLEA-----    222

Reference_M1_    299 KQVEKDLANLTAELDKVKEEKQISDASRQGLRRDLDASREAKKQVEKALE    348

M4_sequence      223 --------------------------------------------------    222

Reference_M1_    349 EANSKLAALEKLNKELEESKKLTEKEKAELQAKLEAEAKALKEQLAKQAE    398

M4_sequence      223 --------------------------------------------------    222

Reference_M1_    399 ELAKLRAGKASDSQTPDTKPGNKAVPGKGQAPQAGTKPNQNKAPMKETKR    448

M4_sequence      223 --------------------------------------------------    222

Reference_M1_    449 QLPSTGETANPFFTAAALTVMATAGVAAVVKRKEEN    484

M4_sequence      223 ------------------------------------    222


#---------------------------------------
#---------------------------------------
```

```
########################################
# Program: needle
# Commandline: needle
#    -auto
#    -stdout
#    -asequence emboss_needle-I20210916-185742-0233-9205053-p2m.asequence
#    -bsequence emboss_needle-I20210916-185742-0233-9205053-p2m.bsequence
#    -datafile EBLOSUM62
#    -gapopen 10.0
#    -gapextend 0.5
#    -endopen 10.0
#    -endextend 0.5
#    -aformat3 pair
#    -sprotein1
#    -sprotein2
# Align_format: pair
# Report_file: stdout
########################################

#=======================================
# Aligned_sequences: 2
# 1: Reference_M1_sequence
# 2: M5_sequence
# Matrix: EBLOSUM62
# Gap_penalty: 10.0
# Extend_penalty: 0.5
#
# Length: 495
# Identity:      79/495 (16.0%)
# Similarity:   110/495 (22.2%)
# Gaps:         312/495 (63.0%)
# Score: 225.5
#=======================================

Reference_M1_      1 MAKNNTNRHYSLRKLKTGTASVAVALTVLGAGF-ANQTEVKANGDGNPRE     49
                     ||:.|||:||||||||.|||||||||:|||||. .|..||.|   ...|.
M5_sequence        1 MARENTNKHYSLRKLKKGTASVAVALSVLGAGLVVNTNEVSA---AVTRG     47

Reference_M1_     50 VIEDLAANNPAIQNIRLRYENKDLKARLENAMEVAGRDFKRAEELEKAKQ     99
                     .|.|......|:....|  ||.|||.:.|...       ...|.|:...:
M5_sequence       48 TINDPQRAKEALDKYEL--ENHDLKTKNEGLK-------TENEGLKTENE     88

Reference_M1_    100 ALEDQRKDLETKLKELQQDYDLAKESTSWDRQRLEKELEEKKEALELAID    149
                     .|:.:.:.|:|:.||.:.:.|..|:      ||  ..|..:||.||..:.
M5_sequence       89 GLKTENEGLKTEKKEHEAENDKLKQ------QR--DTLSTQKETLEREVQ    130

Reference_M1_    150 QASRDYHRATA------LEKELEEKKKALELA-IDQASQDYNRA--NVLE    190
                       :..|:..|.      |.|||.:.::  ||| ..|.|::..:|  .:||
M5_sequence      131 --NTQYNNETLKIKNGDLTKELNKTRQ--ELANKQQESKENEKALNELLE    176

Reference_M1_    191 KEL-ETITREQEINRNLLGNAKLELDQLSSEKEQLTIEKAKLEEEKQISD    239
                     |.: :.|.:||| |:..:|
M5_sequence      177 KTVKDKIAKEQE-NKETIG-------------------------------    194

Reference_M1_    240 ASRQSLRRDLDASREAKKQVEKDLANLTAELDKVKEDKQISDASRQGLRR    289

M5_sequence      195 --------------------------------------------------    194

Reference_M1_    290 DLDASREAKKQVEKDLANLTAELDKVKEEKQISDASRQGLRRDLDASREA    339

M5_sequence      195 --------------------------------------------------    194

Reference_M1_    340 KKQVEKALEEANSKLAALEKLNKELEESKKLTEKEKAELQAKLEAEAKAL    389

M5_sequence      195 --------------------------------------------------    194

Reference_M1_    390 KEQLAKQAEELAKLRAGKASDSQTPDTKPGNKAVPGKGQAPQAGTKPNQN    439

M5_sequence      195 --------------------------------------------------    194

Reference_M1_    440 KAPMKETKRQLPSTGETANPFFTAAALTVMATAGVAAVVKRKEEN    484

M5_sequence      195 ---------------------------------------------    194


#---------------------------------------
#---------------------------------------
```

```
########################################
# Program: needle
# Commandline: needle
#    -auto
#    -stdout
#    -asequence emboss_needle-I20210916-185905-0776-44155798-p1m.asequence
#    -bsequence emboss_needle-I20210916-185905-0776-44155798-p1m.bsequence
#    -datafile EBLOSUM62
#    -gapopen 10.0
#    -gapextend 0.5
#    -endopen 10.0
#    -endextend 0.5
#    -aformat3 pair
#    -sprotein1
#    -sprotein2
# Align_format: pair
# Report_file: stdout
########################################

#=======================================
# Aligned_sequences: 2
# 1: Reference_M1_sequence
# 2: M8_sequence
# Matrix: EBLOSUM62
# Gap_penalty: 10.0
# Extend_penalty: 0.5
#
# Length: 490
# Identity:     124/490 (25.3%)
# Similarity:   165/490 (33.7%)
# Gaps:         258/490 (52.7%)
# Score: 449.5
#=======================================

Reference_M1_      1 MAKNNTNRHYSLRKLKTGTASVAVALTVLGAGFANQTEVKANGD-----G     45
                     ||:.:||:.||||||||||||||||:.||||||||||||||...     .
M8_sequence        1 MARKDTNKQYSLRKLKTGTASVAVAVAVLGAGFANQTEVKAESPKSHSIS     50

Reference_M1_     46 NPREVIEDLAANNPAIQNIRLRYENKDLKARLENAMEVAGRDFKRAEELE     95
                     |..::|.:|   |..|:      ||.|||.:|...:::    .....|.:
M8_sequence       51 NNEQLINEL---NDLIE------ENNDLKDKLARNLDL----LDNTREKD     87

Reference_M1_     96 KAKQALEDQRKDLETKLKELQQDYDLAKESTSWDRQRLEKELEEKKEALE    145
                     ...:||..:.:||..|..:.|...               |:||||::.||
M8_sequence       88 PQYRALMGENQDLREKEGKYQDKI---------------KKLEEKEKNLE    122

Reference_M1_    146 LAIDQASRDYHRATALE-KELEEKKKALELAIDQASQDYNRANVLEKELE    194
                     ...:...|.|.:....| ||.:|::|.||                  |||
M8_sequence      123 KKSEDVERHYLKKLDQEHKEQQERQKNLE------------------ELE    154

Reference_M1_    195 TITREQEINRNLLGNAKLELDQLSSEKEQLTIEKAKLEEEKQISDASRQS    244
                     .             .::.|:|:...|:.|   ::.:||.|||||:|||:|
M8_sequence      155 R-------------QSQREIDKRYQEQLQ---KQQQLETEKQISEASRKS    188

Reference_M1_    245 LRRDLDASREAKKQVEKDLANLTAELDKVKEDKQISDASRQGLRRDLDAS    294
                     |.|||:|||.|||.:|       ||..|:||:|||||||||||.|||:||
M8_sequence      189 LSRDLEASRAAKKDLE-------AEHQKLKEEKQISDASRQGLSRDLEAS    231

Reference_M1_    295 REAKKQVEKDLANLTAELDKVKEEKQISDASRQGLRRDLDASREAKKQVE    344
                     |||||:|
M8_sequence      232 REAKKKV-------------------------------------------    238

Reference_M1_    345 KALEEANSKLAALEKLNKELEESKKLTEKEKAELQAKLEAEAKALKEQLA    394

M8_sequence      239 --------------------------------------------------    238

Reference_M1_    395 KQAEELAKLRAGKASDSQTPDTKPGNKAVPGKGQAPQAGTKPNQNKAPMK    444

M8_sequence      239 --------------------------------------------------    238

Reference_M1_    445 ETKRQLPSTGETANPFFTAAALTVMATAGVAAVVKRKEEN    484

M8_sequence      239 ----------------------------------------    238


#---------------------------------------
#---------------------------------------
```

```
########################################
# Program: needle
# Commandline: needle
#    -auto
#    -stdout
#    -asequence emboss_needle-I20210916-190019-0322-98296274-p2m.asequence
#    -bsequence emboss_needle-I20210916-190019-0322-98296274-p2m.bsequence
#    -datafile EBLOSUM62
#    -gapopen 10.0
#    -gapextend 0.5
#    -endopen 10.0
#    -endextend 0.5
#    -aformat3 pair
#    -sprotein1
#    -sprotein2
# Align_format: pair
# Report_file: stdout
########################################
#=======================================
# Aligned_sequences: 2
# 1: Reference_M1_sequence
# 2: M12_sequence
# Matrix: EBLOSUM62
# Gap_penalty: 10.0
# Extend_penalty: 0.5
#
# Length: 587
# Identity:     318/587 (54.2%)
# Similarity:   371/587 (63.2%)
# Gaps:         115/587 (19.6%)
# Score: 1326.5
#=======================================

Reference_M1_      1 MAKNNTNRHYSLRKLKTGTASVAVALTVLGAGFANQTEVKANGDGNPREV     50
                     ||||.|||||||||||||||||||||||:|||......|:|:.
M12_sequence       1 MAKNTTNRHYSLRKLKTGTASVAVALTVVGAGLVAGQTVRADH-------     43

Reference_M1_     51 IEDLAANNPAIQNI-----RLR----------YENKD-------------     72
                      .||.|....::::     ||:          |:||.
M12_sequence      44 -SDLVAEKQRLEDLGQKFERLKQRSELYLQQYYDNKSNGYKGDWYVQQLK     92

Reference_M1_     73 -LKARLENAM-EVAGRDFKRA---------------EELEKAKQALEDQR    105
                      |...||.|. |::|...|.|               .||||:.:...|..
M12_sequence      93 MLNRDLEQAYNELSGEAHKDALGKLGIDNADLKAKITELEKSVEEKNDVL    142

Reference_M1_    106 KDLETKLKELQQDYDLAKESTSWDRQRLEKELEEKK----------EALE    145
                     ..::.:|:|.::|....:|..:.|..|.::|:.||:          :.|:
M12_sequence     143 SQIKKELEEAEKDIQFGREVHAADLLRHKQEIAEKENVISKLNGELQPLK    192

Reference_M1_    146 LAIDQASR----DYHRATALEKEL----EEKKKALELAI---DQASQDYN    184
                     ..:|:..|    :..:..:||::|    |..||..|||.   ..|.::||
M12_sequence     193 QKVDETDRNLQQEKQKVLSLEQQLAVTKENAKKDFELAALGHQLADKEYN    242

Reference_M1_    185 RANVLEKELETITREQEINRNLLG--NAKLELDQLSSEKEQLTIEKAKLE    232
                      |.:.|.|.:....::......||  :|..|.....:||:.   :..:||
M12_sequence     243 -AKIAELESKLADAKKDFELAALGHQHAHNEYQAKLAEKDG---QIKQLE    288

Reference_M1_    233 EEKQISDASRQSLRRDLDASREAKKQVEKDLANLTAELDKVKEDKQ----    278
                     |:|||.||||:...|||:|.|:|||..|.:|.||.|||.||.|.||
M12_sequence     289 EQKQILDASRKGTARDLEAVRQAKKATEAELNNLKAELAKVTEQKQILDA    338

Reference_M1_    279 -------------------------------ISDASRQGLRRDLDASREA    297
                                                    ||:|||:||||||||||||
M12_sequence     339 SRKGTARDLEAVRKAKAQVEAALKQLEEQNKISEASRKGLRRDLDASREA    388

Reference_M1_    298 KKQVEKDLANLTAELDKVKEEKQISDASRQGLRRDLDASREAKKQVEKAL    347
                     ||||||||||||||||||||||||||||||||||||||||||||||||||
M12_sequence     389 KKQVEKDLANLTAELDKVKEEKQISDASRQGLRRDLDASREAKKQVEKAL    438

Reference_M1_    348 EEANSKLAALEKLNKELEESKKLTEKEKAELQAKLEAEAKALKEQLAKQA    397
                     ||||||||||||||||||||||||||||||||||||||||||||||||||
M12_sequence     439 EEANSKLAALEKLNKELEESKKLTEKEKAELQAKLEAEAKALKEQLAKQA    488

Reference_M1_    398 EELAKLRAGKASDSQTPDTKPGNKAVPGKGQAPQAGTKPNQNKAPMKETK    447
                     ||||||||||||||||||.|||||||||||||||||||||||||||||||
M12_sequence     489 EELAKLRAGKASDSQTPDAKPGNKAVPGKGQAPQAGTKPNQNKAPMKETK    538

Reference_M1_    448 RQLPSTGETANPFFTAAALTVMATAGVAAVVKRKEEN    484
                     ||||||||.||||||||||||||||||||||||||||
M12_sequence     539 RQLPSTGEAANPFFTAAALTVMATAGVAAVVKRKEEN    575
#---------------------------------------
#---------------------------------------
```

```
########################################
# Program: needle
# Commandline: needle
#    -auto
#    -stdout
#    -asequence emboss_needle-I20210916-190109-0733-82553029-p2m.asequence
#    -bsequence emboss_needle-I20210916-190109-0733-82553029-p2m.bsequence
#    -datafile EBLOSUM62
#    -gapopen 10.0
#    -gapextend 0.5
#    -endopen 10.0
#    -endextend 0.5
#    -aformat3 pair
#    -sprotein1
#    -sprotein2
# Align_format: pair
# Report_file: stdout
########################################

#=======================================
#
# Aligned_sequences: 2
# 1: Reference_M1_sequence
# 2: M28_sequence
# Matrix: EBLOSUM62
# Gap_penalty: 10.0
# Extend_penalty: 0.5
#
# Length: 489
# Identity:     217/489 (44.4%)
# Similarity:   275/489 (56.2%)
# Gaps:         136/489 (27.8%)
# Score: 820.5
#
#
#=======================================

Reference_M1_      1 MAKNNTNRHYSLRKLKTGTASVAVALTVLGAGFANQTEVKANGDGNPREV     50
                     ||:.:||:.||||||||||||||||:.|||||||||||||
M28_sequence       1 MARKDTNKQYSLRKLKTGTASVAVAVAVLGAGFANQTEVK----------     40

Reference_M1_     51 IEDLAANNPAIQNIRLRYENKDLKARLENAMEVAGRDFKRAEELEKAKQA    100
                         ||.:|                             |..|........
M28_sequence      41 ----AAESP-----------------------------KSTETSANGADK     57

Reference_M1_    101 LEDQRKDLETKLKELQQDY----DLAKESTSWDRQRLE-KELEEKKEALE    145
                     |.|....|.|:.::|:.:|    |..:|...:...|.| ::|.||:...:
M28_sequence      58 LADAYNTLLTEHEKLRDEYYTLIDAKEEEPRYKALRGENQDLREKEGKYQ    107

Reference_M1_    146 LAIDQASRDYHRATALEKELEEKKKALELAIDQASQDYNRANVLEKELET    195
                     ..|              |:||||:|.||    :.|:|..| :.|:|
M28_sequence     108 DKI--------------KKLEEKEKNLE----KKSEDVER-HYLKK----    134

Reference_M1_    196 ITREQEINRNLLGNAKLELDQLSSEKEQLTIEKAKLEEEKQISDASRQSL    245
                                       |||...|:|         |.:|.:.:..|||
M28_sequence     135 ------------------LDQEHKEQE---------ERQKNLEELERQS-    156

Reference_M1_    246 RRDLDASREAKKQVEKDLANLTAELDKVKEDKQISDASRQGLRRDLDASR    295
                     :|::|  :..::|::|.        .:::.:||||:|||:.|.|||:|||
M28_sequence     157 QREID--KRYQEQLQKQ--------QQLETEKQISEASRKSLSRDLEASR    196

Reference_M1_    296 EAKKQVEKDLANLTAELDKVKEEKQISDASRQGLRRDLDASREAKKQVEK    345
                     .|||.:|       ||..|:||||||||||||||.|||:|||||||:||.
M28_sequence     197 AAKKDLE-------AEHQKLKEEKQISDASRQGLSRDLEASREAKKKVEA    239

Reference_M1_    346 ALEEANSKLAALEKLNKELEESKKLTEKEKAELQAKLEAEAKALKEQLAK    395
                     .|.||||||.|||||||||||.|||:|||||||||:||||||||||||||
M28_sequence     240 DLAEANSKLQALEKLNKELEEGKKLSEKEKAELQARLEAEAKALKEQLAK    289

Reference_M1_    396 QAEELAKLRAGKASDSQTPDTKPGNKAVPGKGQAPQAGTKPNQNKAPMKE    445
                     ||||||||:.     :|||:.|          .||||    |::::.|.:
M28_sequence     290 QAEELAKLKG-----NQTPNAK----------VAPQA----NRSRSAMTQ    320

Reference_M1_    446 TKRQLPSTGETANPFFTAAALTVMATAGVAAVVKRKEEN    484
                     .||.||||||.|||||||||.|||.:||:.| :||||||
M28_sequence     321 QKRTLPSTGEAANPFFTAAAATVMVSAGMLA-LKRKEEN    358


#---------------------------------------
#---------------------------------------
```

```
########################################
# Program: needle
# Commandline: needle
#    -auto
#    -stdout
#    -asequence emboss_needle-I20210916-190152-0642-38607375-p2m.asequence
#    -bsequence emboss_needle-I20210916-190152-0642-38607375-p2m.bsequence
#    -datafile EBLOSUM62
#    -gapopen 10.0
#    -gapextend 0.5
#    -endopen 10.0
#    -endextend 0.5
#    -aformat3 pair
#    -sprotein1
#    -sprotein2
# Align_format: pair
# Report_file: stdout
########################################

#=======================================
# Aligned_sequences: 2
# 1: Reference_M1_sequence
# 2: M75_sequence
# Matrix: EBLOSUM62
# Gap_penalty: 10.0
# Extend_penalty: 0.5
#
# Length: 485
# Identity:     245/485 (50.5%)
# Similarity:   306/485 (63.1%)
# Gaps:          99/485 (20.4%)
# Score: 1021.0
#=======================================

Reference_M1_      1 MAKNNTNRHYSLRKLKTGTASVAVALTVLGAGFANQTEVKANGDGNPREV     50
                     ||:.:||:.||||||||||||||||:.||||||||||||||..:....|
M75_sequence       1 MARKDTNKQYSLRKLKTGTASVAVAVAVLGAGFANQTEVKAEEERTFTE-     49

Reference_M1_     51 IEDLAANNPAIQNIRLRYENKDLKARLENAMEVAGRDFKRAEELEKAKQA    100
                                    |.||     ||.:
M75_sequence      50 ---------------LPYE-----ARYK----------------------     57

Reference_M1_    101 LEDQRKDLETKLKELQQDYDLAKESTSWDRQRLEKELEEKKEALELAIDQ    150
                                               :|     :.|.:|.:|.....:|:
M75_sequence      58 --------------------------AW-----KSENDELRENYRRTLDK     76

Reference_M1_    151 ASRDYHRATALEKELEEKKKALELA-IDQASQDYNRANVLEKELETITRE    199
                     .:.:..:.|.||:  :.||...||| :.:.......|:  :|:::.:|..
M75_sequence      77 FNTEQGKTTRLEE--QNKKLHSELASVTETLTSVTEAD--DKKIKDLTDR    122

Reference_M1_    200 QEINRNLLGNAKLELDQLSSEKEQLTIEKAKLEEEKQISDASRQSLRRDL    249
                     .:|:.|||||||.::::|::||:.|..:..||||:||||||||:||.|||
M75_sequence     123 DKISSNLLGNAKDQINKLTTEKDTLAEKAKKLEEDKQISDASRKSLSRDL    172

Reference_M1_    250 DASREAKKQVEKDLANLTAELDKVKEDKQISDASRQGLRRDLDASREAKK    299
                     :|||.|||::|.:...|..|..|:||:|||||||||||.|||:|||||||
M75_sequence     173 EASRAAKKELEANHQKLETEHKKLKEEKQISDASRQGLSRDLEASREAKK    222

Reference_M1_    300 QVEKDLANLTAELDKVKEEKQISDASRQGLRRDLDASREAKKQVEKALEE    349
                     :||.|||.||||..|:||||||||||||||.|||:|||||||:||..|.|
M75_sequence     223 KVEADLAALTAEHQKLKEEKQISDASRQGLSRDLEASREAKKKVEADLAE    272

Reference_M1_    350 ANSKLAALEKLNKELEESKKLTEKEKAELQAKLEAEAKALKEQLAKQAEE    399
                     |||||.|||||||||||.|||:|||||||||:||||||||||||||||||
M75_sequence     273 ANSKLQALEKLNKELEEGKKLSEKEKAELQARLEAEAKALKEQLAKQAEE    322

Reference_M1_    400 LAKLRAGKASDSQTPDTKPGNKAVPGKGQAPQAGTKPNQNKAPMKETKRQ    449
                     ||||:.     :|||:.|          .||||    |::::.|.:.||.
M75_sequence     323 LAKLKG-----NQTPNAK----------VAPQA----NRSRSAMTQQKRT    353

Reference_M1_    450 LPSTGETANPFFTAAALTVMATAGVAAVVKRKEEN    484
                     ||||||.|||||||||.|||.:||:.| :||||||
M75_sequence     354 LPSTGEAANPFFTAAAATVMVSAGMLA-LKRKEEN    387


#---------------------------------------
#---------------------------------------
```

```
########################################
# Program: needle
# Commandline: needle
#    -auto
#    -stdout
#    -asequence emboss_needle-I20210916-190239-0819-2068156-p2m.asequence
#    -bsequence emboss_needle-I20210916-190239-0819-2068156-p2m.bsequence
#    -datafile EBLOSUM62
#    -gapopen 10.0
#    -gapextend 0.5
#    -endopen 10.0
#    -endextend 0.5
#    -aformat3 pair
#    -sprotein1
#    -sprotein2
# Align_format: pair
# Report_file: stdout
########################################

#=======================================
# Aligned_sequences: 2
# 1: Reference_M1_sequence
# 2: M79_sequence
# Matrix: EBLOSUM62
# Gap_penalty: 10.0
# Extend_penalty: 0.5
#
# Length: 486
# Identity:     125/486 (25.7%)
# Similarity:   157/486 (32.3%)
# Gaps:         266/486 (54.7%)
# Score: 461.0
#=======================================

Reference_M1_      1 MAKNNTNRHYSLRKLKTGTASVAVALTVLGAGFANQTEVKANGDGNPREV     50
                     ||:.:||:.||||||||||||||||:.||||||||||||||.....|:
M79_sequence       1 MARKDTNKQYSLRKLKTGTASVAVAVAVLGAGFANQTEVKAAEIKKPQ--     48

Reference_M1_     51 IEDLAANNPAIQNIRLRYENKDLKARLENAMEVAGRDFKRAEELEKAKQ-     99
                      .|.|.|.|...|..|: ||::||...|..:..       |::.||..|
M79_sequence      49 -ADSAWNWPKEYNALLK-ENEELKVEREKYLSY-------ADDKEKDPQY     89

Reference_M1_    100 -ALEDQRKDLETKLKELQQDYDLAKESTSWDRQRLEKELEEKKEALELAI    148
                      ||..:.:||..:..:.|...           :.||||.:||:|..|
M79_sequence      90 RALMGENQDLRKREGQYQDKI-----------EELEKERKEKQERQE---    125

Reference_M1_    149 DQASRDYHRATALEKELEEKKKALELAIDQASQDYNRANVLEKELETITR    198
                      |..|.|.  ...:|..:|::|                           :
M79_sequence     126 -QLERQYQ--IEADKHYQEQQK---------------------------K    145

Reference_M1_    199 EQEINRNLLGNAKLELDQLSSEKEQLTIEKAKLEEEKQISDASRQSLRRD    248
                     .|:                  |::||..||.||.::|||||||||.|.||
M79_sequence     146 HQQ------------------EQQQLEAEKQKLAKDKQISDASRQGLSRD    177

Reference_M1_    249 LDASREAKKQVEKDLANLTAELDKVKEDKQISDASRQGLRRDLDASREAK    298
                     |:|||||||:||.|||.||||..|:||||||||||||||.|||:|
M79_sequence     178 LEASREAKKKVEADLAALTAEHQKLKEDKQISDASRQGLSRDLEA-----    222

Reference_M1_    299 KQVEKDLANLTAELDKVKEEKQISDASRQGLRRDLDASREAKKQVEKALE    348

M79_sequence     223 --------------------------------------------------    222

Reference_M1_    349 EANSKLAALEKLNKELEESKKLTEKEKAELQAKLEAEAKALKEQLAKQAE    398

M79_sequence     223 --------------------------------------------------    222

Reference_M1_    399 ELAKLRAGKASDSQTPDTKPGNKAVPGKGQAPQAGTKPNQNKAPMKETKR    448

M79_sequence     223 --------------------------------------------------    222

Reference_M1_    449 QLPSTGETANPFFTAAALTVMATAGVAAVVKRKEEN    484

M79_sequence     223 ------------------------------------    222


#---------------------------------------
#---------------------------------------
```

```
########################################
# Program: needle
# Commandline: needle
#    -auto
#    -stdout
#    -asequence emboss_needle-I20210916-190331-0721-53653255-p2m.asequence
#    -bsequence emboss_needle-I20210916-190331-0721-53653255-p2m.bsequence
#    -datafile EBLOSUM62
#    -gapopen 10.0
#    -gapextend 0.5
#    -endopen 10.0
#    -endextend 0.5
#    -aformat3 pair
#    -sprotein1
#    -sprotein2
# Align_format: pair
# Report_file: stdout
########################################

#=======================================
#
# Aligned_sequences: 2
# 1: Reference_M1_sequence
# 2: M81_sequence
# Matrix: EBLOSUM62
# Gap_penalty: 10.0
# Extend_penalty: 0.5
#
# Length: 484
# Identity:     278/484 (57.4%)
# Similarity:   316/484 (65.3%)
# Gaps:          88/484 (18.2%)
# Score: 1204.5
#=======================================

Reference_M1_      1 MAKNNTNRHYSLRKLKTGTASVAVALTVLGAGFANQTEVKANGDGNPREV     50
                     |.:.:||:.||||||||||||||||:.||||||||||||||.|.
M81_sequence       1 MVRKDTNKQYSLRKLKTGTASVAVAVAVLGAGFANQTEVKAAGS------     44

Reference_M1_     51 IEDLAANNPAIQNIRLRYENKDLKARLENAMEVAGRDFKRAEELEKAKQA    100
                         ..|.|..|...|..||:||:          ||:.|...:|||
M81_sequence      45 ----EENVPKQQYNALWEENEDLR----------GRERKYIAKLEK----     76

Reference_M1_    101 LEDQRKDLETKLKELQQDYDLAKESTSWDRQRLEKELEEKKEALELAIDQ    150
                                 :|:|..
M81_sequence      77 ------------EEIQNG--------------------------------     82

Reference_M1_    151 ASRDYHRATALEKELEEKKKALELAIDQASQDYNRANVLEKELETITREQ    200
                                  ||.||.:.||..|... ||     |:|...:.|.|:.
M81_sequence      83 -------------ELNEKNRKLEADIADL-QD-----VIEDNDQEIKRKD    113

Reference_M1_    201 EINRNLLGNAKLELDQLSSEKEQLTIEKAKLEEEKQISDASRQSLRRDLD    250
                     .:....|..:|.::..|::||:.|..:..||||:||||||||:||.|||:
M81_sequence     114 RMYEAFLKQSKDQVKDLTAEKDTLAEKAKKLEEDKQISDASRKSLSRDLE    163

Reference_M1_    251 ASREAKKQVEKDLANLTAELDKVKEDKQISDASRQGLRRDLDASREAKKQ    300
                     .||.|||::|.....|..|..|:||||||||||||||.|||:|||||||:
M81_sequence     164 GSRAAKKELEAKHQKLETEHQKLKEDKQISDASRQGLSRDLEASREAKKK    213

Reference_M1_    301 VEKDLANLTAELDKVKEEKQISDASRQGLRRDLDASREAKKQVEKALEEA    350
                     ||.|||.||||..|:||||||||||||||.|||:|||||||:||..|.||
M81_sequence     214 VEADLAALTAEHQKLKEEKQISDASRQGLSRDLEASREAKKKVEADLAEA    263

Reference_M1_    351 NSKLAALEKLNKELEESKKLTEKEKAELQAKLEAEAKALKEQLAKQAEEL    400
                     ||||.|||||||||||.|||:|||||||||:|||||||||||||||||||
M81_sequence     264 NSKLQALEKLNKELEEGKKLSEKEKAELQARLEAEAKALKEQLAKQAEEL    313

Reference_M1_    401 AKLRAGKASDSQTPDTKPGNKAVPGKGQAPQAGTKPNQNKAPMKETKRQL    450
                     |||||||||||||||.|||||.||||||||||||||||||||||||||||
M81_sequence     314 AKLRAGKASDSQTPDAKPGNKVVPGKGQAPQAGTKPNQNKAPMKETKRQL    363

Reference_M1_    451 PSTGETANPFFTAAALTVMATAGVAAVVKRKEEN    484
                     |||||.|||||||||.|||.:||:.| :||||||
M81_sequence     364 PSTGEAANPFFTAAAATVMVSAGMLA-LKRKEEN    396


#---------------------------------------
#---------------------------------------
```

```
########################################
# Program: needle
# Commandline: needle
#    -auto
#    -stdout
#    -asequence emboss_needle-I20210916-190422-0059-21208464-p2m.asequence
#    -bsequence emboss_needle-I20210916-190422-0059-21208464-p2m.bsequence
#    -datafile EBLOSUM62
#    -gapopen 10.0
#    -gapextend 0.5
#    -endopen 10.0
#    -endextend 0.5
#    -aformat3 pair
#    -sprotein1
#    -sprotein2
# Align_format: pair
# Report_file: stdout
########################################

#=======================================
# Aligned_sequences: 2
# 1: Reference_M1_sequence
# 2: M87_sequence
# Matrix: EBLOSUM62
# Gap_penalty: 10.0
# Extend_penalty: 0.5
#
# Length: 487
# Identity:     131/487 (26.9%)
# Similarity:   175/487 (35.9%)
# Gaps:         252/487 (51.7%)
# Score: 481.0
#=======================================

Reference_M1_      1 MAKNNTNRHYSLRKLKTGTASVAVALTVLGAGFANQTEVKANGDGNPREV     50
                     ||:.:||:.||||||||||||||||:.||||||||||||||.   :||||
M87_sequence       1 MARKDTNKQYSLRKLKTGTASVAVAVAVLGAGFANQTEVKAE---SPREV     47

Reference_M1_     51 IEDLAANNPAIQNIRLRYENKDLKARLENAMEVAGRDFKRAEELEKAKQA    100
                     ..:|||:              ..|.::|.|.|       :|.:|||   .
M87_sequence      48 TNELAAS--------------VWKKKVEEAKE-------KASKLEK---Q     73

Reference_M1_    101 LEDQRKD---LETKLKELQQDYDLAKESTSWDRQRLEKELEEKKEALELA    147
                     ||:.:||   :|.||::...|||           :|||  |.|:.|.:|.
M87_sequence      74 LEEAQKDYSEIEGKLEQFWHDYD-----------KLEK--ENKEYASQLG    110

Reference_M1_    148 IDQASRDYHRATALEKELEEKKKALELAIDQASQDYNRANVLEKELETIT    197
                     .:|..|:......|.|..||.|:..:...:|..:        :|.||.:.
M87_sequence     111 KNQEEREKLELEYLRKSDEEYKEHQQYRQEQEER--------QKNLEELE    152

Reference_M1_    198 REQEINRNLLGNAKLELDQLSSEKEQLTIEKAKLEEEKQISDASRQSLRR    247
                     |:.          |.|:|:...|:.|   ::.:||.|||||:|||:||.|
M87_sequence     153 RQN----------KREIDKRYQEQLQ---KQQQLETEKQISEASRKSLSR    189

Reference_M1_    248 DLDASREAKKQVEKDLANLTAELDKVKEDKQISDASRQGLRRDLDASREA    297
                     ||:|||.|||::|       ||..|:||:||||||||:.|.|||:|||||
M87_sequence     190 DLEASRAAKKELE-------AEHQKLKEEKQISDASRKSLSRDLEASREA    232

Reference_M1_    298 KKQVEKDLANLTAELDKVKEEKQISDASRQGLRRDLDASREAKKQVEKAL    347
                     ||:||.
M87_sequence     233 KKKVEA--------------------------------------------    238

Reference_M1_    348 EEANSKLAALEKLNKELEESKKLTEKEKAELQAKLEAEAKALKEQLAKQA    397

M87_sequence     239 --------------------------------------------------    238

Reference_M1_    398 EELAKLRAGKASDSQTPDTKPGNKAVPGKGQAPQAGTKPNQNKAPMKETK    447

M87_sequence     239 --------------------------------------------------    238

Reference_M1_    448 RQLPSTGETANPFFTAAALTVMATAGVAAVVKRKEEN    484

M87_sequence     239 -------------------------------------    238


#---------------------------------------
#---------------------------------------
```

```
########################################
# Program: needle
# Commandline: needle
#    -auto
#    -stdout
#    -asequence emboss_needle-I20210916-190502-0908-95709551-p1m.asequence
#    -bsequence emboss_needle-I20210916-190502-0908-95709551-p1m.bsequence
#    -datafile EBLOSUM62
#    -gapopen 10.0
#    -gapextend 0.5
#    -endopen 10.0
#    -endextend 0.5
#    -aformat3 pair
#    -sprotein1
#    -sprotein2
# Align_format: pair
# Report_file: stdout
########################################

#=======================================
#
# Aligned_sequences: 2
# 1: Reference_M1_sequence
# 2: M89_sequence
# Matrix: EBLOSUM62
# Gap_penalty: 10.0
# Extend_penalty: 0.5
#
# Length: 487
# Identity:     148/487 (30.4%)
# Similarity:   191/487 (39.2%)
# Gaps:         246/487 (50.5%)
# Score: 574.0
#
#=======================================

Reference_M1_      1 MAKNNTNRHYSLRKLKTGTASVAVALTVLGAGFANQTEVKANGDGNPREV     50
                     ||:.:||:.||||||||||||||||:.||||||||||.|||:.|...|.|
M89_sequence       1 MARKDTNKQYSLRKLKTGTASVAVAVAVLGAGFANQTTVKADSDNINRSV     50

Reference_M1_     51 IEDLAANNPAIQNIRLRYENKDLKARLENAMEVAGRDFKRAEELEKAKQA    100
                                                                      :
M89_sequence      51 -------------------------------------------------S     51

Reference_M1_    101 LEDQRKDLETKLKELQQDYDLAKESTSWDRQRLEK--ELEEKKEALELAI    148
                     ::|..|:|..|:.:|:::..          :.|:|  ||:|:.:|.|.:.
M89_sequence      52 VKDNEKELHNKIADLEEERG----------EHLDKIDELKEELKAKEKSS     91

Reference_M1_    149 DQASRDYHRATALE-KELEEKKKALELAIDQASQDYNRANVLEKELETIT    197
                     :...|.|.|....| ||.:|::|.|| .:::.||         :|:|...
M89_sequence      92 ENVERHYLRKLDQEYKEQQERQKNLE-ELERQSQ---------REVEKRY    131

Reference_M1_    198 REQEINRNLLGNAKLELDQLSSEKEQLTIEKAKLEEEKQISDASRQSLRR    247
                     :||                  .:|:|      :||.|||||:|||:||.|
M89_sequence     132 QEQ------------------LQKQQ------QLETEKQISEASRKSLSR    157

Reference_M1_    248 DLDASREAKKQVEKDLANLTAELDKVKEDKQISDASRQGLRRDLDASREA    297
                     ||:|||.|||.:|       ||..|:||:|||||||||||.|||:|||||
M89_sequence     158 DLEASRAAKKDLE-------AEHQKLKEEKQISDASRQGLSRDLEASREA    200

Reference_M1_    298 KKQVEKDLANLTAELDKVKEEKQISDASRQGLRRDLDASREAKKQVEKAL    347
                     ||:||.|||.||||..|:||||||||||||||.|||:|||||||
M89_sequence     201 KKKVEADLAALTAEHQKLKEEKQISDASRQGLSRDLEASREAKK------    244

Reference_M1_    348 EEANSKLAALEKLNKELEESKKLTEKEKAELQAKLEAEAKALKEQLAKQA    397

M89_sequence     245 --------------------------------------------------    244

Reference_M1_    398 EELAKLRAGKASDSQTPDTKPGNKAVPGKGQAPQAGTKPNQNKAPMKETK    447

M89_sequence     245 --------------------------------------------------    244

Reference_M1_    448 RQLPSTGETANPFFTAAALTVMATAGVAAVVKRKEEN    484

M89_sequence     245 -------------------------------------    244


#---------------------------------------
#---------------------------------------
```

```
########################################
# Program: needle
# Commandline: needle
#    -auto
#    -stdout
#    -asequence emboss_needle-I20210916-190546-0891-61986443-p2m.asequence
#    -bsequence emboss_needle-I20210916-190546-0891-61986443-p2m.bsequence
#    -datafile EBLOSUM62
#    -gapopen 10.0
#    -gapextend 0.5
#    -endopen 10.0
#    -endextend 0.5
#    -aformat3 pair
#    -sprotein1
#    -sprotein2
# Align_format: pair
# Report_file: stdout
########################################

#=======================================
# Aligned_sequences: 2
# 1: Reference_M1_sequence
# 2: M179_sequence
# Matrix: EBLOSUM62
# Gap_penalty: 10.0
# Extend_penalty: 0.5
#
# Length: 498
# Identity:     294/498 (59.0%)
# Similarity:   332/498 (66.7%)
# Gaps:          79/498 (15.9%)
# Score: 1218.5
#=======================================

Reference_M1_      1 MAKNNTNRHYSLRKLKTGTASVAVALTVLGAGFA-NQTEVKANGDGNPRE     49
                     |.:.:|||||||||||||||||||||:|||||.| |||||.|.|.  ...|.
M179_sequence      1 MVRKDTNRHYSLRKLKTGTASVAVALSVLGAGLAVNQTEVSAK--SVTRS     48

Reference_M1_     50 VIEDLAANNPAIQNIRLRYENKDLKARLE-NAMEVAGRDFKRAEELEKAK     98
                     ..:|...:..||    ..||.::.|...| ||             |....
M179_sequence     49 TAQDPDKSRQAI----TEYEVENHKLTQEKNA-------------LTNRN     81

Reference_M1_     99 QALEDQRKDLETKLKELQQDYDLAKESTSWDRQRLEKELEEKK-------    141
                     |.|.|:..:|:|..:.|:|..|....    .|.:|||:::||:
M179_sequence     82 QELTDENGELKTANEALRQRGDTLFN----QRVKLEKQVQEKEHNNKTLK    127

Reference_M1_    142 -EALELAIDQASRDYHRATALEKELEEKKALELAIDQASQDYNRA---N    187
                      |..||..:...        |.|:|:|.::  |||..|.....|..   .
M179_sequence    128 IENGELKTENGD--------LTKKLDETRQ--ELANKQQESKENEKTLNE    167

Reference_M1_    188 VLEKEL-ETITREQEINRNLLGNAKLELDQLSSEKEQLTIEKAKLEEEKQ    236
                     :|||.: :.|.:||| |:..:|..|..||:...:|      .||.::.||
M179_sequence    168 LLEKTVKDKIAKEQE-NKETIGTLKKLLDETVKDK------IAKEQKSKQ    210

Reference_M1_    237 ISDASRQSLRRDLDASREAKKQVEKDLANLTAELDKVKEDKQISDASRQG    286
                                              |...|..||.|.:|..:||||||||
M179_sequence    211 -------------------------DFGALKQELAKKEEQNKISDASRQG    235

Reference_M1_    287 LRRDLDASREAKKQVEKDLANLTAELDKVKEEKQISDASRQGLRRDLDAS    336
                     |||||:||||||||||||||||||||||||||||:|||||||||||||||
M179_sequence    236 LRRDLNASREAKKQVEKDLANLTAELDKVKEEKQVSDASRQGLRRDLDAS    285

Reference_M1_    337 REAKKQVEKALEEANSKLAALEKLNKELEESKKLTEKEKAELQAKLEAEA    386
                     ||||||||||||||||||||||||||||||||||||||||||||||||||
M179_sequence    286 REAKKQVEKALEEANSKLAALEKLNKELEESKKLTEKEKAELQAKLEAEA    335

Reference_M1_    387 KALKEQLAKQAEELAKLRAGKASDSQTPDTKPGNKAVPGKGQAPQAGTKP    436
                     |||||:|||||||||||||||||||||||.||||||||||||||||||||
M179_sequence    336 KALKEKLAKQAEELAKLRAGKASDSQTPDAKPGNKAVPGKGQAPQAGTKP    385

Reference_M1_    437 NQNKAPMKETKRQLPSTGETANPFFTAAALTVMATAGVAAVVKRKEEN    484
                     |||||||||||||||||||.|||||||||||||||||||||||||:||
M179_sequence    386 NQNKAPMKETKRQLPSTGEAANPFFTAAALTVMATAGVAAVVKRKKEN    433


#---------------------------------------
#---------------------------------------
```

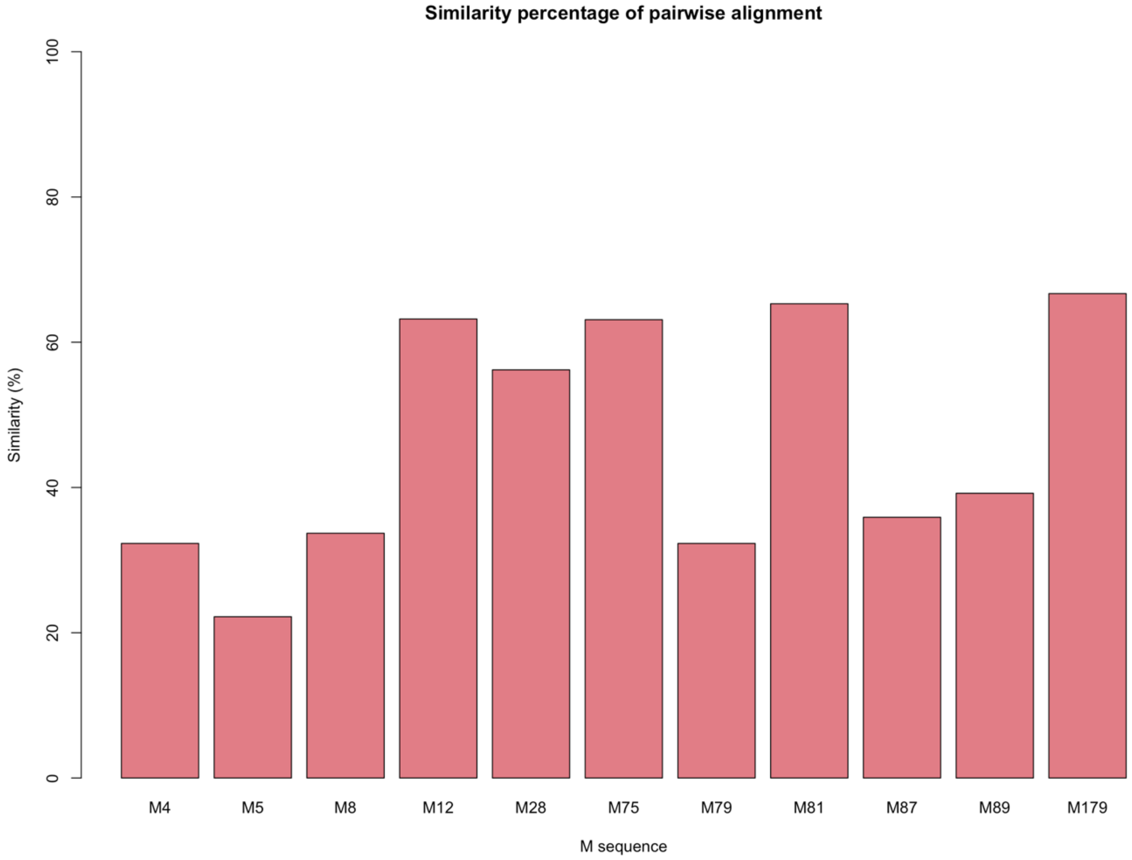


*Figure S1. Similarity percentage for each M sequence aligned to M1 reference sequence*

**Binding peptides similarity in each sequence**

Here, only the two peptides of M1 protein in the binding interface with ab25 *i.e.*, “KALELAIDQASQDYNRANVLEKE”, and “REAKKQVEKDLANLTAELDKVKE” were searched in each target M sequence to detect the similar patterns. Below, we report the result of the alignment and the similarity percentage with each of the detected peptides.

```
########################################
# Program: needle
# Commandline: needle
#    -auto
#    -stdout
#    -asequence emboss_needle-I20210917-134146-0008-34466034-p1m.asequence
#    -bsequence emboss_needle-I20210917-134146-0008-34466034-p1m.bsequence
#    -datafile EBLOSUM62
#    -gapopen 10.0
#    -gapextend 0.5
#    -endopen 10.0
#    -endextend 0.5
#    -aformat3 pair
#    -sprotein1
#    -sprotein2
# Align_format: pair
# Report_file: stdout
########################################

#=======================================
#
# Aligned_sequences: 2
# 1: Binding_pep1
# 2: M4_similar_p1
# Matrix: EBLOSUM62
# Gap_penalty: 10.0
# Extend_penalty: 0.5
#
# Length: 26
# Identity:      10/26 (38.5%)
# Similarity:    12/26 (46.2%)
# Gaps:           3/26 (11.5%)
# Score: 29.0
#
#
#=======================================

Binding_pep1       1 KALELAIDQASQDYN---RANVLEKE     23
                     ||.|:...||...:|   ..|.|.||
M4_similar_p1      1 KAAEIKKPQADSAWNWPKEYNALLKE     26

#---------------------------------------
#---------------------------------------
```

```
########################################
# Program: needle
# Commandline: needle
#    -auto
#    -stdout
#    -asequence emboss_needle-I20210917-134735-0837-62507407-p1m.asequence
#    -bsequence emboss_needle-I20210917-134735-0837-62507407-p1m.bsequence
#    -datafile EBLOSUM62
#    -gapopen 10.0
#    -gapextend 0.5
#    -endopen 10.0
#    -endextend 0.5
#    -aformat3 pair
#    -sprotein1
#    -sprotein2
# Align_format: pair
# Report_file: stdout
########################################

#=======================================
#
# Aligned_sequences: 2
# 1: Binding_pep2
# 2: M4_similar_p2
# Matrix: EBLOSUM62
# Gap_penalty: 10.0
# Extend_penalty: 0.5
#
# Length: 23
# Identity:      17/23 (73.9%)
# Similarity:    19/23 (82.6%)
# Gaps:           0/23 ( 0.0%)
# Score: 76.0
#
#
#=======================================

Binding_pep2       1 REAKKQVEKDLANLTAELDKVKE     23
                     |||||:||.|||.||||..|:||
M4_similar_p2      1 REAKKKVEADLAALTAEHQKLKE     23


#---------------------------------------
#---------------------------------------
```

```
########################################
# Program: needle
# Commandline: needle
#    -auto
#    -stdout
#    -asequence emboss_needle-I20210917-135430-0963-60062530-p2m.asequence
#    -bsequence emboss_needle-I20210917-135430-0963-60062530-p2m.bsequence
#    -datafile EBLOSUM62
#    -gapopen 10.0
#    -gapextend 0.5
#    -endopen 10.0
#    -endextend 0.5
#    -aformat3 pair
#    -sprotein1
#    -sprotein2
# Align_format: pair
# Report_file: stdout
########################################

#=======================================
#
# Aligned_sequences: 2
# 1: Binding_pep1
# 2: M5_similar_p1
# Matrix: EBLOSUM62
# Gap_penalty: 10.0
# Extend_penalty: 0.5
#
# Length: 23
# Identity:       6/23 (26.1%)
# Similarity:    10/23 (43.5%)
# Gaps:           0/23 ( 0.0%)
# Score: 17.0
#
#
#=======================================

Binding_pep1       1 KALELAIDQASQDYNRANVLEKE     23
                     |||...:::..:|........||
M5_similar_p1      1 KALNELLEKTVKDKIAKEQENKE     23


#---------------------------------------
#---------------------------------------
```

```
########################################
# Program: needle
# Commandline: needle
#    -auto
#    -stdout
#    -asequence emboss_needle-I20210917-135546-0415-63161967-p2m.asequence
#    -bsequence emboss_needle-I20210917-135546-0415-63161967-p2m.bsequence
#    -datafile EBLOSUM62
#    -gapopen 10.0
#    -gapextend 0.5
#    -endopen 10.0
#    -endextend 0.5
#    -aformat3 pair
#    -sprotein1
#    -sprotein2
# Align_format: pair
# Report_file: stdout
########################################

#=======================================
#
# Aligned_sequences: 2
# 1: Binding_pep2
# 2: M5_similar_p2
# Matrix: EBLOSUM62
# Gap_penalty: 10.0
# Extend_penalty: 0.5
#
# Length: 23
# Identity:       3/23 (13.0%)
# Similarity:     8/23 (34.8%)
# Gaps:          10/23 (43.5%)
# Score: 18.0
#
#
#=======================================

Binding_pep2       1 REAKKQVEKDLANLTAELDKVKE     23
                       ...|:.:|:::.|
M5_similar_p2      1 STQKETLEREVQN----------     13


#---------------------------------------
#---------------------------------------
```

```
########################################
# Program: needle
# Commandline: needle
#    -auto
#    -stdout
#    -asequence emboss_needle-I20210917-135644-0668-46832847-p2m.asequence
#    -bsequence emboss_needle-I20210917-135644-0668-46832847-p2m.bsequence
#    -datafile EBLOSUM62
#    -gapopen 10.0
#    -gapextend 0.5
#    -endopen 10.0
#    -endextend 0.5
#    -aformat3 pair
#    -sprotein1
#    -sprotein2
# Align_format: pair
# Report_file: stdout
########################################

#=======================================
#
# Aligned_sequences: 2
# 1: Binding_pep1
# 2: M8_similar_p1
# Matrix: EBLOSUM62
# Gap_penalty: 10.0
# Extend_penalty: 0.5
#
# Length: 25
# Identity:       6/25 (24.0%)
# Similarity:    13/25 (52.0%)
# Gaps:           2/25 ( 8.0%)
# Score: 17.5
#
#
#=======================================

Binding_pep1       1 KALELAIDQASQDYNR--ANVLEKE     23
                     :..|..:::.|:|..|  ...|::|
M8_similar_p1      1 EEKEKNLEKKSEDVERHYLKKLDQE     25


#---------------------------------------
#---------------------------------------
```

```
########################################
# Program: needle
# Commandline: needle
#    -auto
#    -stdout
#    -asequence emboss_needle-I20210917-135804-0655-27464049-p2m.asequence
#    -bsequence emboss_needle-I20210917-135804-0655-27464049-p2m.bsequence
#    -datafile EBLOSUM62
#    -gapopen 10.0
#    -gapextend 0.5
#    -endopen 10.0
#    -endextend 0.5
#    -aformat3 pair
#    -sprotein1
#    -sprotein2
# Align_format: pair
# Report_file: stdout
########################################

#=======================================
#
# Aligned_sequences: 2
# 1: Binding_pep2
# 2: M8_similar_p2
# Matrix: EBLOSUM62
# Gap_penalty: 10.0
# Extend_penalty: 0.5
#
# Length: 23
# Identity:      10/23 (43.5%)
# Similarity:    13/23 (56.5%)
# Gaps:           3/23 (13.0%)
# Score: 34.0
# 
#
#=======================================

Binding_pep2       1 REAKKQVEKDLANLTAELDKVKE     23
                     .||.:..:||   |.||..|:||
M8_similar_p2      1 LEASRAAKKD---LEAEHQKLKE     20


#---------------------------------------
#---------------------------------------
```

```
########################################
# Program: needle
# Commandline: needle
#    -auto
#    -stdout
#    -asequence emboss_needle-I20210917-135919-0762-92494436-p2m.asequence
#    -bsequence emboss_needle-I20210917-135919-0762-92494436-p2m.bsequence
#    -datafile EBLOSUM62
#    -gapopen 10.0
#    -gapextend 0.5
#    -endopen 10.0
#    -endextend 0.5
#    -aformat3 pair
#    -sprotein1
#    -sprotein2
# Align_format: pair
# Report_file: stdout
########################################

#=======================================
#
# Aligned_sequences: 2
# 1: Binding_pep1
# 2: M12_similar_p1
# Matrix: EBLOSUM62
# Gap_penalty: 10.0
# Extend_penalty: 0.5
#
# Length: 23
# Identity:       7/23 (30.4%)
# Similarity:    12/23 (52.2%)
# Gaps:           0/23 ( 0.0%)
# Score: 29.0
#
#
#=======================================

Binding_pep1       1 KALELAIDQASQDYNRANVLEKE     23
                     |.:|.|:::|:........|.||
M12_similar_p      1 KQVEKALEEANSKLAALEKLNKE     23


#---------------------------------------
#---------------------------------------
```

```
########################################
# Program: needle
# Commandline: needle
#    -auto
#    -stdout
#    -asequence emboss_needle-I20210917-140032-0915-10411681-p2m.asequence
#    -bsequence emboss_needle-I20210917-140032-0915-10411681-p2m.bsequence
#    -datafile EBLOSUM62
#    -gapopen 10.0
#    -gapextend 0.5
#    -endopen 10.0
#    -endextend 0.5
#    -aformat3 pair
#    -sprotein1
#    -sprotein2
# Align_format: pair
# Report_file: stdout
########################################

#=======================================
#
# Aligned_sequences: 2
# 1: Binding_pep2
# 2: M12_similar_p2
# Matrix: EBLOSUM62
# Gap_penalty: 10.0
# Extend_penalty: 0.5
#
# Length: 23
# Identity:      23/23 (100.0%)
# Similarity:    23/23 (100.0%)
# Gaps:           0/23 ( 0.0%)
# Score: 110.0
#
#
#=======================================

Binding_pep2       1 REAKKQVEKDLANLTAELDKVKE     23
                     |||||||||||||||||||||||
M12_similar_p      1 REAKKQVEKDLANLTAELDKVKE     23


#---------------------------------------
#---------------------------------------
```

```
########################################
# Program: needle
# Commandline: needle
#    -auto
#    -stdout
#    -asequence emboss_needle-I20210917-140131-0771-94163540-p2m.asequence
#    -bsequence emboss_needle-I20210917-140131-0771-94163540-p2m.bsequence
#    -datafile EBLOSUM62
#    -gapopen 10.0
#    -gapextend 0.5
#    -endopen 10.0
#    -endextend 0.5
#    -aformat3 pair
#    -sprotein1
#    -sprotein2
# Align_format: pair
# Report_file: stdout
########################################

#=======================================
#
# Aligned_sequences: 2
# 1: Binding_pep1
# 2: M28_similar_p1
# Matrix: EBLOSUM62
# Gap_penalty: 10.0
# Extend_penalty: 0.5
#
# Length: 23
# Identity:       6/23 (26.1%)
# Similarity:    10/23 (43.5%)
# Gaps:           0/23 ( 0.0%)
# Score: 22.0
#
#
#=======================================

Binding_pep1       1 KALELAIDQASQDYNRANVLEKE     23
                       |.:|..:.:|:........|.||
M28_similar_p      1 KKVEADLAEANSKLQALEKLNKE     23


#---------------------------------------
#---------------------------------------
```

```
########################################
# Program: needle
# Commandline: needle
#    -auto
#    -stdout
#    -asequence emboss_needle-I20210917-140229-0375-95050436-p2m.asequence
#    -bsequence emboss_needle-I20210917-140229-0375-95050436-p2m.bsequence
#    -datafile EBLOSUM62
#    -gapopen 10.0
#    -gapextend 0.5
#    -endopen 10.0
#    -endextend 0.5
#    -aformat3 pair
#    -sprotein1
#    -sprotein2
# Align_format: pair
# Report_file: stdout
########################################

#=======================================
#
# Aligned_sequences: 2
# 1: Binding_pep2
# 2: M28_similar_p2
# Matrix: EBLOSUM62
# Gap_penalty: 10.0
# Extend_penalty: 0.5
#
# Length: 23
# Identity:      11/23 (47.8%)
# Similarity:    17/23 (73.9%)
# Gaps:           0/23 ( 0.0%)
# Score: 54.0
#
#
#=======================================

Binding_pep2       1 REAKKQVEKDLANLTAELDKVKE     23
                     |||||:||.|||...::|..:::
M28_similar_p      1 REAKKKVEADLAEANSKLQALEK     23


#---------------------------------------
#---------------------------------------
```

```
########################################
# Program: needle
# Commandline: needle
#    -auto
#    -stdout
#    -asequence emboss_needle-I20210917-140324-0121-43219167-p2m.asequence
#    -bsequence emboss_needle-I20210917-140324-0121-43219167-p2m.bsequence
#    -datafile EBLOSUM62
#    -gapopen 10.0
#    -gapextend 0.5
#    -endopen 10.0
#    -endextend 0.5
#    -aformat3 pair
#    -sprotein1
#    -sprotein2
# Align_format: pair
# Report_file: stdout
########################################

#=======================================
#
# Aligned_sequences: 2
# 1: Binding_pep1
# 2: M75_similar_p1
# Matrix: EBLOSUM62
# Gap_penalty: 10.0
# Extend_penalty: 0.5
#
# Length: 23
# Identity:       6/23 (26.1%)
# Similarity:    10/23 (43.5%)
# Gaps:           0/23 ( 0.0%)
# Score: 22.0
# 
#
#=======================================

Binding_pep1       1 KALELAIDQASQDYNRANVLEKE     23
                     |.:|..:.:|:........|.||
M75_similar_p      1 KKVEADLAEANSKLQALEKLNKE     23


#---------------------------------------
#---------------------------------------
```

```
########################################
# Program: needle
# Commandline: needle
#    -auto
#    -stdout
#    -asequence emboss_needle-I20210917-140421-0869-93139689-p2m.asequence
#    -bsequence emboss_needle-I20210917-140421-0869-93139689-p2m.bsequence
#    -datafile EBLOSUM62
#    -gapopen 10.0
#    -gapextend 0.5
#    -endopen 10.0
#    -endextend 0.5
#    -aformat3 pair
#    -sprotein1
#    -sprotein2
# Align_format: pair
# Report_file: stdout
########################################

#=======================================
#
# Aligned_sequences: 2
# 1: Binding_pep2
# 2: M75_similar_p2
# Matrix: EBLOSUM62
# Gap_penalty: 10.0
# Extend_penalty: 0.5
#
# Length: 23
# Identity:      17/23 (73.9%)
# Similarity:    19/23 (82.6%)
# Gaps:           0/23 ( 0.0%)
# Score: 76.0
#
#
#=======================================

Binding_pep2       1 REAKKQVEKDLANLTAELDKVKE     23
                     |||||:||.|||.||||..|:||
M75_similar_p      1 REAKKKVEADLAALTAEHQKLKE     23


#---------------------------------------
#---------------------------------------
```

```
########################################
# Program: needle
# Commandline: needle
#    -auto
#    -stdout
#    -asequence emboss_needle-I20210917-140533-0491-60880675-p1m.asequence
#    -bsequence emboss_needle-I20210917-140533-0491-60880675-p1m.bsequence
#    -datafile EBLOSUM62
#    -gapopen 10.0
#    -gapextend 0.5
#    -endopen 10.0
#    -endextend 0.5
#    -aformat3 pair
#    -sprotein1
#    -sprotein2
# Align_format: pair
# Report_file: stdout
########################################

#=======================================
#
# Aligned_sequences: 2
# 1: Binding_pep1
# 2: M79_similar_p1
# Matrix: EBLOSUM62
# Gap_penalty: 10.0
# Extend_penalty: 0.5
#
# Length: 26
# Identity:      10/26 (38.5%)
# Similarity:    12/26 (46.2%)
# Gaps:           3/26 (11.5%)
# Score: 29.0
#
#
#=======================================

Binding_pep1       1 KALELAIDQASQDYN---RANVLEKE     23
                     ||.|:...||...:|   ..|.|.||
M79_similar_p      1 KAAEIKKPQADSAWNWPKEYNALLKE     26


#---------------------------------------
#---------------------------------------
```

```
########################################
# Program: needle
# Commandline: needle
#    -auto
#    -stdout
#    -asequence emboss_needle-I20210917-140643-0717-60803203-p2m.asequence
#    -bsequence emboss_needle-I20210917-140643-0717-60803203-p2m.bsequence
#    -datafile EBLOSUM62
#    -gapopen 10.0
#    -gapextend 0.5
#    -endopen 10.0
#    -endextend 0.5
#    -aformat3 pair
#    -sprotein1
#    -sprotein2
# Align_format: pair
# Report_file: stdout
########################################

#=======================================
#
# Aligned_sequences: 2
# 1: Binding_pep2
# 2: M79_similar_p2
# Matrix: EBLOSUM62
# Gap_penalty: 10.0
# Extend_penalty: 0.5
#
# Length: 23
# Identity:      17/23 (73.9%)
# Similarity:    19/23 (82.6%)
# Gaps:           0/23 ( 0.0%)
# Score: 76.0
#
#
#=======================================

Binding_pep2       1 REAKKQVEKDLANLTAELDKVKE     23
                     |||||:||.|||.||||..|:||
M79_similar_p      1 REAKKKVEADLAALTAEHQKLKE     23


#---------------------------------------
#---------------------------------------
```

```
########################################
# Program: needle
# Commandline: needle
#    -auto
#    -stdout
#    -asequence emboss_needle-I20210917-140742-0256-53359928-p2m.asequence
#    -bsequence emboss_needle-I20210917-140742-0256-53359928-p2m.bsequence
#    -datafile EBLOSUM62
#    -gapopen 10.0
#    -gapextend 0.5
#    -endopen 10.0
#    -endextend 0.5
#    -aformat3 pair
#    -sprotein1
#    -sprotein2
# Align_format: pair
# Report_file: stdout
########################################

#=======================================
#
# Aligned_sequences: 2
# 1: Binding_pep1
# 2: M81_similar_p1
# Matrix: EBLOSUM62
# Gap_penalty: 10.0
# Extend_penalty: 0.5
#
# Length: 23
# Identity:       6/23 (26.1%)
# Similarity:    10/23 (43.5%)
# Gaps:           0/23 ( 0.0%)
# Score: 22.0
#
#
#=======================================

Binding_pep1       1 KALELAIDQASQDYNRANVLEKE     23
                     |.:|..:.:|:........|.||
M81_similar_p      1 KKVEADLAEANSKLQALEKLNKE     23


#---------------------------------------
#---------------------------------------
```

```
########################################
# Program: needle
# Commandline: needle
#    -auto
#    -stdout
#    -asequence emboss_needle-I20210917-140900-0106-40553270-p2m.asequence
#    -bsequence emboss_needle-I20210917-140900-0106-40553270-p2m.bsequence
#    -datafile EBLOSUM62
#    -gapopen 10.0
#    -gapextend 0.5
#    -endopen 10.0
#    -endextend 0.5
#    -aformat3 pair
#    -sprotein1
#    -sprotein2
# Align_format: pair
# Report_file: stdout
########################################

#=======================================
#
# Aligned_sequences: 2
# 1: Binding_pep2
# 2: M81_similar_p2
# Matrix: EBLOSUM62
# Gap_penalty: 10.0
# Extend_penalty: 0.5
#
# Length: 23
# Identity:      17/23 (73.9%)
# Similarity:    19/23 (82.6%)
# Gaps:           0/23 ( 0.0%)
# Score: 76.0
#
#
#=======================================

Binding_pep2       1 REAKKQVEKDLANLTAELDKVKE     23
                     |||||:||.|||.||||..|:||
M81_similar_p      1 REAKKKVEADLAALTAEHQKLKE     23


#---------------------------------------
#---------------------------------------
```

```
########################################
# Program: needle
# Commandline: needle
#    -auto
#    -stdout
#    -asequence emboss_needle-I20210917-140947-0333-77671278-p1m.asequence
#    -bsequence emboss_needle-I20210917-140947-0333-77671278-p1m.bsequence
#    -datafile EBLOSUM62
#    -gapopen 10.0
#    -gapextend 0.5
#    -endopen 10.0
#    -endextend 0.5
#    -aformat3 pair
#    -sprotein1
#    -sprotein2
# Align_format: pair
# Report_file: stdout
########################################

#=======================================
#
# Aligned_sequences: 2
# 1: Binding_pep1
# 2: M87_similar_p1
# Matrix: EBLOSUM62
# Gap_penalty: 10.0
# Extend_penalty: 0.5
#
# Length: 34
# Identity:       9/34 (26.5%)
# Similarity:    15/34 (44.1%)
# Gaps:          11/34 (32.4%)
# Score: 32.0
#
#
#=======================================

Binding_pep1       1 KALELAIDQASQDYNRA-----------NVLEKE     23
                     ..||..:::|.:||:..           :.||||
M87_similar_p      1 SKLEKQLEEAQKDYSEIEGKLEQFWHDYDKLEKE     34


#---------------------------------------
#---------------------------------------
```

```
########################################
# Program: needle
# Commandline: needle
#    -auto
#    -stdout
#    -asequence emboss_needle-I20210917-141049-0262-1155708-p2m.asequence
#    -bsequence emboss_needle-I20210917-141049-0262-1155708-p2m.bsequence
#    -datafile EBLOSUM62
#    -gapopen 10.0
#    -gapextend 0.5
#    -endopen 10.0
#    -endextend 0.5
#    -aformat3 pair
#    -sprotein1
#    -sprotein2
# Align_format: pair
# Report_file: stdout
########################################

#=======================================
#
# Aligned_sequences: 2
# 1: Binding_pep2
# 2: M87_similar_p2
# Matrix: EBLOSUM62
# Gap_penalty: 10.0
# Extend_penalty: 0.5
#
# Length: 23
# Identity:      10/23 (43.5%)
# Similarity:    13/23 (56.5%)
# Gaps:           7/23 (30.4%)
# Score: 35.0
# 
#
#=======================================

Binding_pep2       1 REAKKQVEKDLANLTAELDKVKE     23
                     |.|||::|       ||..|:||
M87_similar_p      1 RAAKKELE-------AEHQKLKE     16


#---------------------------------------
#---------------------------------------
```

```
########################################
# Program: needle
# Commandline: needle
#    -auto
#    -stdout
#    -asequence emboss_needle-I20210917-141135-0412-59042471-p2m.asequence
#    -bsequence emboss_needle-I20210917-141135-0412-59042471-p2m.bsequence
#    -datafile EBLOSUM62
#    -gapopen 10.0
#    -gapextend 0.5
#    -endopen 10.0
#    -endextend 0.5
#    -aformat3 pair
#    -sprotein1
#    -sprotein2
# Align_format: pair
# Report_file: stdout
########################################

#=======================================
#
# Aligned_sequences: 2
# 1: Binding_pep1
# 2: M89_similar_p1
# Matrix: EBLOSUM62
# Gap_penalty: 10.0
# Extend_penalty: 0.5
#
# Length: 23
# Identity:       7/23 (30.4%)
# Similarity:    12/23 (52.2%)
# Gaps:           2/23 ( 8.7%)
# Score: 20.5
#
#
#=======================================

Binding_pep1       1 KALELAIDQASQDYNRANVLEKE     23
                     ||.|.:.:....:.|.|.  |::|
M89_similar_p      1 KAKEKSSENVERHYLRK--LDQE     21


#---------------------------------------
#---------------------------------------
```

```
########################################
# Program: needle
# Commandline: needle
#    -auto
#    -stdout
#    -asequence emboss_needle-I20210917-141215-0123-81889031-p1m.asequence
#    -bsequence emboss_needle-I20210917-141215-0123-81889031-p1m.bsequence
#    -datafile EBLOSUM62
#    -gapopen 10.0
#    -gapextend 0.5
#    -endopen 10.0
#    -endextend 0.5
#    -aformat3 pair
#    -sprotein1
#    -sprotein2
# Align_format: pair
# Report_file: stdout
########################################

#=======================================
#
# Aligned_sequences: 2
# 1: Binding_pep2
# 2: M89_similar_p2
# Matrix: EBLOSUM62
# Gap_penalty: 10.0
# Extend_penalty: 0.5
#
# Length: 23
# Identity:      17/23 (73.9%)
# Similarity:    19/23 (82.6%)
# Gaps:           0/23 ( 0.0%)
# Score: 76.0
#
#
#=======================================

Binding_pep2       1 REAKKQVEKDLANLTAELDKVKE     23
                     |||||:||.|||.||||..|:||
M89_similar_p      1 REAKKKVEADLAALTAEHQKLKE     23


#---------------------------------------
#---------------------------------------
```

```
########################################
# Program: needle
# Commandline: needle
#    -auto
#    -stdout
#    -asequence emboss_needle-I20210917-141302-0615-69440915-p1m.asequence
#    -bsequence emboss_needle-I20210917-141302-0615-69440915-p1m.bsequence
#    -datafile EBLOSUM62
#    -gapopen 10.0
#    -gapextend 0.5
#    -endopen 10.0
#    -endextend 0.5
#    -aformat3 pair
#    -sprotein1
#    -sprotein2
# Align_format: pair
# Report_file: stdout
########################################

#=======================================
#
# Aligned_sequences: 2
# 1: Binding_pep1
# 2: M179_similar_p1
# Matrix: EBLOSUM62
# Gap_penalty: 10.0
# Extend_penalty: 0.5
#
# Length: 23
# Identity:       7/23 (30.4%)
# Similarity:    12/23 (52.2%)
# Gaps:           0/23 ( 0.0%)
# Score: 29.0
#
#
#=======================================

Binding_pep1       1 KALELAIDQASQDYNRANVLEKE     23
                     |.:|.|:::|:........|.||
M179_similar_      1 KQVEKALEEANSKLAALEKLNKE     23


#---------------------------------------
#---------------------------------------
```

```
########################################
# Program: needle
# Commandline: needle
#    -auto
#    -stdout
#    -asequence emboss_needle-I20210917-141404-0386-78941509-p1m.asequence
#    -bsequence emboss_needle-I20210917-141404-0386-78941509-p1m.bsequence
#    -datafile EBLOSUM62
#    -gapopen 10.0
#    -gapextend 0.5
#    -endopen 10.0
#    -endextend 0.5
#    -aformat3 pair
#    -sprotein1
#    -sprotein2
# Align_format: pair
# Report_file: stdout
########################################

#=======================================
#
# Aligned_sequences: 2
# 1: Binding_pep2
# 2: M179_similar_p2
# Matrix: EBLOSUM62
# Gap_penalty: 10.0
# Extend_penalty: 0.5
#
# Length: 23
# Identity:      23/23 (100.0%)
# Similarity:    23/23 (100.0%)
# Gaps:           0/23 ( 0.0%)
# Score: 110.0
#
#
#=======================================

Binding_pep2       1 REAKKQVEKDLANLTAELDKVKE     23
                     |||||||||||||||||||||||
M179_similar_      1 REAKKQVEKDLANLTAELDKVKE     23


#---------------------------------------
#---------------------------------------
```

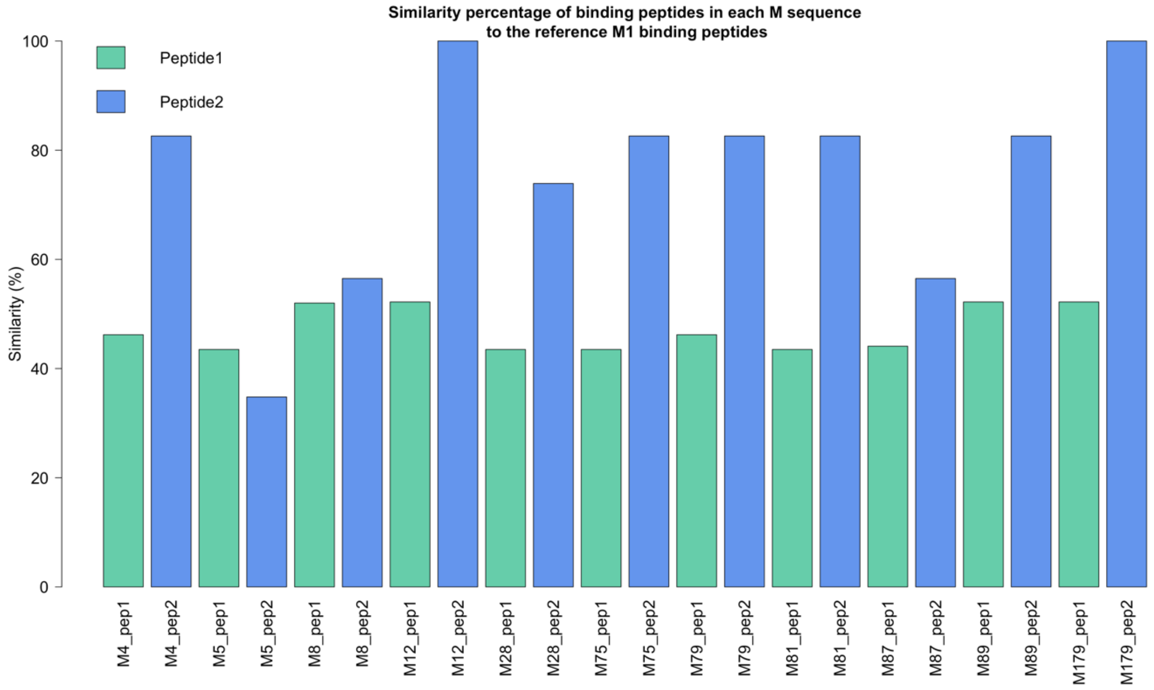


*Figure S2. Similarity percentage for binding peptides of M1 against most similar peptide in each M sequence*

**Sequence similarity with M79 partial sequence as the reference**

Here, the partial sequence of M79 as detected from sequencing data, is pairwise aligned against the rest of M proteins in this study. For those, M proteins that the full-length sequence is detected, we truncated the sequence to obtain a valid similarity score.

```
#########################################
# Program: needle
# Rundate: Wed  6 Oct 2021 09:49:10
# Commandline: needle
#    -auto
#    -stdout
#    -asequence emboss_needle-I20211006-094725-0867-6737632-p1m.asequence
#    -bsequence emboss_needle-I20211006-094725-0867-6737632-p1m.bsequence
#    -datafile EBLOSUM62
#    -gapopen 10.0
#    -gapextend 0.5
#    -endopen 10.0
#    -endextend 0.5
#    -aformat3 pair
#    -sprotein1
#    -sprotein2
# Align_format: pair
# Report_file: stdout
########################################

#=======================================
#
# Aligned_sequences: 2
# 1: Ref_M79_sequence
# 2: M1_sequence
# Matrix: EBLOSUM62
# Gap_penalty: 10.0
# Extend_penalty: 0.5
#
# Length: 295
# Identity:     125/295 (42.4%)
# Similarity:   157/295 (53.2%)
# Gaps:          75/295 (25.4%)
# Score: 461.0
#
#
#=======================================

Ref_M79_seque      1 MARKDTNKQYSLRKLKTGTASVAVAVAVLGAGFANQTEVKAAEIKKPQ--     48
                     ||:.:||:.||||||||||||||||:.||||||||||||||.....|:  
M1_sequence        1 MAKNNTNRHYSLRKLKTGTASVAVALTVLGAGFANQTEVKANGDGNPREV     50

Ref_M79_seque     49 -ADSAWNWPKEYNALLK-ENEELKVEREKYLSY-------ADDKEKDPQY     89
                      .|.|.|.|...|..|: ||::||...|..:..       |::.||..| 
M1_sequence       51 IEDLAANNPAIQNIRLRYENKDLKARLENAMEVAGRDFKRAEELEKAKQ-     99

Ref_M79_seque     90 RALMGENQDLRKREGQYQDKI-----------EELEKERKEKQERQE---    125
                      ||..:.:||..:..:.|...           :.||||.:||:|..|   
M1_sequence      100 -ALEDQRKDLETKLKELQQDYDLAKESTSWDRQRLEKELEEKKEALELAI    148

Ref_M79_seque    126 -QLERQYQ--IEADKHYQEQQK---------------------------K    145
                      |..|.|.  ...:|..:|::|                           :
M1_sequence      149 DQASRDYHRATALEKELEEKKALELAIDQASQDYNRANVLEKELETITR    198

Ref_M79_seque    146 HQQ------------------EQQQLEAEKQKLAKDKQISDASRQGLSRD    177
                     .|:                  |::||..||.||.::|||||||||.|.||
M1_sequence      199 EQEINRNLLGNAKLELDQLSSEKEQLTIEKAKLEEEKQISDASRQSLRRD    248

Ref_M79_seque    178 LEASREAKKKVEADLAALTAEHQKLKEDKQISDASRQGLSRDLEA    222
                     |:|||||||:||.|||.||||..|:||||||||||||||.|||:|
M1_sequence      249 LDASREAKKQVEKDLANLTAELDKVKEDKQISDASRQGLRRDLDA    293


#---------------------------------------
#---------------------------------------
```

```
########################################
# Program: needle
# Rundate: Wed  6 Oct 2021 09:51:02
# Commandline: needle
#    -auto
#    -stdout
#    -asequence emboss_needle-I20211006-095058-0469-54470083-p2m.asequence
#    -bsequence emboss_needle-I20211006-095058-0469-54470083-p2m.bsequence
#    -datafile EBLOSUM62
#    -gapopen 10.0
#    -gapextend 0.5
#    -endopen 10.0
#    -endextend 0.5
#    -aformat3 pair
#    -sprotein1
#    -sprotein2
# Align_format: pair
# Report_file: stdout
########################################

#=======================================
#
# Aligned_sequences: 2
# 1: Ref_M79_sequence
# 2: M4_sequence
# Matrix: EBLOSUM62
# Gap_penalty: 10.0
# Extend_penalty: 0.5
#
# Length: 222
# Identity:     222/222 (100.0%)
# Similarity:   222/222 (100.0%)
# Gaps:           0/222 ( 0.0%)
# Score: 1113.0
#
#
#=======================================

Ref_M79_seque      1 MARKDTNKQYSLRKLKTGTASVAVAVAVLGAGFANQTEVKAAEIKKPQAD     50
                     ||||||||||||||||||||||||||||||||||||||||||||||||||
M4_sequence        1 MARKDTNKQYSLRKLKTGTASVAVAVAVLGAGFANQTEVKAAEIKKPQAD     50

Ref_M79_seque     51 SAWNWPKEYNALLKENEELKVEREKYLSYADDKEKDPQYRALMGENQDLR    100
                     ||||||||||||||||||||||||||||||||||||||||||||||||||
M4_sequence       51 SAWNWPKEYNALLKENEELKVEREKYLSYADDKEKDPQYRALMGENQDLR    100

Ref_M79_seque    101 KREGQYQDKIEELEKERKEKQERQEQLERQYQIEADKHYQEQQKKHQQEQ    150
                     ||||||||||||||||||||||||||||||||||||||||||||||||||
M4_sequence      101 KREGQYQDKIEELEKERKEKQERQEQLERQYQIEADKHYQEQQKKHQQEQ    150

Ref_M79_seque    151 QQLEAEKQKLAKDKQISDASRQGLSRDLEASREAKKKVEADLAALTAEHQ    200
                     ||||||||||||||||||||||||||||||||||||||||||||||||||
M4_sequence      151 QQLEAEKQKLAKDKQISDASRQGLSRDLEASREAKKKVEADLAALTAEHQ    200

Ref_M79_seque    201 KLKEDKQISDASRQGLSRDLEA    222
                     ||||||||||||||||||||||
M4_sequence      201 KLKEDKQISDASRQGLSRDLEA    222


#---------------------------------------
#---------------------------------------
```

```
########################################
# Program: needle
# Rundate: Wed  6 Oct 2021 09:44:41
# Commandline: needle
#    -auto
#    -stdout
#    -asequence emboss_needle-I20211006-095144-0370-41504014-p1m.asequence
#    -bsequence emboss_needle-I20211006-095144-0370-41504014-p1m.bsequence
#    -datafile EBLOSUM62
#    -gapopen 10.0
#    -gapextend 0.5
#    -endopen 10.0
#    -endextend 0.5
#    -aformat3 pair
#    -sprotein1
#    -sprotein2
# Align_format: pair
# Report_file: stdout
########################################

#=======================================
#
# Aligned_sequences: 2
# 1: Ref_M79_sequence
# 2: M5_sequence
# Matrix: EBLOSUM62
# Gap_penalty: 10.0
# Extend_penalty: 0.5
#
# Length: 231
# Identity:      70/231 (30.3%)
# Similarity:   112/231 (48.5%)
# Gaps:          46/231 (19.9%)
# Score: 219.5
# 
#
#=======================================

Ref_M79_seque      1 MARKDTNKQYSLRKLKTGTASVAVAVAVLGAGF-ANQTEVKAA----EIK     45
                     |||::|||.|||||||.||||||||::|||||. .|..||.||    .|.
M5_sequence        1 MARENTNKHYSLRKLKKGTASVAVALSVLGAGLVVNTNEVSAAVTRGTIN     50

Ref_M79_seque     46 KPQ--ADSAWNWPKEYNALLKENEELKVEREKYLSYADDKEKDPQYRALM     93
                     .||  .::...:..|.:.|..:||.||.|.|...:         :...|.
M5_sequence       51 DPQRAKEALDKYELENHDLKTKNEGLKTENEGLKT---------ENEGLK     91

Ref_M79_seque     94 GENQDLRKREGQYQDKIEELEKERKEKQERQEQLERQYQIEADKHYQEQ-    142
                     .||:.|:..:.:::.:.::|:::|.....::|.|||:.|   :..|..: 
M5_sequence       92 TENEGLKTEKKEHEAENDKLKQQRDTLSTQKETLEREVQ---NTQYNNET    138

Ref_M79_seque    143 -QKKHQQEQQQLEAEKQKLAKDKQISDASRQGLSRDLEASREAKKKVEAD    191
                      :.|:....::|...:|:||..:|.|..:.:.|:..||  :..|.|:   
M5_sequence      139 LKIKNGDLTKELNKTRQELANKQQESKENEKALNELLE--KTVKDKI---    183

Ref_M79_seque    192 LAALTAEHQKLKEDKQISDASRQGLSRDLEA    222
                          |:.|:.||...               
M5_sequence      184 -----AKEQENKETIG---------------    194


#---------------------------------------
#---------------------------------------
```

```
########################################
# Program: needle
# Rundate: Wed  6 Oct 2021 09:47:09
# Commandline: needle
#    -auto
#    -stdout
#    -asequence emboss_needle-I20211006-095238-0087-7711006-p1m.asequence
#    -bsequence emboss_needle-I20211006-095238-0087-7711006-p1m.bsequence
#    -datafile EBLOSUM62
#    -gapopen 10.0
#    -gapextend 0.5
#    -endopen 10.0
#    -endextend 0.5
#    -aformat3 pair
#    -sprotein1
#    -sprotein2
# Align_format: pair
# Report_file: stdout
########################################

#=======================================
#
# Aligned_sequences: 2
# 1: Ref_M79_sequence
# 2: M8_sequence
# Matrix: EBLOSUM62
# Gap_penalty: 10.0
# Extend_penalty: 0.5
#
# Length: 256
# Identity:     149/256 (58.2%)
# Similarity:   170/256 (66.4%)
# Gaps:          52/256 (20.3%)
# Score: 641.0
#
#
#=======================================

Ref_M79_seque      1 MARKDTNKQYSLRKLKTGTASVAVAVAVLGAGFANQTEVKAAEIKKPQAD     50
                     |||||||||||||||||||||||||||||||||||||||||...|.....
M8_sequence        1 MARKDTNKQYSLRKLKTGTASVAVAVAVLGAGFANQTEVKAESPKSHSIS     50

Ref_M79_seque     51 SAWNWPKEYNALLKENEELKVEREKYLSYADD-KEKDPQYRALMGENQDL     99
                     :......|.|.|::||.:||.:..:.|...|: :||||||||||||||||
M8_sequence       51 NNEQLINELNDLIEENNDLKDKLARNLDLLDNTREKDPQYRALMGENQDL    100

Ref_M79_seque    100 RKREGQYQDKIEELE----------------------KERKEKQERQ---    124
                     |::||:|||||::||                      :|.||:||||
M8_sequence      101 REKEGKYQDKIKKLEEKEKNLEKKSEDVERHYLKKLDQEHKEQQERQKNL    150

Ref_M79_seque    125 EQLERQYQIEADKHYQEQQKKHQQEQQQLEAEKQKLAKDKQISDASRQGL    174
                     |:||||.|.|.||.||||    .|:|||||.|       ||||:|||:.|
M8_sequence      151 EELERQSQREIDKRYQEQ----LQKQQQLETE-------KQISEASRKSL    189

Ref_M79_seque    175 SRDLEASREAKKKVEADLAALTAEHQKLKEDKQISDASRQGLSRDLEA--    222
                     ||||||||.|||.:|       ||||||||:|||||||||||||||||
M8_sequence      190 SRDLEASRAAKKDLE-------AEHQKLKEEKQISDASRQGLSRDLEASR    232

Ref_M79_seque    223 ------    222

M8_sequence      233 EAKKKV    238


#---------------------------------------
#---------------------------------------
```

```
########################################
# Program: needle
# Rundate: Wed  6 Oct 2021 09:53:19
# Commandline: needle
#    -auto
#    -stdout
#    -asequence emboss_needle-I20211006-095514-0934-92394355-p2m.asequence
#    -bsequence emboss_needle-I20211006-095514-0934-92394355-p2m.bsequence
#    -datafile EBLOSUM62
#    -gapopen 10.0
#    -gapextend 0.5
#    -endopen 10.0
#    -endextend 0.5
#    -aformat3 pair
#    -sprotein1
#    -sprotein2
# Align_format: pair
# Report_file: stdout
########################################

#=======================================
#
# Aligned_sequences: 2
# 1: Ref_M79_sequence
# 2: M12_sequence
# Matrix: EBLOSUM62
# Gap_penalty: 10.0
# Extend_penalty: 0.5
#
# Length: 323
# Identity:      89/323 (27.6%)
# Similarity:   134/323 (41.5%)
# Gaps:         117/323 (36.2%)
# Score: 270.5
# 
#
#=======================================

Ref_M79_seque      1 MARKDTNKQYSLRKLKTGTASVAVAVAVLGAGFANQTEVKAAEIKKPQAD     50
                     ||:..||:.||||||||||||||||:.|:|||......|:|
M12_sequence       1 MAKNTTNRHYSLRKLKTGTASVAVALTVVGAGLVAGQTVRA---------     41

Ref_M79_seque     51 SAWNWPKEYNALLKEN----------EELKVEREKYL-SYADDK------     83
                            :::.|:.|.          |.||...|.|| .|.|:|
M12_sequence      42 -------DHSDLVAEKQRLEDLGQKFERLKQRSELYLQQYYDNKSNGYKG     84

Ref_M79_seque     84 ------------EKDPQYRALMGE------------NQDLRKREGQYQDK    109
                                 :.:..|..|.||            |.||:.:..:.:..
M12_sequence      85 DWYVQQLKMLNRDLEQAYNELSGEAHKDALGKLGIDNADLKAKITELEKS    134

Ref_M79_seque    110 IEE----LEKERKEKQE-------------------RQEQLERQYQI---    133
                     :||    |.:.:||.:|                   :||..|::..|
M12_sequence     135 VEEKNDVLSQIKKELEEAEKDIQFGREVHAADLLRHKQEIAEKENVISKL    184

Ref_M79_seque    134 ------------EADKHYQEQQKKHQQEQQQLEAEKQKLAKDKQISDASR    171
                                 |.|::.|::::|....:|||...|:...||.:::....
M12_sequence     185 NGELQPLKQKVDETDRNLQQEKQKVLSLEQQLAVTKENAKKDFELAALGH    234

Ref_M79_seque    172 QGLSRDLEA------SREAKKKVEADLAALTAEH----------------    199
                     |...::..|      |:.|..|.:.:||||..:|
M12_sequence     235 QLADKEYNAKIAELESKLADAKKDFELAALGHQHAHNEYQAKLAEKDGQI    284

Ref_M79_seque    200 QKLKEDKQISDASRQGLSRDLEA    222
                     ::|:|.|||.||||:|.:|||||
M12_sequence     285 KQLEEQKQILDASRKGTARDLEA    307


#---------------------------------------
#---------------------------------------
```

```
########################################
# Program: needle
# Rundate: Wed  6 Oct 2021 09:53:38
# Commandline: needle
#    -auto
#    -stdout
#    -asequence emboss_needle-I20211006-100101-0387-61592505-p1m.asequence
#    -bsequence emboss_needle-I20211006-100101-0387-61592505-p1m.bsequence
#    -datafile EBLOSUM62
#    -gapopen 10.0
#    -gapextend 0.5
#    -endopen 10.0
#    -endextend 0.5
#    -aformat3 pair
#    -sprotein1
#    -sprotein2
# Align_format: pair
# Report_file: stdout
########################################

#=======================================
#
# Aligned_sequences: 2
# 1: Ref_M79_sequence
# 2: M28_sequence
# Matrix: EBLOSUM62
# Gap_penalty: 10.0
# Extend_penalty: 0.5
#
# Length: 250
# Identity:     150/250 (60.0%)
# Similarity:   173/250 (69.2%)
# Gaps:          49/250 (19.6%)
# Score: 642.0
#
#
#=======================================

Ref_M79_seque      1 MARKDTNKQYSLRKLKTGTASVAVAVAVLGAGFANQTEVKAAEIKKPQAD     50
                     |||||||||||||||||||||||||||||||||||||||||||..|....
M28_sequence       1 MARKDTNKQYSLRKLKTGTASVAVAVAVLGAGFANQTEVKAAESPKSTET     50

Ref_M79_seque     51 SAWNWPK---EYNALLKENEELKVEREKYLSYADDKEKDPQYRALMGENQ     97
                     ||....|   .||.||.|:|:|   |::|.:..|.||::|:|:||.||||
M28_sequence      51 SANGADKLADAYNTLLTEHEKL---RDEYYTLIDAKEEEPRYKALRGENQ     97

Ref_M79_seque     98 DLRKREGQYQDKIEELE----------------------KERKEKQERQ-    124
                     |||::||:|||||::||                      :|.||::||| 
M28_sequence      98 DLREKEGKYQDKIKKLEEKEKNLEKKSEDVERHYLKKLDQEHKEQEERQK    147

Ref_M79_seque    125 --EQLERQYQIEADKHYQEQQKKHQQEQQQLEAEKQKLAKDKQISDASRQ    172
                       |:||||.|.|.||.||||    .|:|||||.|       ||||:|||:
M28_sequence     148 NLEELERQSQREIDKRYQEQ----LQKQQQLETE-------KQISEASRK    186

Ref_M79_seque    173 GLSRDLEASREAKKKVEADLAALTAEHQKLKEDKQISDASRQGLSRDLEA    222
                     .|||||||||.|||.:|       ||||||||:|||||||||||||||||
M28_sequence     187 SLSRDLEASRAAKKDLE-------AEHQKLKEEKQISDASRQGLSRDLEA    229


#---------------------------------------
#---------------------------------------
```

```
########################################
# Program: needle
# Rundate: Wed  6 Oct 2021 09:58:18
# Commandline: needle
#    -auto
#    -stdout
#    -asequence emboss_needle-I20211006-100349-0215-55773762-p1m.asequence
#    -bsequence emboss_needle-I20211006-100349-0215-55773762-p1m.bsequence
#    -datafile EBLOSUM62
#    -gapopen 10.0
#    -gapextend 0.5
#    -endopen 10.0
#    -endextend 0.5
#    -aformat3 pair
#    -sprotein1
#    -sprotein2
# Align_format: pair
# Report_file: stdout
########################################

#=======================================
#
# Aligned_sequences: 2
# 1: Ref_M79_sequence
# 2: M75_sequence
# Matrix: EBLOSUM62
# Gap_penalty: 10.0
# Extend_penalty: 0.5
#
# Length: 224
# Identity:     112/224 (50.0%)
# Similarity:   143/224 (63.8%)
# Gaps:          10/224 ( 4.5%)
# Score: 450.0
#
#
#=======================================

Ref_M79_seque      1 MARKDTNKQYSLRKLKTGTASVAVAVAVLGAGFANQTEVKAAEIKKPQAD     50
                     |||||||||||||||||||||||||||||||||||||||||.|   .:..
M75_sequence       1 MARKDTNKQYSLRKLKTGTASVAVAVAVLGAGFANQTEVKAEE---ERTF     47

Ref_M79_seque     51 SAWNWPKEYNALLKENEELKVEREKYLSYADDKEKDPQYRALMGENQDLR    100
                     :...:...|.|...||:||:....:.|    ||....|.:....|.|: :
M75_sequence      48 TELPYEARYKAWKSENDELRENYRRTL----DKFNTEQGKTTRLEEQN-K     92

Ref_M79_seque    101 KREGQYQDKIEELEKERKEKQERQEQLERQYQIEAD--KHYQEQQKKHQQ    148
                     |...:.....|.|....:...::.:.|..:.:|.::  .:.::|..|...
M75_sequence      93 KLHSELASVTETLTSVTEADDKKIKDLTDRDKISSNLLGNAKDQINKLTT    142

Ref_M79_seque    149 EQQQLEAEKQKLAKDKQISDASRQGLSRDLEASREAKKKVEADLAALTAE    198
                     |:..|..:.:||.:|||||||||:.|||||||||.|||::||:...|..|
M75_sequence     143 EKDTLAEKAKKLEEDKQISDASRKSLSRDLEASRAAKKELEANHQKLETE    192

Ref_M79_seque    199 HQKLKEDKQISDASRQGLSRDLEA    222
                     |:||||:|||||||||||||||||
M75_sequence     193 HKKLKEEKQISDASRQGLSRDLEA    216


#---------------------------------------
#---------------------------------------
```

```
########################################
# Program: needle
# Rundate: Wed  6 Oct 2021 10:07:10
# Commandline: needle
#    -auto
#    -stdout
#    -asequence emboss_needle-I20211006-100706-0795-49043706-p2m.asequence
#    -bsequence emboss_needle-I20211006-100706-0795-49043706-p2m.bsequence
#    -datafile EBLOSUM62
#    -gapopen 10.0
#    -gapextend 0.5
#    -endopen 10.0
#    -endextend 0.5
#    -aformat3 pair
#    -sprotein1
#    -sprotein2
# Align_format: pair
# Report_file: stdout
########################################

#=======================================
#
# Aligned_sequences: 2
# 1: Ref_M79_sequence
# 2: M81_sequence
# Matrix: EBLOSUM62
# Gap_penalty: 10.0
# Extend_penalty: 0.5
#
# Length: 223
# Identity:     124/223 (55.6%)
# Similarity:   148/223 (66.4%)
# Gaps:          18/223 ( 8.1%)
# Score: 509.5
#
#
#=======================================

Ref_M79_seque      1 MARKDTNKQYSLRKLKTGTASVAVAVAVLGAGFANQTEVKAAEIKKPQAD     50
                     |.|||||||||||||||||||||||||||||||||||||||       |.
M81_sequence       1 MVRKDTNKQYSLRKLKTGTASVAVAVAVLGAGFANQTEVKA-------AG     43

Ref_M79_seque     51 SAWNWPK-EYNALLKENEELKVEREKYLSYADDKEKDPQYRALMGENQDL     99
                     |..|.|| :||||.:|||:|:....||::..:.:|  .|...|..:|:.|
M81_sequence      44 SEENVPKQQYNALWEENEDLRGRERKYIAKLEKEE--IQNGELNEKNRKL     91

Ref_M79_seque    100 RKREGQYQDKIEELEKERKEKQERQEQLERQYQIEADKHYQEQQKKHQQE    149
                     .......||.||:.::|.|.|       :|.|:... |..::|.|....|
M81_sequence      92 EADIADLQDVIEDNDQEIKRK-------DRMYEAFL-KQSKDQVKDLTAE    133

Ref_M79_seque    150 QQQLEAEKQKLAKDKQISDASRQGLSRDLEASREAKKKVEADLAALTAEH    199
                     :..|..:.:||.:|||||||||:.||||||.||.|||::||....|..||
M81_sequence     134 KDTLAEKAKKLEEDKQISDASRKSLSRDLEGSRAAKKELEAKHQKLETEH    183

Ref_M79_seque    200 QKLKEDKQISDASRQGLSRDLEA    222
                     |||||||||||||||||||||||
M81_sequence     184 QKLKEDKQISDASRQGLSRDLEA    206


#---------------------------------------
#---------------------------------------
```

```
########################################
# Program: needle
# Rundate: Wed  6 Oct 2021 10:10:16
# Commandline: needle
#    -auto
#    -stdout
#    -asequence emboss_needle-I20211006-101012-0859-31002285-p2m.asequence
#    -bsequence emboss_needle-I20211006-101012-0859-31002285-p2m.bsequence
#    -datafile EBLOSUM62
#    -gapopen 10.0
#    -gapextend 0.5
#    -endopen 10.0
#    -endextend 0.5
#    -aformat3 pair
#    -sprotein1
#    -sprotein2
# Align_format: pair
# Report_file: stdout
########################################

#=======================================
#
# Aligned_sequences: 2
# 1: Ref_M79_sequence
# 2: M87_sequence
# Matrix: EBLOSUM62
# Gap_penalty: 10.0
# Extend_penalty: 0.5
#
# Length: 246
# Identity:     131/246 (53.3%)
# Similarity:   159/246 (64.6%)
# Gaps:          42/246 (17.1%)
# Score: 516.5
# 
#
#=======================================

Ref_M79_seque      1 MARKDTNKQYSLRKLKTGTASVAVAVAVLGAGFANQTEVKA---AEIKKP     47
                     |||||||||||||||||||||||||||||||||||||||||   .|:...
M87_sequence       1 MARKDTNKQYSLRKLKTGTASVAVAVAVLGAGFANQTEVKAESPREVTNE     50

Ref_M79_seque     48 QADSAW------------NWPKEYNALLKENEELKVEREKYLSYADDKEK     85
                     .|.|.|            ...|:.....|:..|::.:.|::....|..||
M87_sequence      51 LAASVWKKKVEEAKEKASKLEKQLEEAQKDYSEIEGKLEQFWHDYDKLEK    100

Ref_M79_seque     86 D-PQYRALMGENQDLR-KREGQYQDKIEELEKE----RKEKQERQ---EQ    126
                     : .:|.:.:|:||:.| |.|.:|..|.:|..||    |:|::|||   |:
M87_sequence     101 ENKEYASQLGKNQEEREKLELEYLRKSDEEYKEHQQYRQEQEERQKNLEE    150

Ref_M79_seque    127 LERQYQIEADKHYQEQQKKHQQEQQQLEAEKQKLAKDKQISDASRQGLSR    176
                     ||||.:.|.||.||||    .|:|||||.|       ||||:|||:.|||
M87_sequence     151 LERQNKREIDKRYQEQ----LQKQQQLETE-------KQISEASRKSLSR    189

Ref_M79_seque    177 DLEASREAKKKVEADLAALTAEHQKLKEDKQISDASRQGLSRDLEA    222
                     ||||||.|||::|       ||||||||:||||||||:.|||||||
M87_sequence     190 DLEASRAAKKELE-------AEHQKLKEEKQISDASRKSLSRDLEA    228


#---------------------------------------
#---------------------------------------
```

```
########################################
# Program: needle
# Rundate: Wed  6 Oct 2021 10:12:06
# Commandline: needle
#    -auto
#    -stdout
#    -asequence emboss_needle-I20211006-101051-0273-48298929-p1m.asequence
#    -bsequence emboss_needle-I20211006-101051-0273-48298929-p1m.bsequence
#    -datafile EBLOSUM62
#    -gapopen 10.0
#    -gapextend 0.5
#    -endopen 10.0
#    -endextend 0.5
#    -aformat3 pair
#    -sprotein1
#    -sprotein2
# Align_format: pair
# Report_file: stdout
########################################

#=======================================
#
# Aligned_sequences: 2
# 1: Ref_M79_sequence
# 2: M89_sequence
# Matrix: EBLOSUM62
# Gap_penalty: 10.0
# Extend_penalty: 0.5
#
# Length: 222
# Identity:      91/222 (41.0%)
# Similarity:   124/222 (55.9%)
# Gaps:          61/222 (27.5%)
# Score: 364.0
#
#
#=======================================

Ref_M79_seque      1 MARKDTNKQYSLRKLKTGTASVAVAVAVLGAGFANQTEVKAAEIKKPQAD     50
                     |||||||||||||||||||||||||||||||||||||.|||      .:|
M89_sequence       1 MARKDTNKQYSLRKLKTGTASVAVAVAVLGAGFANQTTVKA------DSD     44

Ref_M79_seque     51 SAWNWPKEYNALLKENEELKVEREKYLSYADDKEKDPQYRALMGENQDLR    100
                     :.             |..:.|:         |.||:     |..:..||.
M89_sequence      45 NI-------------NRSVSVK---------DNEKE-----LHNKIADLE     67

Ref_M79_seque    101 KREGQYQDKIEELEKERKEKQERQEQLERQYQIEADKHYQEQQKKHQQEQ    150
                     :..|::.|||:||::|.|.|::..|.:||.|..:.|:.|:|||:: |:..
M89_sequence      68 EERGEHLDKIDELKEELKAKEKSSENVERHYLRKLDQEYKEQQER-QKNL    116

Ref_M79_seque    151 QQLEAEKQKLAKDKQISDASRQGLSRDLEASREAKKKVEADLAALTAEHQ    200
                     ::||.:.|                       ||.:|:.:..|    .:.|
M89_sequence     117 EELERQSQ-----------------------REVEKRYQEQL----QKQQ    139

Ref_M79_seque    201 KLKEDKQISDASRQGLSRDLEA    222
                     :|:.:||||:|||:.|||||||
M89_sequence     140 QLETEKQISEASRKSLSRDLEA    161


#---------------------------------------
#---------------------------------------
```

```
########################################
# Program: needle
# Rundate: Wed  6 Oct 2021 10:13:16
# Commandline: needle
#    -auto
#    -stdout
#    -asequence emboss_needle-I20211006-101315-0196-46816129-p2m.asequence
#    -bsequence emboss_needle-I20211006-101315-0196-46816129-p2m.bsequence
#    -datafile EBLOSUM62
#    -gapopen 10.0
#    -gapextend 0.5
#    -endopen 10.0
#    -endextend 0.5
#    -aformat3 pair
#    -sprotein1
#    -sprotein2
# Align_format: pair
# Report_file: stdout
########################################

#=======================================
#
# Aligned_sequences: 2
# 1: Ref_M79_sequence
# 2: M179_sequence
# Matrix: EBLOSUM62
# Gap_penalty: 10.0
# Extend_penalty: 0.5
#
# Length: 289
# Identity:     118/289 (40.8%)
# Similarity:   154/289 (53.3%)
# Gaps:          72/289 (24.9%)
# Score: 447.0
# 
#
#=======================================

Ref_M79_seque      1 MARKDTNKQYSLRKLKTGTASVAVAVAVLGAGFA-NQTEVKAAEI-----     44
                     |.|||||:.||||||||||||||||::|||||.| |||||.|..:
M179_sequence      1 MVRKDTNRHYSLRKLKTGTASVAVALSVLGAGLAVNQTEVSAKSVTRSTA     50

Ref_M79_seque     45 ----KKPQADSAW-----NWPKEYNALLKENEEL----------------     69
                         |..||.:.:     ...:|.|||...|:||
M179_sequence     51 QDPDKSRQAITEYEVENHKLTQEKNALTNRNQELTDENGELKTANEALRQ    100

Ref_M79_seque     70 --------KVEREKYLSYADDKEKDPQYRALMGENQDLRKREGQYQDKIE    111
                             :|:.||.:     :||:...:.|..||.:|:...|....|::
M179_sequence    101 RGDTLFNQRVKLEKQV-----QEKEHNNKTLKIENGELKTENGDLTKKLD    145

Ref_M79_seque    112 ELEKERKEKQERQEQLERQY-----QIEADKHYQEQQK------------    144
                     |..:|...||:..::.|:..     :...||..:||:.
M179_sequence    146 ETRQELANKQQESKENEKTLNELLEKTVKDKIAKEQENKETIGTLKKLLD    195

Ref_M79_seque    145 --------KHQQEQQQLEAEKQKLAKDKQ---ISDASRQGLSRDLEASRE    183
                             |.|:.:|...|.||:|||.::   |||||||||.|||.||||
M179_sequence    196 ETVKDKIAKEQKSKQDFGALKQELAKKEEQNKISDASRQGLRRDLNASRE    245

Ref_M79_seque    184 AKKKVEADLAALTAEHQKLKEDKQISDASRQGLSRDLEA    222
                     |||:||.|||.||||..|:||:||:||||||||.|||:|
M179_sequence    246 AKKQVEKDLANLTAELDKVKEEKQVSDASRQGLRRDLDA    284


#---------------------------------------
#---------------------------------------
```

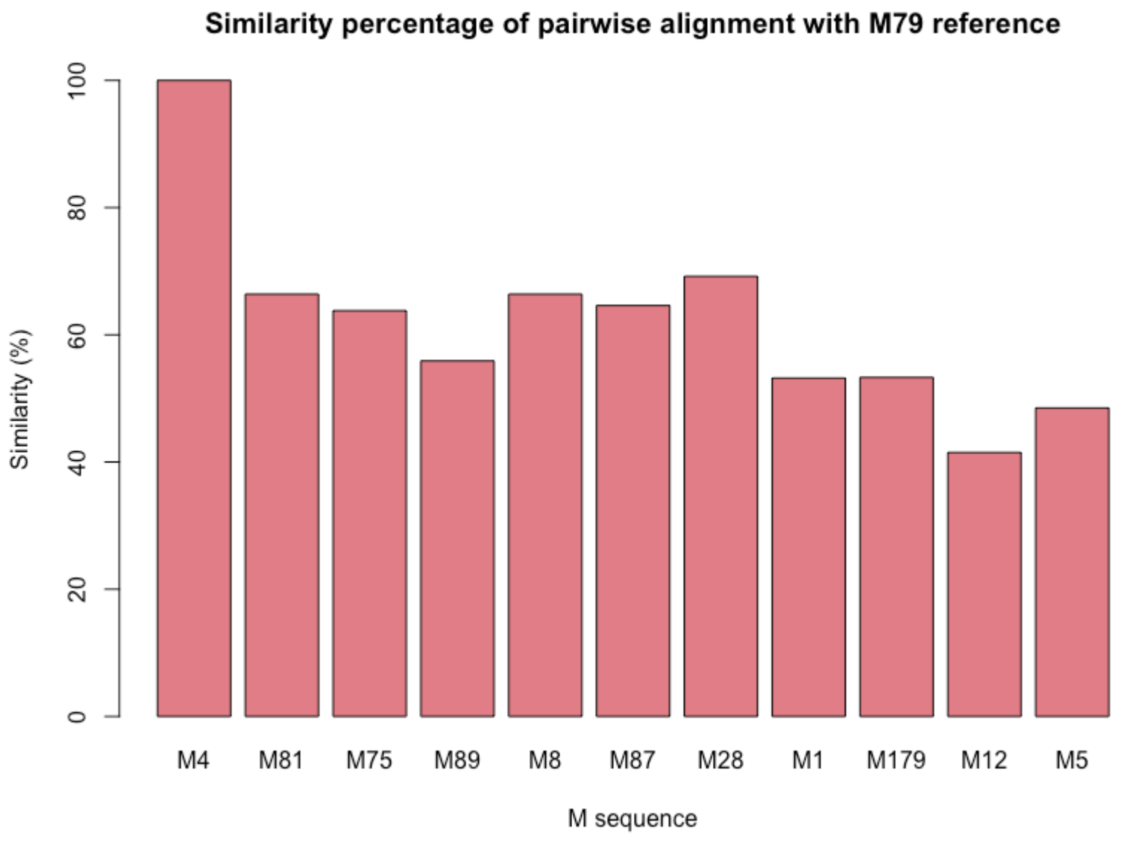


*Figure S3. Similarity percentage for truncated M sequences against M79 partial reference sequence*
